# Genetic diversity loss in the Anthropocene

**DOI:** 10.1101/2021.10.13.464000

**Authors:** Moises Exposito-Alonso, Tom R. Booker, Lucas Czech, Tadashi Fukami, Lauren Gillespie, Shannon Hateley, Christopher C. Kyriazis, Patricia L. M. Lang, Laura Leventhal, David Nogues-Bravo, Veronica Pagowski, Megan Ruffley, Jeffrey P. Spence, Sebastian E. Toro Arana, Clemens L. Weiß, Erin Zess

## Abstract

More species than ever before are at risk of extinction due to anthropogenic habitat loss and climate change. But even species that are not threatened have seen reductions in their populations and geographic ranges, likely impacting their genetic diversity. Although preserving genetic diversity is key to maintaining adaptability of species, we lack predictive tools and global estimates of genetic diversity loss across ecosystems. By bridging theories of biodiversity and population genetics, we introduce a mathematical framework to understand the loss of naturally occurring DNA mutations within decreasing habitat within a species. Analysing genome-wide variation data of 10,095 geo-referenced individuals from 20 plant and animal species, we show that genome-wide diversity follows a power law with geographic area (the mutations-area relationship), which can predict genetic diversity loss in spatial computer simulations of local population extinctions. Given pre-21^st^ century values of ecosystem transformations, we estimate that over 10% of genetic diversity may already be lost, surpassing the United Nations targets for genetic preservation. These estimated losses could rapidly accelerate with advancing climate change and habitat destruction, highlighting the need for forecasting tools that facilitate implementation of policies to protect genetic resources globally.

---

Anthropogenic habitat loss and climate change (*1, 2*) have led to the extinction of hundreds of species over the last centuries (*1, 2*) and approximately one million more species (25% of all known species) are at risk of extinction (*3*). It has been estimated that an even larger fraction—at least 47%—of plant and animal species have lost part of their geographic range in response to the last centuries of anthropogenic activities (*4, 5*). Though this loss might seem inconsequential compared to losing an entire species, this range contraction reduces genetic diversity, which dictates species’ ability to adapt to new environmental conditions (*6–8*). The loss of geographic range can spiral into a feedback loop where diversity loss further increases the risk of species extinction (*9, 10*).

Although genetic diversity is a key dimension of biodiversity (*11*), it has been overlooked in international conservation initiatives. Only in 2021 did the United Nations’ Convention of Biological Diversity propose to preserve at least 90% of all species’ genetic diversity (*12, 13*). Although analyses of genetic markers in animal populations sampled over time with the aim of quantifying recent genetic change are emerging (*14, 15*) and simulation studies with species distribution models or sensitivity analyses suggest within-species range variation may be strongly impacted (*5, 16, 17*), theory and scalable approaches to estimate genome-wide diversity loss across species do not yet exist, impairing prioritization and evaluation of conservation targets. Here, we introduce a framework to estimate global genetic diversity loss by bridging biodiversity theory with population genetics, and by combining data on global ecosystem transformations with newly available genomic datasets.

The first studies that predicted biodiversity reductions in response to habitat loss and climate change in the 1990s and the 2000s projected species extinctions using the relationship of biodiversity with geographic area—termed the species-area relationship (SAR) (*18*) (see **Supplementary Materials [SM] I** for a comparison of mathematical models for predicting biodiversity). In this framework, ecosystems with a larger area (*A*) harbour a larger number of species (*S*) resulting from a balance of limited dispersal, habitat heterogeneity, and colonisation-extinction-speciation dynamics. The more a study area is extended, the more species are found. The SAR has been empirically shown to follow a power law, *S = A^z^*. It scales consistently across continents and ecosystems (*19*), with a higher *z* characterising more speciose and spatially structured ecosystems. Given estimates of decreasing ecosystem areas over time (*A_t-1_ > A_t_*), Thomas et al. (*20*) proposed rough estimates of the percentage of species extinctions in the 21^st^ century ranging from 15 to 37% (**SM I.3**). Though this may be an oversimplification, SAR has become a common tool for policy groups including the Intergovernmental Science-Policy Platform on Biodiversity and Ecosystem Services (IPBES) (*3*).

As species richness is for to ecosystems’ biodiversity, within-species variation can be quantitatively described by the richness of genetic mutations within a species, defined here as DNA nucleotide variants appearing in individuals of a species. Although population genetics theory has long established that larger populations have higher genetic diversity (*21*), and it is known that geographic isolation between populations within the same species results in geographically separated accumulation of different mutations, there have been no attempts to describe the extent of genetic diversity loss driven by species’ geographic range reduction using an analogous “mutations-area relationship” (MAR).

We suspected that such a mutations-area relationship must exist given that another general assumption is shared with species studies, namely that when mutations appear they are first in only one individual, and they typically remain at low frequency in a population, though a few prevail to high frequency through stochastic genetic drift and natural selection (*22*). This principle of “commonness of rarity” is well-known for species (i.e. most species in an ecosystem are rare while only a few are common) and, together with limited spatial dispersal of species and communities, is a key statistical condition that led to the power-law SAR.

To examine the expectation of a power-law MAR, we begin quantifying the rarity of mutations using millions of biallelic genetic variants of the *Arabidopsis thaliana* 1001 genomes dataset (**Fig. 1A**) (*23*) by fitting several common models of species abundances (*24*) to the distribution of mutation frequencies (*q*), termed the Site Frequency Spectrum in population genetics (**Fig. 1B, SM II.1**). The canonical L-shaped probability distribution (*1/q*) of this spectrum—which is expected under population-equilibrium and the absence of natural selection processes—fit this data well (Fig. 1B), although the more parameter rich Preston’s species abundance log-normal model achieved the best AIC value (**Fig. 1B, SM III.1, Table S3, Table S10**). Despite the small differences in fit, these models all showcase the similarities of abundance distributions of mutations within species and species within ecosystems, suggesting that they may behave similarly in their relationship to geographic area (*22, 24*).

**Fig. 1.**
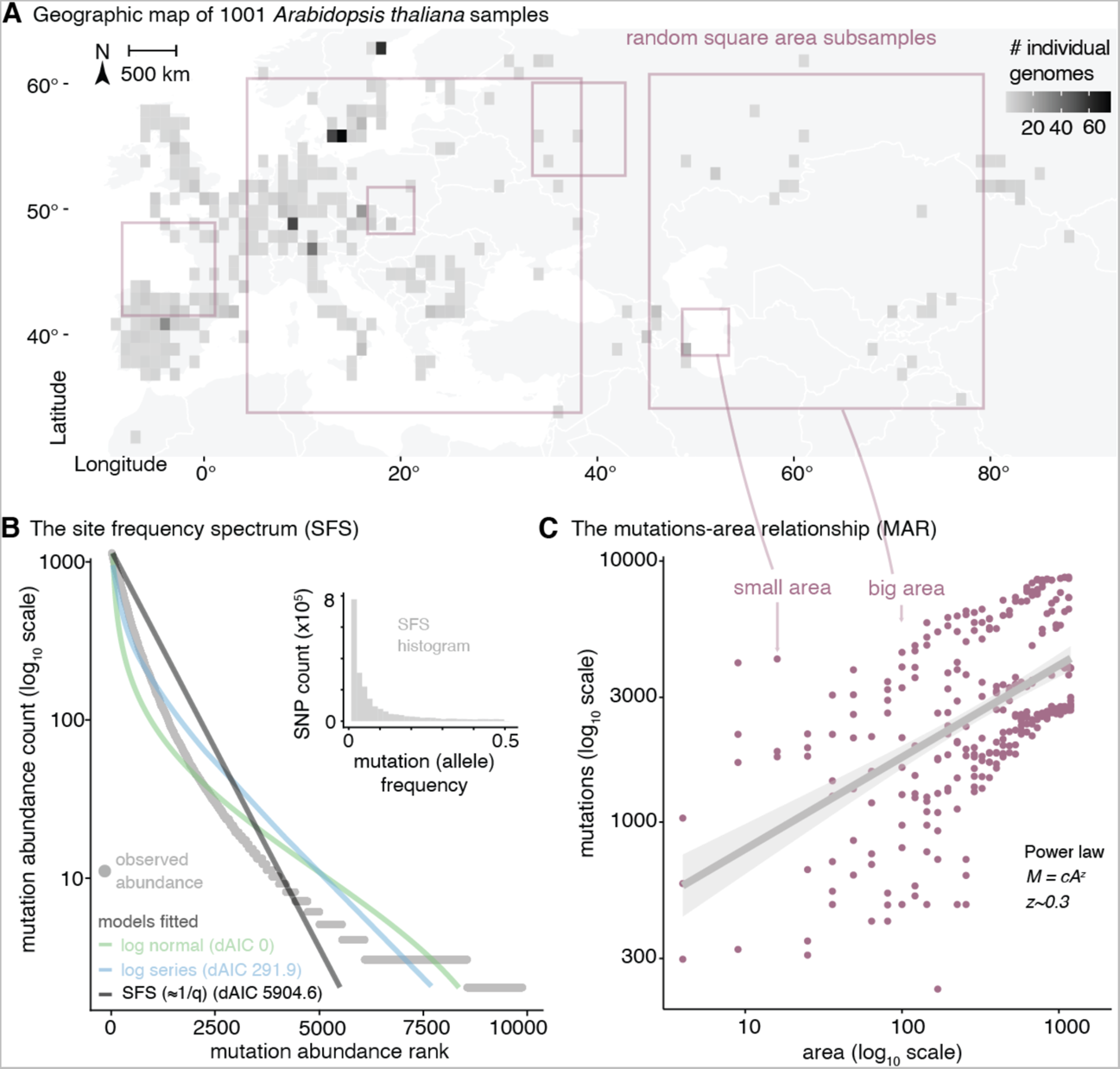
Mutations across populations follow a log-normal abundance distribution and a power law with species range area. (**A**) Density of individuals projected in a 1 x 1 degree latitude/longitude map of Europe and exemplary subsample areas of different sizes. (**B**) Distribution of mutation (SNPs) frequencies in 1,001 *Arabidopsis thaliana* plants using a site frequency spectrum histogram (grey inset) and a Whittaker’s rank abundance curve plot, and the fitted models of common species abundance functions in *A. thaliana* using a dataset random sample of 10,000 mutations also used in (C). The AIC fit of the three models is indicated with respect to the top model, log-normal. (**C**) The mutations-area relationship (MAR) in log-log space built from 10 random subsamples of different areas of increasing size within *A. thaliana*’s geographic range along with the number of mutations discovered for each area subset.

To quantify how genetic diversity within a species increases with geographic area, we constructed the MAR by subsampling different regions of different sizes of *Arabidopsis thaliana*’s native range using over one thousand geo-referenced genomes (**Fig. 1A, C**). As a metric of genetic diversity, we modelled the number of mutations (*M*) in space (number of segregating sites) consistent with the species-centric approach of SAR, which uses species richness as the metric of biodiversity (**SM II.2**). The MAR also followed the power law relationship *M = cA^z^* with a scaling value *z_MAR_* = 0.324 (CI95% = 0.238–0.41) (**Fig. 1C**). Naturally, subsamples of larger areas may also contain more individuals, and therefore should also have more mutations. But the observed power law relationship goes beyond what is expected from the increase of number of samples in an area (which only accounts for increases of *M* ≈ *log(A),* see theoretical derivation **SM II.3**). The remainder may be attributed to population genetic drift and spatial natural selection causing structuring of genetic diversity across populations. The discovered power law scaling appears robust to different methods of area quantification, the effects of non-random spatial patterns, random area sampling, fully nested outward or inward sampling (*19*), raster area calculations, raster grid resolution (~10–1,000 km side cell size), and is adjusted for limited sample sizes (**SM II.3.2, III.3, Fig. S14–18, Tables S7–9**).

We then wondered whether MAR can predict the loss of genetic diversity due to species’ range contractions. We explored several scenarios of range contraction in *A. thaliana* by removing *in silico* grid cells in a map representing populations that are lost (**Fig. 2B**). Our simulations included random local population extinction as if deforestation was scattered across large continents, radial expansion of an extinction front due to intense localised mortality, or local extinction in the warmest regions within a species range (*4, 25*), among others (**SM III.4**). The MAR-based predictions of genetic loss, using *1-(1-A_t_ / A_t-1_)^z^* and assuming *z* = 0.3, conservatively followed the simulated local loss in *A. thaliana* (pseudo-*R^2^* = 0.87, taking all simulations together) (**SM II.4, III.4)**.

**Fig. 2.**
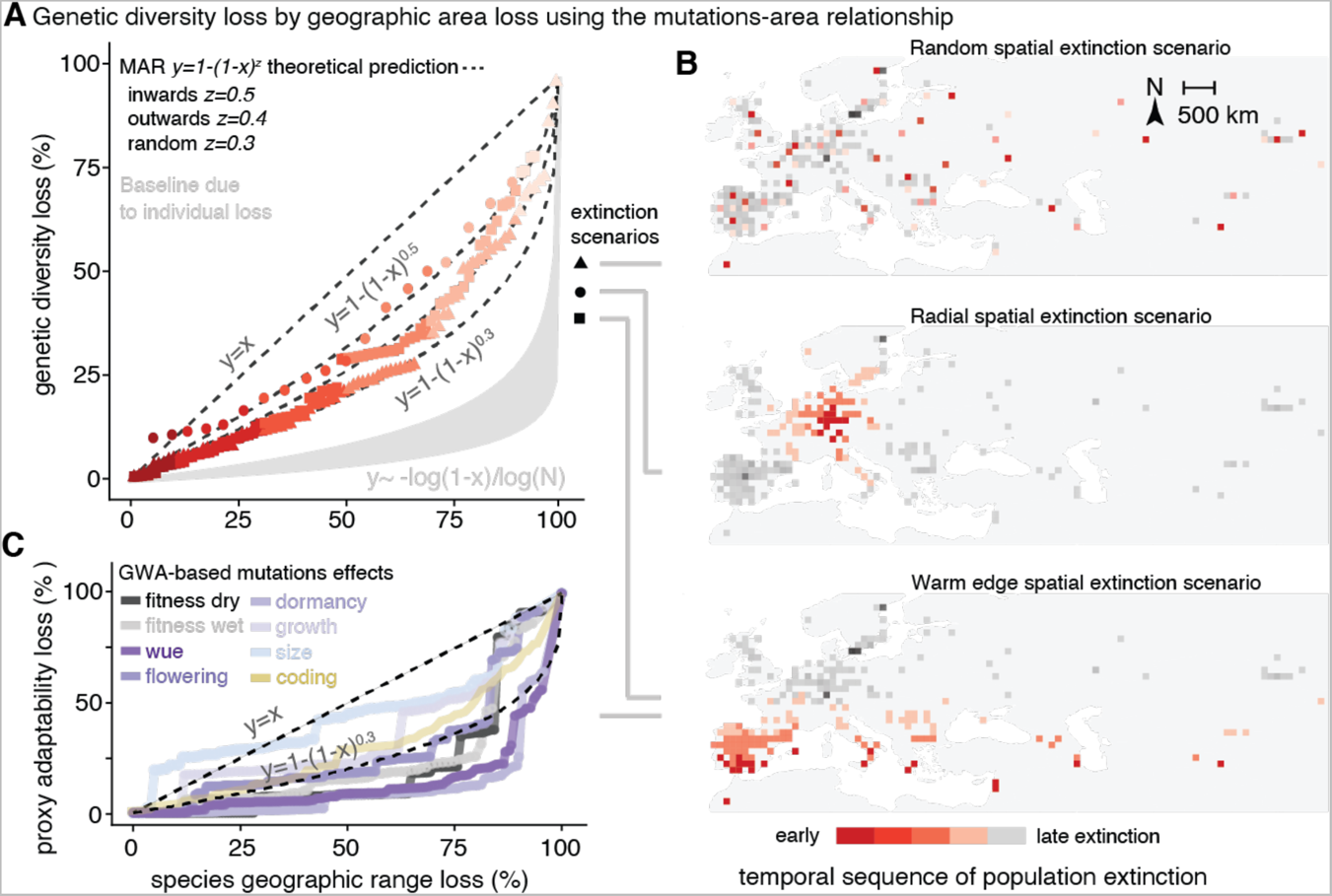
The power law of genetic diversity loss with range area loss. **(A)** Percentage of loss of total genetic diversity in *Arabidopsis thaliana* from several stochastic simulations (red) of local extinction in (B), and theoretical model projections of genetic diversity loss using the MAR (dotted lines). The expectation for genetic diversity loss based only on individuals is in grey (using starting populations of *N=10^4^-10^9^*) (**SM II.4**). **(B)** Cartoon of several possible range contractions simulated by progressively removing grid cells across the map of Eurasia (red/grey boxes) following different hypothesised spatial extinction patterns. (**C**) A metric of adaptive capacity loss during warm edge extinction in (B). Using Genome Wide Associations (GWA) to estimate effects of mutation on fitness in different rainfall conditions, water use efficiency [wue], flowering time, seed dormancy, plant growth rate, and plant size. Plotted are the fraction loss of the summed squared effects (∑*a^2^*) of 10,000 mutations from the top 1% tails of effects. We also plot (yellow) the fraction of protein-coding alleles lost (nonsynonymous, stop codon loss/gain, and frameshift mutations).

Since genetic diversity is ultimately created by spontaneous DNA errors passed onto offspring every generation, the loss of genetic diversity seems reversible, as these mutations could happen again. However, the recovery of genetic diversity through natural mutagenesis is extremely slow (57), especially for mutations affecting adaptation. Simulating a species undergoing only a 5–10% in area reduction, it would take at least *≈*140–520 generations to recover its original genetic diversity (2,100–7,800 years for a fast-growing tree or medium-lifespan mammal of 15 year generation length), although for most simulations, recovery virtually never happened over millennia (see **SM II.4–5, Fig. S11, SM III.6**).

To test the generality of the MAR, we searched in public nucleotide repositories for datasets of hundreds to thousands of whole-genome sequenced individuals for the same species sampled across geographic areas within their native ranges (**Table 1, SM IV**). In total, we identified 20 wild plant and animal species with such published resources and assembled a dataset amassing a total of 10,095 individuals of these species, with 1,522 to 88,332,015 naturally occurring mutations per species, covering a geographic area ranging from 0.03 to 115 million km^2^. Fitting MAR for these diverse species, we recovered *z_MAR_* values similar to *A. thaliana*, with many species overlapping in confidence intervals, with the exception of some outliers (mean (SE) *z_MAR_* = 0.31 (±0.038), median = 0.26, IQR = ±0.15, range=0.10–0.82, mean (SE) *z*_MAR_* scaled = 0.26 (±0.048). See **Table 1, SM IV, Fig. S22, Table S10**). Theoretical derivations show that *z_MAR_* is a consequence of fundamental evolutionary and ecological forces (mutation, dispersal, selection) and should range from 0 to 1, depending on the strength of population structure (**SM II.3**, see **Fig. S10** for its relationship with isolation-by-distance). These predictions were further confirmed by spatial population genetics coalescent and individual-based simulations in 2D and continuous space (**SM II.3**), as well as with mainland-island community assembly simulations according to the Unified Neutral Theory of Biodiversity (UNTB) (**SM V.3**).

**Table 1.**
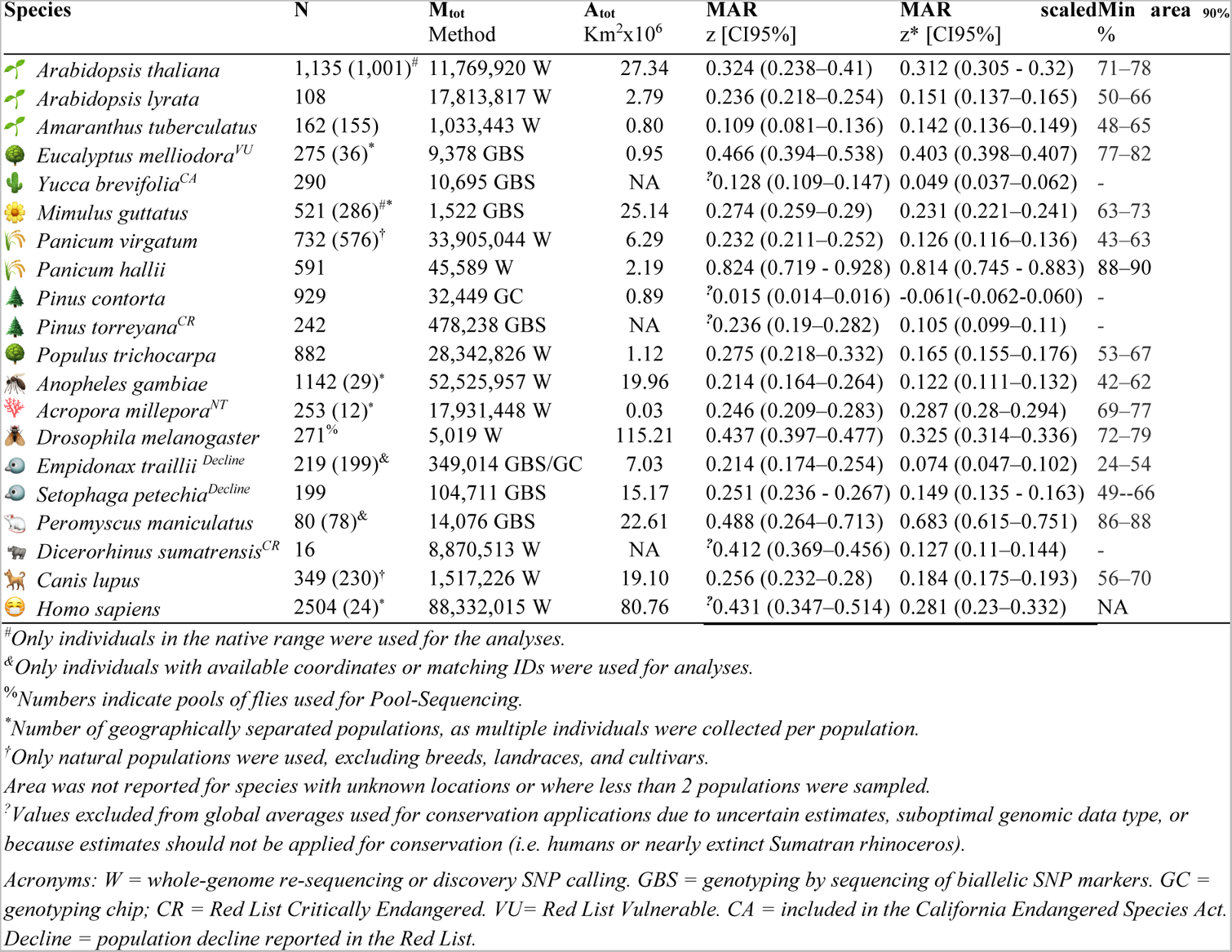
The mutations-area relationship across diverse species. Summary statistics of individuals sampled broadly across species distributions, sequencing method and mutations studied, and convex hull area extent of all samples within a species. The mutations-area relationship (MAR) parameter *z*, which captures how spatially restricted mutations are, including a scaled correction *z*\* for low sampling genomic effort. Percent area that needs to be kept for a species to maintain 90% of its genetic diversity, using the per-species MAR value estimates. Area predictions are not provided for threatened species, as these have likely already lost substantial genetic diversity and require protection of their full geographic range (Fig. 3).

| Species | N | M<sub>tot</sub><br>Method | A<sub>tot</sub><br>Km<sup>2</sup>x10<sup>6</sup> | MAR<br>$z$ [CI95%] | MAR<br>$z^*$ [CI95%] | scaledMin<br>area 90%<br>% |
| --- | --- | --- | --- | --- | --- | --- |
| *Arabidopsis thaliana* | 1,135 (1,001)<sup>#</sup> | 11,769,920 W | 27.34 | 0.324 (0.238–0.41) | 0.312 (0.305 - 0.32) | 71–78 |
| *Arabidopsis lyrata* | 108 | 17,813,817 W | 2.79 | 0.236 (0.218–0.254) | 0.151 (0.137–0.165) | 50–66 |
| *Amaranthus tuberculatus* | 162 (155) | 1,033,443 W | 0.80 | 0.109 (0.081–0.136) | 0.142 (0.136–0.149) | 48–65 |
| *Eucalyptus melliodora*<sup>VU</sup> | 275 (36)<sup>*</sup> | 9,378 GBS | 0.95 | 0.466 (0.394–0.538) | 0.403 (0.398–0.407) | 77–82 |
| *Yucca brevifolia*<sup>CA</sup> | 290 | 10,695 GBS | NA | <sup>?</sup>0.128 (0.109–0.147) | 0.049 (0.037–0.062) | - |
| *Mimulus guttatus* | 521 (286)<sup>#*</sup> | 1,522 GBS | 25.14 | 0.274 (0.259–0.29) | 0.231 (0.221–0.241) | 63–73 |
| *Panicum virgatum* | 732 (576)<sup>†</sup> | 33,905,044 W | 6.29 | 0.232 (0.211–0.252) | 0.126 (0.116–0.136) | 43–63 |
| *Panicum hallii* | 591 | 45,589 W | 2.19 | 0.824 (0.719 - 0.928) | 0.814 (0.745 - 0.883) | 88–90 |
| *Pinus contorta* | 929 | 32,449 GC | 0.89 | <sup>?</sup>0.015 (0.014–0.016) | -0.061(-0.062-0.060) | - |
| *Pinus torreyana*<sup>CR</sup> | 242 | 478,238 GBS | NA | <sup>?</sup>0.236 (0.19–0.282) | 0.105 (0.099–0.11) | - |
| *Populus trichocarpa* | 882 | 28,342,826 W | 1.12 | 0.275 (0.218–0.332) | 0.165 (0.155–0.176) | 53–67 |
| *Anopheles gambiae* | 1142 (29)<sup>*</sup> | 52,525,957 W | 19.96 | 0.214 (0.164–0.264) | 0.122 (0.111–0.132) | 42–62 |
| *Acropora millepora*<sup>NT</sup> | 253 (12)<sup>*</sup> | 17,931,448 W | 0.03 | 0.246 (0.209–0.283) | 0.287 (0.28–0.294) | 69–77 |
| *Drosophila melanogaster* | 271<sup>%</sup> | 5,019 W | 115.21 | 0.437 (0.397–0.477) | 0.325 (0.314–0.336) | 72–79 |
| *Empidonax traillii*<sup>Decline</sup> | 219 (199)<sup>&</sup> | 349,014 GBS/GC | 7.03 | 0.214 (0.174–0.254) | 0.074 (0.047–0.102) | 24–54 |
| *Setophaga petechia*<sup>Decline</sup> | 199 | 104,711 GBS | 15.17 | 0.251 (0.236 - 0.267) | 0.149 (0.135 - 0.163) | 49--66 |
| *Peromyscus maniculatus* | 80 (78)<sup>&</sup> | 14,076 GBS | 22.61 | 0.488 (0.264–0.713) | 0.683 (0.615–0.751) | 86–88 |
| *Dicerorhinus sumatrensis*<sup>CR</sup> | 16 | 8,870,513 W | NA | <sup>?</sup>0.412 (0.369–0.456) | 0.127 (0.11–0.144) | - |
| *Canis lupus* | 349 (230)<sup>†</sup> | 1,517,226 W | 19.10 | 0.256 (0.232–0.28) | 0.184 (0.175–0.193) | 56–70 |
| *Homo sapiens* | 2504 (24)<sup>*</sup> | 88,332,015 W | 80.76 | <sup>?</sup>0.431 (0.347–0.514) | 0.281 (0.23–0.332) | NA |

<sup>#</sup>Only individuals in the native range were used for the analyses.
<sup>&</sup>Only individuals with available coordinates or matching IDs were used for analyses.
<sup>%</sup>Numbers indicate pools of flies used for Pool-Sequencing.
<sup>\*</sup>Number of geographically separated populations, as multiple individuals were collected per population.
<sup>†</sup>Only natural populations were used, excluding breeds, landraces, and cultivars.
Area was not reported for species with unknown locations or where less than 2 populations were sampled.
<sup>?</sup>Values excluded from global averages used for conservation applications due to uncertain estimates, suboptimal genomic data type, or because estimates should not be applied for conservation (i.e. humans or nearly extinct Sumatran rhinoceros).
Acronyms: W = whole-genome re-sequencing or discovery SNP calling. GBS = genotyping by sequencing of biallelic SNP markers. GC = genotyping chip; CR = Red List Critically Endangered. VU = Red List Vulnerable. CA = included in the California Endangered Species Act. Decline = population decline reported in the Red List.

Although we expect species-specific traits related to dispersibility or gene flow to affect *z_MAR_* (e.g. migration rate and environmental selection in population genetic simulations significantly influences *z_MAR_*, **Table S2**), no significant association was found between *z_MAR_* and different ecologically-relevant traits, mating systems, home continents, etc., for the 20 species analysed. Perhaps this is simply that there are still too few species that have large population genomic data to find such a signal (**Table 1, Table S12–13)**. Nevertheless, the relative consistency of *z_MAR_* across largely different species may be promising for conservation purposes, as an average *z_MAR_* ~0.3 (IQR ±0.15, **Table 1, Table S11**) could be predictive of large-scale trends of genetic diversity loss in many range-reduced species that lack genomic information. Further, although species will naturally have different starting levels of total genetic diversity prior to range reductions, for instance, due to genome size, structure, or mating system differences (*26*), the application of *z_MAR_* provides relative estimates of genetic diversity loss. For instance, assuming *z_MAR_* ~0.3, we would predict that an area reduction of ~50% creates an approximate loss of ~20% of genetic diversity relative to the total genetic diversity of a given species.

Finally, we used MAR to estimate the average global genetic diversity loss caused by pre-21^st^ century land transformations. Although accurate species-specific geographic area reduction data in the last centuries are scarce, we leveraged global land cover transformations from primary ecosystems to urban or cropland systems (*3, 27*) (**Table S14–15**). Using the average scaled *z*_MAR_* (**Table S18**) and several global averages of Earth’s land and coastal transformations for present day (38% global area transformation from (*27*), 34% from (*28*), and 43-50% from (*29*)), we estimate a 10-16% global genetic diversity loss on average across species (**Fig. 3A**). While these estimates may correctly approximate central values across species in an ecosystem, we expect a substantial variation in the extent of loss across species, ranging theoretically from 0 to 100% (**Fig. 3, Fig. S26**). One cause of this variation is the heterogeneity in land cover transformations across ecosystems; for example, more pristine high-altitude systems have only lost 0.3% of their area, while highly managed temperate forests and woodlands have lost 67% (**Fig. 3B, Table S14–15)**.

**Fig. 3.**
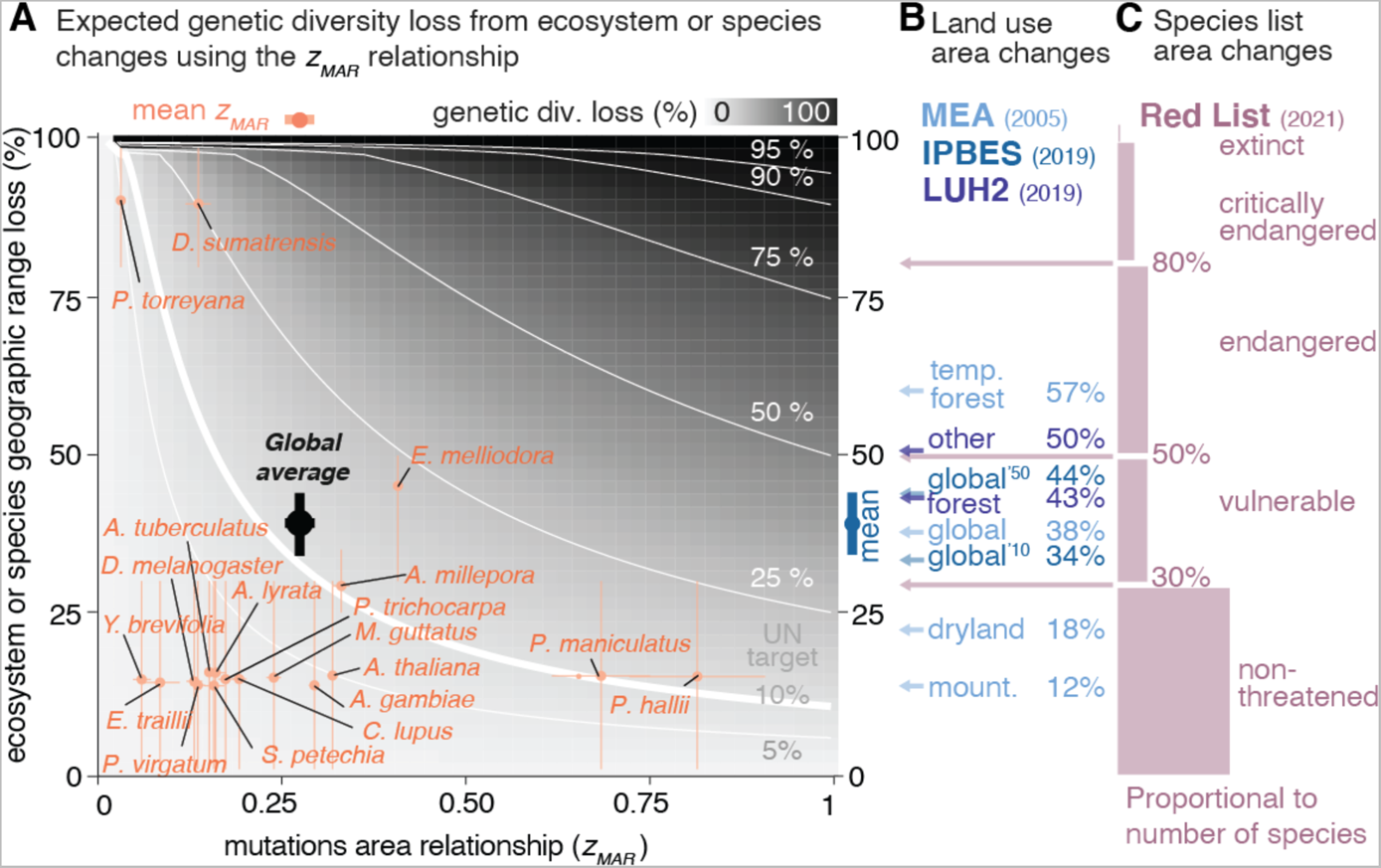
The parameter space of genetic diversity loss mapping pre-21^st^ century ecosystem transformations and species threat categories against possible values of the mutations-area relationship. **(A)** Possible values of two key parameters, the mutations-area relationship scaling parameter (MAR) and % of area reduction of a species geographic range (as a proxy of entire ecosystem transformation). The theoretical % of genetic diversity loss is represented as filled grey colour, with isolines in white. Estimates of scaled *z*_MAR_* from Table 1 per species are in orange with their 95% confidence intervals (for unscaled *z_MAR_* see Fig. S23). Although exact area losses per species are unknown, species are plotted based on their IUCN Red list (C) status, using the broad ranges of minimum and maximum recent population or area decline per category. The global average is calculated with the average *z_MAR_* across species and % of the Earth transformed from IPBES. **(B)** Percentage of transformed ecosystem area from the Millennium Ecosystem Assessment (MEA) (*27*) are represented by light blue arrows, from the Intergovernmental Science-Policy Panel for Biodiversity and Ecosystem Services (IPBES) (*28*) for 2010 and 2050 are dark blue arrows, and from the Land Use Harmonization 2 (LUH2) dataset (*29*) are in dark purple. (**C**) The minimum criterion value of population or geographic area loss to be classified in each category of the IUCN Red List are indicated with pink arrows (the near threatened category does not have a range of values, instead we used 30% ±10%). The number of plant species (for which population abundance loss approximates area loss) included in each category is shown as box sizes (*1*). The IUCN ranges were used to place ranges of estimates in (A) per species.

Another cause for the variability in genetic loss among species (even within the same ecosystem) may be their differential geographic ranges and abundances, life histories, or species-specific threats. We gathered data from species red-listed by the International Union for Conservation of Nature (IUCN) (*1*), which evaluates recent population or geographic range area reduction over ±10 years / ±3 generations to place assessed species in different threat categories using several thresholds (guidelines for assessments and thresholds available at www.iucn.org). Again, assuming that with the average *z_MAR_ ~*0.3 we can capture general patterns, we translate these category thresholds into genetic diversity loss (**Fig. 3C,** see **SM V, Table S17)**. *Vulnerable* species, having lost at least 30% of their geographic distribution, may have experienced >9% of genetic diversity loss, *endangered* species, which have lost over 50% of their geographic distribution, should have incurred >16% of genetic diversity loss, and *critically endangered* species, with over 80% area reduction, likely suffered >33% of genetic diversity loss (**Fig. 3B**). This clearly showcases that even species in no imminent risk of extinction (e.g. least concern, near threatened, vulnerable), such as the majority of species for which population genomic data exists, may already be losing substantial genetic diversity (**Fig. 3A**).

The ultimate challenge is to understand how genetic diversity loss relates to loss of adaptive capacity of a species. To this end, we leveraged the extensive knowledge of the effect of mutations in ecologically relevant traits in *A. thaliana* from Genome-Wide Associations (GWA) (**Fig. 2C, SM III**). We again conducted spatial warm edge extinction simulations, this time tracking metrics of adaptive capacity, including the total sum of effects estimated from GWA of remaining mutations (∑*_i_ a_i_* for *i*=1…10,000 variants of putative *a_i_* effect), the additive genetic variance (*Va=* ∑*_i_ p_i_(1-p_i_)a_i_^2^*, which accounts for each variant’s population frequency *p_i_*), and the loss of nonsynonymous mutations (**SM III.5**). Although determining the effect of mutations through GWA is technically challenging even in model species (*30, 31*), and variants may even be either deleterious or advantageous depending on genomic backgrounds (*32*) or environments (*33*), our simulations suggest putatively functional mutations may be lost more slowly (*z<0.3,* **Fig. 2C**) than neutral genetic diversity (**Fig. 2A)**. In fact, the additive variance *Va* parameter, often equated to the rate of adaptation, appears rather stable (*34*) until just before the extinction event when it sharply collapses (**Fig. S21**; see also **Fig. 2C,** and **SM II.3.4** for simulations that replicate this pattern). This is analogous to the famous “rivet popper” metaphor where ecosystem structure and function may suddenly collapse as species are inadvertently lost (*35*). Projections of the MAR using genome-wide variation may crucially serve as early conservation tool in non-threatened species (*36, 37*), before species reach accelerating collapsing extinction dynamics—an acceleration that we expect to be even more dramatic due to elevated drift and accumulation of deleterious mutations of small critically-endangered populations (*38, 39*).

To achieve the recently published United Nations target to protect “at least 90% of genetic diversity within all species”(*13*), it will be necessary to aggressively protect as many populations as possible for each species. Here, we have discovered the existence of a mutations-area relationship (MAR) and provided a mathematical framework to forecast genetic diversity loss with shrinking geographic species ranges. The MAR contrasts with existing studies on the risk of losing entire species by focusing on quantifying the magnitude and dynamics of genetic diversity loss likely ongoing in most species. This framework demonstrates that even with conservative estimates, substantial area protection will be needed to meet the UN Sustainable Development Goals. For vulnerable or endangered species, we may have likely already failed.

## ADDITIONAL INFORMATION

### Author contribution

M.E.-A. conceived and led the project. M.E.-A., J.P.S., M.R., S.H., L.G., L.C., L.L., S.T.A., V.P., E.Z., P.L.M.L., C.C.K., T.B., C.W. conducted research, all authors interpreted the results and wrote the manuscript.

### Data availability

The analysed datasets are publicly available or were shared by authors upon request (see Supplementary Materials for details). Code is available at Github (https://github.com/moiexpositoalonsolab/mar) and Zenodo (https://doi.org/10.5281/zenodo.6408624).

## Acknowledgements

We are grateful for many researchers who made their genomic data publicly available and thus made this research possible: the 1001 Arabidopsis Genomes Consortium, the Anopheles gambiae 1000 Genomes Consortium, the 1000 (Human) Genome Consortium, the European “Drosophila Evolution over Space and Time” (DEST) Consortium, MacLachlan et al., Fuller et al., Ruegg et al., Kingsley et al., Schweizer et al., and Royer et al.. In addition, we are grateful to all the authors who directly shared intermediate genome variant files: Sujan Mamidi, John Lovell, Dan Jacobson, Manesh Shah, Julia Kreiner, Kay Lucek, Yvonne Willi, Juan D. Palacio Mejia, Justin Borevitz, Megan Supple, Mario Vallejo-Marin, Nicolas Dussex, Lionel Di Santo, and Jill Hamilton. We thank John Wiens, Sue Rhee, Detlef Weigel, Ellie Armstrong, Marty Kardos, Dmitri Petrov, Rob Colwell, Rasmus Nielsen, Anna Michalak, Sean Hoban, Jonathan Pritchard, and members of the Moi and Mordecai Labs for comments, discussion, or references. M.E.-A. is supported by the Office of the Director of the National Institutes of Health’s Early Investigator Award with award number: 1DP5OD029506-01; by the U.S. Department of Energy, Office of Biological and Environmental Research, grant number: DE-SC0021286; and by the Carnegie Institution for Science. J.P.S. is supported by an NIH training grant (5T32HG000044-23), and S.T.A by the NIGMS Center of the NIH under award number T32GM007276. S.H. and C.L.W. are supported by Stanford’s Center for Computational, Evolutionary, and Human Genomics. P.L.M.L. is supported by a Human Frontier Science Program Long-Term Fellowship (LT000330/2019-L). M. R. is supported by the NSF’s Plant Genome Postdoctoral Research Fellowship in Biology. L. L. and L.G. are supported by NSF’s GRFP. Computational analyses were done on the High-Performance Computing clusters *Memex* and *Calc* supported by the Carnegie Institution for Science.

## Supplementary Materials

### SUPPLEMENTAL METHODS

#### I. Background on species biodiversity and biogeography

##### I.1 Theoretical models of biodiversity

Studies in biogeography have modelled the species-area relationship with several functions. Below we summarise the different approaches using an example of richness of S = 100 species, with variable abundance or area, A.

We may visualise the different areas or abundances of species as a frequency histogram (Fig. S1, Preston plot), with x-axis: logarithm of abundance bins (historically log2 as a rough approximation to the natural logarithm), and y-axis: number of species at given abundance. Alternatively, as a rank-abundance diagram (Fig. S1, Whittaker plot): x-axis: species list, ranked in order of descending abundance (i.e. from common to rare), and y-axis: logarithm of % relative abundance.

**Fig. S1.**
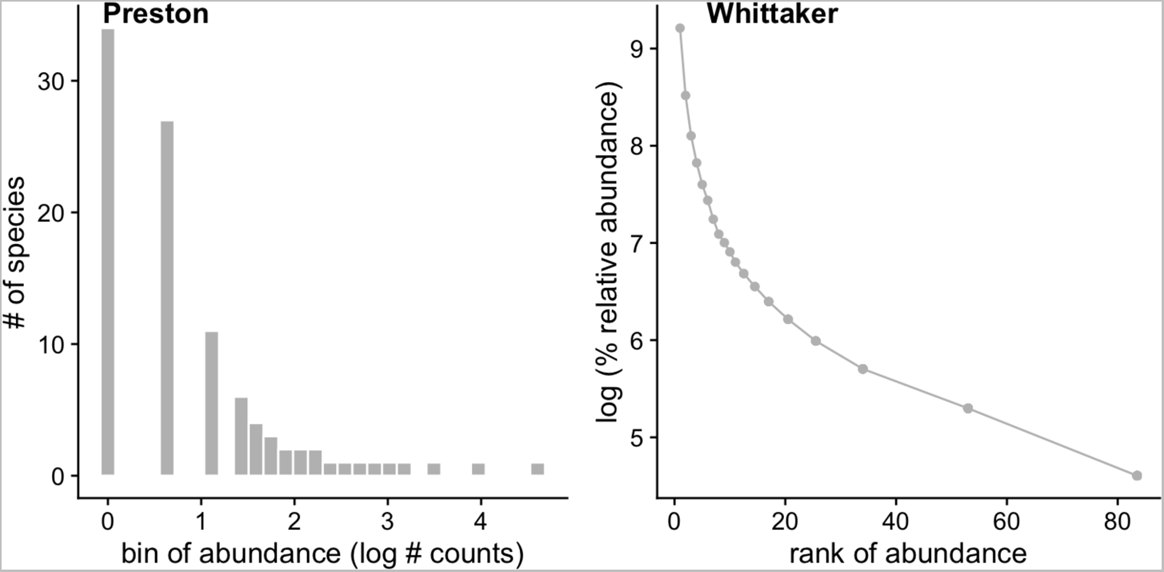
Example of typical plots used for species abundance curve studies. Due to their strong skew, Species Abundance Curves are often plotted using the Preston plot (left) where the x axis represents bins of log2 abundances (also referred to as octaves), or using the Whittaker plot (right) where the x axis is the rank of each species in a dataset and y axis the species’ relative abundance.

###### I.1.1. Niche apportionment approaches

A series of theoretical deterministic and stochastic “niche apportionment models” have been put forward (summarised in (*1*) or (*2, 3*)).

The Motomura (*4*) geometric series suggests that each species that arrives takes half the area. The first would take 50%, the second 50% of 50%, and so forth, which can be expressed as:

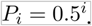

Similarly, one can imagine that as a species colonises a habitat, it takes up a fraction different than 50%. This gives a geometric series with parameters which can be written as

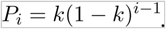

Other geometric series-related models include stochasticity, where *k* instead of being a fixed parameter is a random uniform variable and there is a *k_i_* each time *i* a new species arrives to the ecosystem. The “dominance preemption” model draws from 50-100% at any new arrival of a species, the random fraction model draws from 0-100%. Then the abundance of a species depends on the stochastic process of previous *f = 1…i-1* species arriving first:

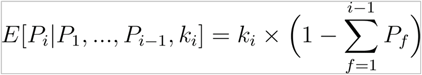

Another approach is the broken stick by MacArthur (*5*), which theorised a habitat is broken into *S-1* places at random, which creates fractions of an area. Then the relative area of a species is:

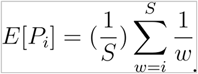

###### I.1.2. Niche statistical approaches of species sampling

Differently from niche partitioning functions, statistical approaches such as the log-series from Fisher (*6*) and log-normal from Preston (*7*) are probability distributions, and approach modelling in a conceptually different way: they model the sampling process of species collections given an underlying relative abundance (see below).

Statistical-based derivations probably began with Fisher (*6*), with the log-series distribution. It assumes that species abundances in the community are independent identically distributed variables, sampling is a Poisson process, sampling is done with replacement, or the fraction sampled is small enough to approximate a sample with replacement. Here,

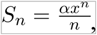

Where is a constant *x* ∈[0,1] related to the sample dataset (typically close to 1), 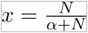, and α is a new constant term (ecosystem-specific) that is used as a measure of biodiversity. Fisher proposed the number of species could be estimated as:

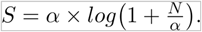

Finally, Preston (*8*) posed that the skewness of previous proposals is due to lack of sampling. With little data, common species are collected sooner, but with more abundant sampling, the rarest species are also well-sampled and have abundances well above 0. Preston then proposed that the octaves (bins of doubling abundance) follow a normal distribution, making the raw abundance log-normal distributed. Given *S*_0_ is the number of species in the model octave of abundance and a variance composite of the log-Normal *σ*^2^, the number of species per abundance (octave) bin *R*(=*log(n)*) is:

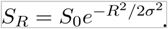

The Unified Neutral Theory of Biodiversity (UNTB) by Hubbell (*1*) takes a stochastic approach of a community with immigrants, extinctions, and speciation in continuous dynamics. Interestingly, the UNTB’s key parameter, *θ*, coincides with Fisher’s *α*, as the log-series is a limiting case of UNTB. Hubbell’s discovery was that *α=2J_m_v*, where *J_m_* is the size of the external metacommunity that provides migrants of species to the focal community, and *ν* is the speciation rate. Alonso and McKane (*9*) derived the so-called Metacommunity Zero-Sum Multinomial (MZSM) distribution from the UNTB. In practice, both distributions have almost-identical fits (lines completely overlapping in Fig. S2 below).

**Fig. S2.**
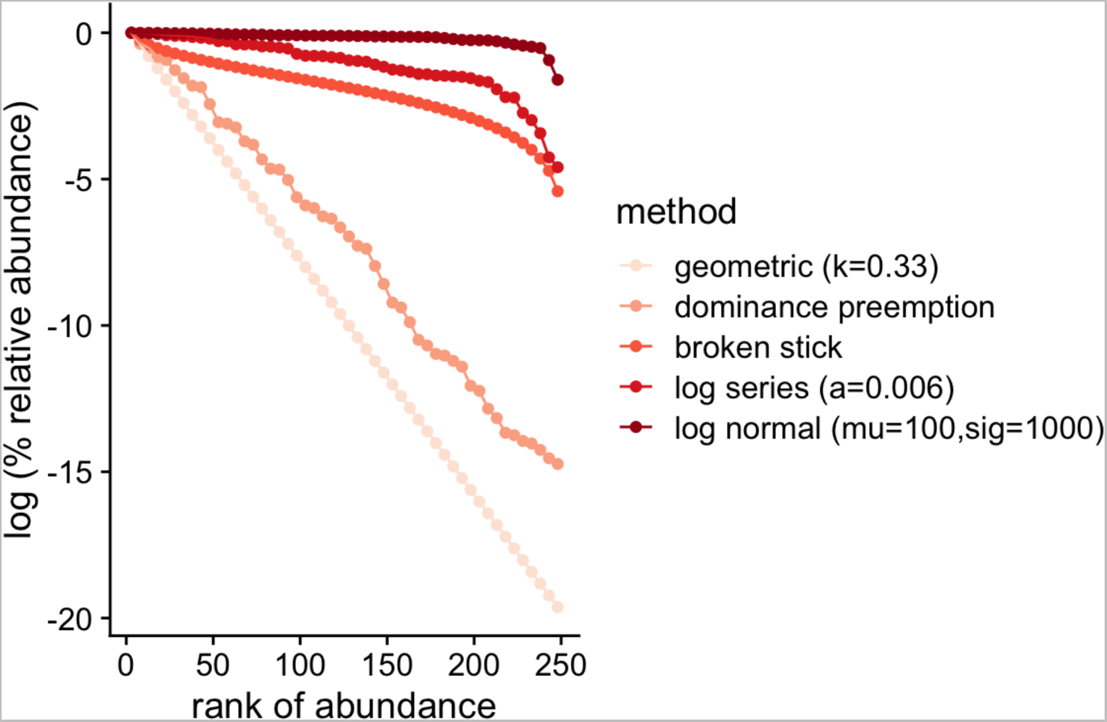
Summary of theoretical models of Species Abundance Curves. Five niche partitioning or statistical models shown in a Whittaker plot. The different models expect different levels of evenness in abundance across the species in the community, from the lowest (geometric series) to the highest (log-normal).

##### I.2 Metric of species diversity

Although a number of metrics exist to measure species diversity, such as the Shannon index, 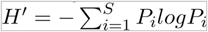 (with *Pi* the relative proportions of species abundances) or Fisher’s non-dimensional parameter, the study of species abundances and area relationships has focused on species richness *S*, that is, the total number of species in a given location or area. Below we therefore focus on species richness.

##### I.3 Biogeography of species and extinction

###### SAD and SAR connection

Due to many species being rare, it is expected that as researchers sample an area, the most common species will be sampled first, and as the area studied increases, more and more species will be discovered. This is thought to happen following a power law relationship, where the number of species in that area *S_A_* increases with the sampled area *A*, with scaling *z* (slope in a log-log plot), and with a constant *c*:

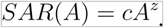

Preston (*7*) derived theoretically that from a log-normal series, one would expect *z=*0.27, under a number of assumptions (Fig. S3). This has been empirically shown to be close to reality (*7, 10*), although there is some variation across ecosystems and spatial scales.

**Fig. S3.**
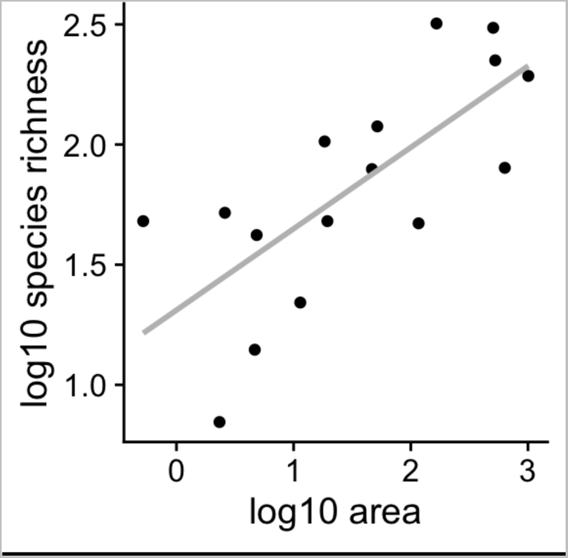
Example of a Species-Area Relationship in Galapagos Islands. Classic species richness dataset from the Galapagos Islands (Preston, 1962). It depicts species richness as a function of island area in a log-log plot.

##### I.4 Estimating extinction of species from the species area relationship

The first estimates of species extinction used the SAR relationship. Given a reduction of ecosystem area, *A*, by an area of *a* (*11, 12*). If these areas, as well as the SAR scaling, *z*, are known, then one can predict the number of species in the future as:

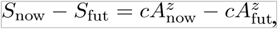

However, we are normally interested in the fraction of species that will go extinct *X_s_* so we can take the ratio:

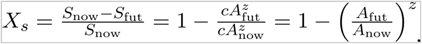

#### II. Population genetics models and the site frequency spectrum

##### II.1 The Wright-Fisher model and the site frequency spectrum

Statisticians and population geneticists from the 20^th^ century, Wright and Fisher, built a simple statistical model of evolution of a population. It assumes that each generation a population of *N* monoecious (hermaphrodite) individuals mate randomly to create a new generation of *N* individuals and then immediately die so that only *N* individuals remain in the population at any given time. This random sampling process causes the frequency of a variant in one generation to possibly differ from its frequency in the previous generation—a process known as genetic drift.

When a nucleotide mutation or variant (e.g. ACG**A**A → ACG**T**A) emerges by a random process of, for instance, DNA replication error, it will first be in 1/*N* individuals (if we consider these diploid, 1/2N chromosomes). Through random sampling that **T** mutation may be lost, stay at the same frequency, or randomly move to higher frequency. Although rarely, just by chance, the mutation may reach 100% frequency. This results in a “commonness of rarity” when looking at mutations in a population, as we have seen in previous sections for species. Since these genetic drift dynamics affect all mutations genome-wide, we therefore expect the majority of mutations to be absent, or rare, and only a much smaller proportion of variants to be at moderate or high frequencies.

The site frequency spectrum (SFS) refers to the distribution of frequencies of variants in a population. This is the number of sites at which we observe a variant at frequency *q* in a sample of *n* individuals. To derive the expected SFS distribution, we turn to Kingman’s Coalescent (*13*). Both models describe the same ideal population of random mating, constant population size, and mutations emerging at a low rate and drifting in frequency. But while the Wright-Fisher model describes the dynamics of a whole population forward-in-time, the Kingman’s Coalescent describes the genealogy of a sample of individuals from a population, going backward in time. By building a model around the individuals that are sampled or that survived, rather than of an entire population, the Coalescent provides a simpler way to derive expectations in small populations or in cases, for example here, where a limited sample of genomes are sequenced. Using the Coalescent (see (*14*) for details), one obtains that the expected number of mutations of a given abundance, *n*, is inversely related to their frequency, *q*:

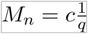

for some constant c that depends on the mutation rate and the population size. This SFS from population genetics theory is remarkably similar to the Species Abundance Relationship. In fact, Fisher himself (*15*) derived an expression similar to the above.

Rearranging terms, one can see this is a constrained version of the log-series Probability Mass Function (PMF), which Fisher also proposed for the distribution of species abundances (*6*). Below, one can graphically see the similarities (Fig. S4):

**Fig. S4.**
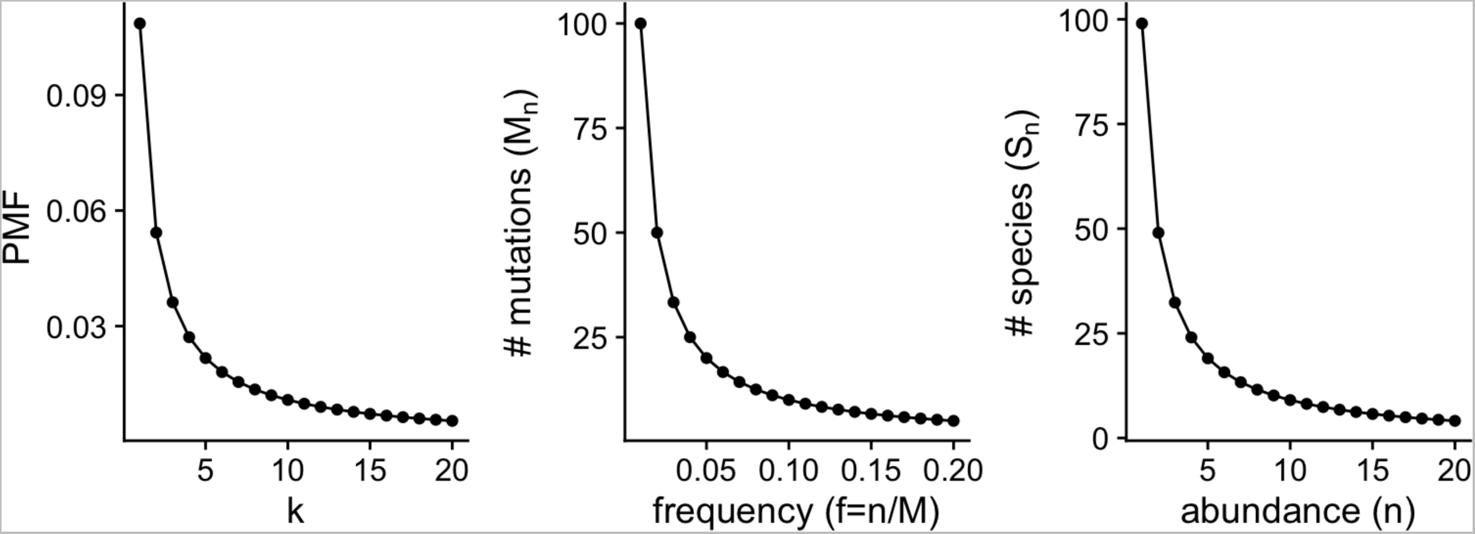
Similarity between the Species Abundance Distribution and the Site Frequency Spectrum. Left is the Probability Mass Function of the log-series (p=0.999), center is the SFS (N=100, c=1), and right is the log-series-based abundance of species (alpha=100, N=10000).

Keeping the abundance, *n*, constant (and low), when the number of individuals *N* → ∞, we know that the constant *x* from Fisher’s SAD approaches 1, 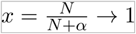. Then, we can rewrite the number of species at any given abundance (*S_n_*) as:

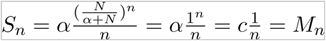

So both have the same form as the log series PMF: 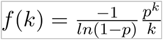 when *p* → 1. In the next section we will see that the constants of the SAD and the SFS are proportional to species and mutation diversity, although the Site Frequency Spectrum (SFS) is a specific case of SAD. One can also see that because the constant in the SFS is the population scaled mutation rate, *c* = θ= *Ne*μ, and Fisher’s α≈θ for large N.

##### II.2 Metrics of genetic diversity

In population genetics, multiple measurements of genetic diversity have been put forward. The most straightforward is the allelic richness, also number of mutations, or also called the number of segregating sites. Segregating sites, *M*, is the direct equivalent of the species richness, *S*, and it depends on the number of samples used and length of DNA sequence explored (Note: we use the non-standard notation, *M*, as the standard in population genetics is *S* [for segregating sites] but this is already in use for species richness. We then use *M* for mutations and *S* for species). This metric can also be thought of as the area under the curve of the SFS. Two other metrics that describe the SFS but that aim to be sequence-length- and individual independent are Watterson’s Theta, θ_W_, and Nucleotide diversity,π, (also called θ_π_). These two metrics of diversity are identical at population equilibrium and are estimates of *4N_e_μ* (when the SFS follows a 1/q relationship), with effective population size *N_e_* and per-generation mutation rate *μ*, whereas they differ in non-equilibrium demographics, under natural selection, or under other behaviors not considered in the Wright-Fisher neutral model, such as different mating systems (*16*).

First, πis described as:

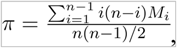

and θ_π_ as:

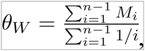

where 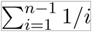 is the *n-1^th^* Harmonic number, which serves to scale the segregating sites based on the assumption that the abundance of mutations follows a 1/q SFS. The diversity metrics π and θ_π_ are both functions of the SFS, as opposed to Fisher’s αfrom the Species Abundance Distribution, which is a parameter that changes the shape of the distribution.

Although often nucleotide diversity π is reported as a typical measure of genetic diversity of a species, since it can be calculated for a single genome and it captures the process of inbreeding of a population (*17*), classic literature relating germplasm management for conservation and breeding has advocated for allelic richness (*18*).

##### II.3 Spatial genetics and the mutations-area relationship (MAR)

Since its inception, a number of concepts in population genetics have dealt with genetic variation in populations of different sizes, or populations separated in space. For instance, one classic result in population genetics is the relationship of *π ≈ 4N_c_ μ,* which relates genetic diversity *π* with the effective population size *N_e_* and the mutation rate of the species *μ.* A relationship which is still studied nowadays in an effort to reconcile data with theory (*17*).

In 1943, Sewall Wright turned to study the genetics of multiple populations within a species. He proposed that populations sampled further apart geographically must differ more in allele frequency due to more independent drift (*19*), leading to the commonly used correlation between geographic distance and the metric of differentiation *F_ST_*. Most prominently, the use of correlation in the accumulation of mutations of populations that are geographically close or share evolutionary history has been uncovered using dimensionality reduction approaches such as PCA (*20*).

Despite these enormous advances in understanding spatial genetic structures, surprisingly little quantitative work has been done to parametrize the loss of genetic diversity by direct loss of habitat.

Because of the abundance of rare mutations in populations, it is straightforward to think that the more area and individuals sampled, the more segregating sites will be found. Analogous to the Species Area Relationship (SAR), *S=cA^z^*, we should thus be able to estimate the equivalent scaling for a mutations-area relationship (MAR):

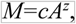

with a scaling *z = z_MAR_*, which corresponds to the slope of best fit in a log-log-plot of *A* and *M* for a given species. (Other functions are often fit empirically for SAR datasets, which we explore later in section III.3. We work with the power law because of its historical use, mathematical convenience, and because other more complicated functions only improved fitting marginally, see Table S4).

This differs from other efforts to understand the number of segregating sites or heterozygosity differences across species that differ in their total census size or geographic distribution (*21, 22*). The MAR instead is built within a species, as its ultimate aim is to relate the number of mutations left in a species as it loses spatial populations.

Below we derive what are the expectations of MAR taking two opposite scenarios of neutral population evolution, and study how many segregating sites or mutations *M* are discovered with increasing area in the simulations. We further test the scenario of meta-populations in space with varying migration rates and neutral or natural selection processes.

###### II.3.1 Panmictic population

The expected number of mutations, *M*, is a constant that depends on the mutation rate, *μ*, and the expected total branch length of the population genealogy, *L*, with *M=μL*. Under the coalescent, the total branch length is equal to the number of lineages or individuals sampled from the population, *n*, times the time of the genealogy during which there are such lineages, *T_n_*, plus *n-1* times the time in the genealogy with such number of lineages, and so forth:

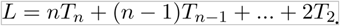

Under the coalescent,

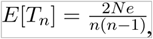

and thus:

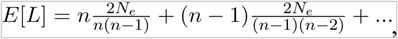

which simplifies to

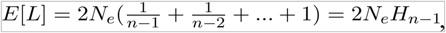

where *H_n-1_* is the (n-1)th harmonic number. This is of course related to one of the diversity metrics (section II.2), where Watterson’s Θ_W_ scales the number of segregating sites (*M*) by the harmonic number of sampled individuals. This is based on the expectation that as more individuals are sampled, we expect to discover more mutations proportional to the above harmonic number. Because such number is not so easy to work with to create an expectation for *z_MAR_*, we further simplify this expectation following the Taylor expansion approximation of the harmonic number:

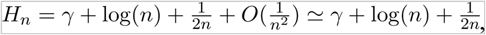

which we can further approximate as:

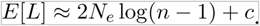

Therefore, assuming a constant mutation rate and effective population size (*N_e_*) under panmixia, *M* grows following *log(n)*. In such a case, a log-log plot (typical power law plot) does not display a linear relationship, and the slope is asymptotic to *z → 0* for *N → ∞*. On the other hand, with low values of *x* (area or individuals sampled close to 0), the slope *z_MAR_* will be incorrectly high. We can show this effect trivially by studying the local derivative of the function *log_10_(M) = log_10_(log(N))*. The local slope of that function is an approximation of our *z_MAR_* parameter. This can be locally estimated at any given point *N* by taking the derivative:

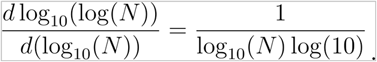

The implication of this nonlinear function is that if we sampled only few individuals or areas of a species (e.g., n=100), even if this species was completely panmictic we would expect a non-zero *z_MAR_*, a value that will change with sampling effort. We can roughly approximate *z_MAR_* by the local slope of the number in the midpoint of the graph, e.g., for n=100 we look at the slope at *n=50*, and obtain *1/(log_10_*(*50*) *x log(10))≅ 0.256*. Therefore, with small sample sizes, this parameter will not be helpful to understand whether a species behaves panmictically or is limited by migration, which may be problematic for estimates of genetic diversity loss later. We can visualise our expectation of the *z_MAR_* under panmixia plotting the first derivative above (Fig. S5). Because—as we will show below—we do expect a power law relationship under a migration-limited scenario, *z_MAR_* should theoretically not change with sample size. The graphical study of the (non-)linearity of the log-log plots between the number of mutations and area sampled should be diagnostic to this problem (We see for instance that *Pinus contorta* has a highly nonlinear relationship, likely due to the use of ascertained intermediate frequency markers instead of genome-wide data, Fig. S22).

Finally, we used msprime (*23*) to corroborate this finding (z_MAR_ being constant with respect to sample size) with simulations, simulating 1600 demes in a 40×40 grid of demes or populations of *N=N_e_=1000* that are completely panmictic (universal gene flow or dispersal, so this is equivalent to a single panmictic deme). We observed the *z_MAR_* for *t=100…10,000* generations in *log_10_* increments. After this time, we sample *n=1…100* individuals in increasingly large groups of adjacent demes. The range of estimates of *z_MAR_* in these simulations was 0.07-0.15.

Fig. S5 indicates that the minimum average *z_MAR_* even under panmixia would continuously increase with lower numbers of individuals of a species sampled. This is due to the fact that the site frequency spectrum is not fully sampled with small numbers of individuals. Therefore, we devised an approach to rescale *z_MAR_*.

**Fig. S5.**
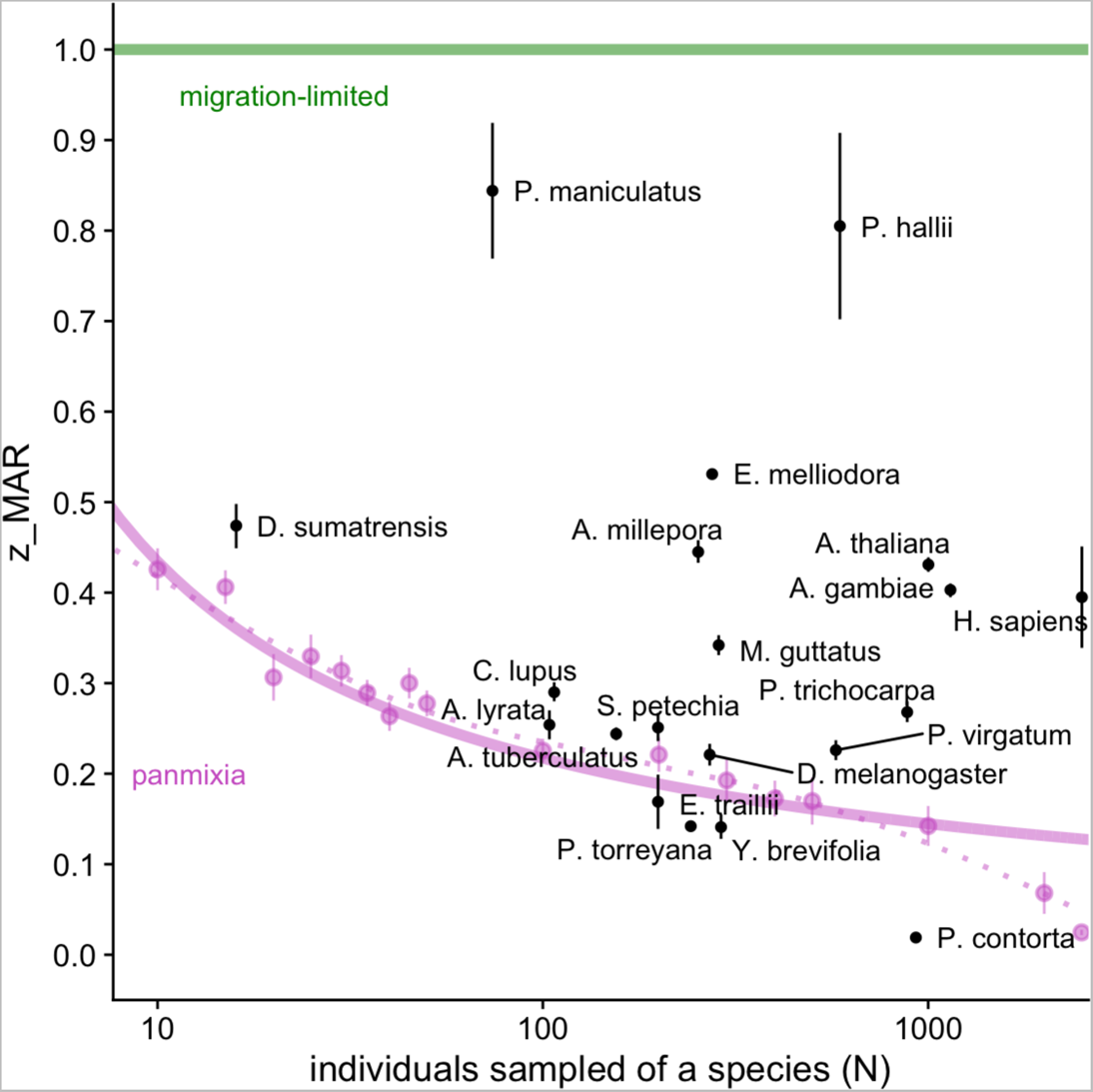
Expected ranges of z _MAR_ given sample sizes. For increasing numbers of individuals sampled, we plot the expected mean z_MAR_ under two theoretical trends of a migration-limited (green) and a panmictic (purple) species (Purple dots indicate averages from SLiM simulations under panmixia to confirm the theoretical trend based on the derivative approach above). In black, z_MAR_ and 95% Confidence Interval of species analyzed in section IV are plotted (see section for details).

##### II.3.2 Scaling z_MAR_ for low sampling and low census size

Let *z_pan-n_ = E[z_MAR_ | n, panmixia],* be the expected value of *z_MAR_* of a panmictic species given that we only have small sampling of *n*. Although theoretically *z_MAR_* should approach 0, with small samples it can be upwardly biased. In order to force the possible values of *z_MAR_* to range 0-1 despite small sample sizes, we can scale it as:

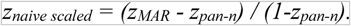

In words, this moves the purple line in Fig. S5 to zero, stretching the space above it accordingly.

Most species have census sizes so large that *z_MAR_* should indeed approach 0 under panmixia, so we should correct the sample estimate *z_MAR_* to range 0-1. However, some species have such low census size *N* that even if we sample all individuals of a species, the sample size will still be small. In those cases, we should not scale *z_MAR_* to range 0-1, but rather scale it from *z_pan-N_ − 1,* where *z_pan-N_ = E[z_MAR_ | N, panmixia]* is the expected value of *z_MAR_* given a census size *N* (plants or animals living in the wild). The updated scaling approach for both census and sample size would then be:

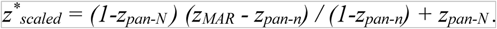

Note that this scaled estimate must be conservative because while we adjust the minimum *z* for the average value expected for low sample sizes, we do not adjust for the maximum possible *z*, which only under very extraordinary theoretical conditions can be *z=*1, namely under an unrealistic complete disconnection of populations by gene flow (see below). Because deriving the maximum *z* would require more biological knowledge of the species’ demography, landscape connectivity, genome structure, etc., and because we rather create conservative estimates, we do not create further scaling approaches.

##### II.3.3 Meta-populations in space

A more realistic simulation than a panmictic population is that of the same 40×40 deme grid where migration can happen between adjacent demes. This migration rate can be changed to understand the effect of population structure and migration on *z_MAR_*. Under no migration (or very low migration), we expect the mutations in two distinct populations (and thus their SFS) to be (almost) completely independent. Hence, when explored demes are doubled (*N_e_* doubles), we discover twice as many mutations. In this case, the number of mutations should scale linearly with the area, so we expect the following to be true: *M=A*, *log(M) = log(A),* and *z_MAR_=1*. Our analyses under different sampling schemes, and with different numbers of “burn-in generations” (generations since a single deme colonised the full 40×40 space) confirm that *z_MAR_* approaches 1 in the limit of low migration (see Table S1 and Fig. S6). Different from the panmictic situation, as we increase the sampled area, we not only increase *n*, which would lead to a *log(A)* in mutations, but also increase *N_e_*.

**Fig. S6.**
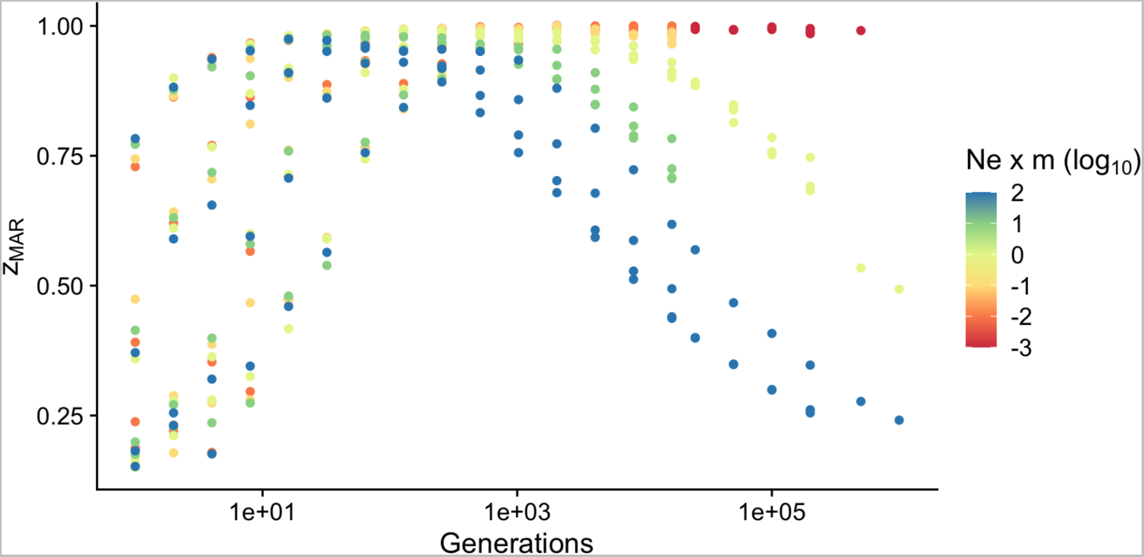
msprime 2D deme simulations and the mutations-area relationship. Simulations with different burn-in and migration rates under neutrality, and their corresponding zmar.

**Table S1.** msprime population genetic simulations in 2D. Simulations summarised by grouping ranges of the resulting _z_MAR parameters. The average parameters of the simulations with similar z_MAR_ EW provided. (Acronyms: Nemt = product of effective population size, migration rate, and simulated generations).

| $z_{MAR}$ | Samples/deme | Generations | Migration rate | $N_{emt}$ |
| --- | --- | --- | --- | --- |
| 0.2 +/- 0.05 | 2.4 | 50001.7 | 0.0271675 | 5000044.23 |
| 0.3 +/- 0.05 | 20.25 | 70003 | 0.0561655 | 7000075.77 |
| 0.4 +/- 0.05 | 26.5714286 | 13057.4286 | 0.04450857 | 1305497.96 |
| 0.5 +/- 0.05 | 12.9230769 | 121759.462 | 0.04017769 | 752221.743 |
| 0.6 +/- 0.05 | 15.6111111 | 3218.77778 | 0.045735 | 321174.768 |
| 0.7 +/- 0.05 | 35.6842105 | 35034.8421 | 0.03395895 | 143791.614 |
| 0.8 +/- 0.05 | 35.030303 | 15655.1212 | 0.03055818 | 58023.5539 |
| 0.9 +/- 0.05 | 36.5806452 | 3057.12903 | 0.0253029 | 15290.4081 |
| 1 +/- 0.05 | 42.0140845 | 13625.4085 | 0.00861178 | 1798.36141 |

These simulations corroborated that we can recover *z_MAR_* values ranging between 0-1 just varying migration and burn-in generation parameters. We found that it was both the time of the system to reach an equilibrium as well as the migration rate that determined *z_MAR_*. In the future, it will be interesting to study different non-equilibrium scenarios to better understand how genetic drift, gene flow, and different landscape structures may shape the *z_MAR_*.

##### II.3.4 Metapopulations in space with local adaptation

In order to simulate local adaptation, we use the individual-based simulation software SLiM (*24*) following the approach of (*25*). These simulations were set up for 196 demes arranged in a 14 x 14 grid. Each grid cell contains a population of *N=1000* and has an environment attribute, *e*, which varied spatially from the lower-left to the upper-right corners (approx. *-7 < e < 7*). 12 locations in the genome were allowed to be under directional natural selection. The selection coefficient was fixed for a simulation, and grid runs were conducted with *0<s<0.05*, but this selection would vary based on the environmental selection value of a grid cell, according to *e × s*. Therefore, these alleles are antagonistic pleiotropic. Selected mutations across the 12 loci in the genome behaved additively (e.g. if an individual in grid cell *i* had two of the selected mutations, fitness would be *w=1+2s × e_i_*). The migration rate varied from one individual in a billion (1*×*10^-9^), to one individual every ten (1*×*10^-1^). Finally, the mutation rate was set to 10^-8^ mutations/bp/generation and the recombination rate to 10^-7^ crossovers/bp/generation.

**Fig. S7.**
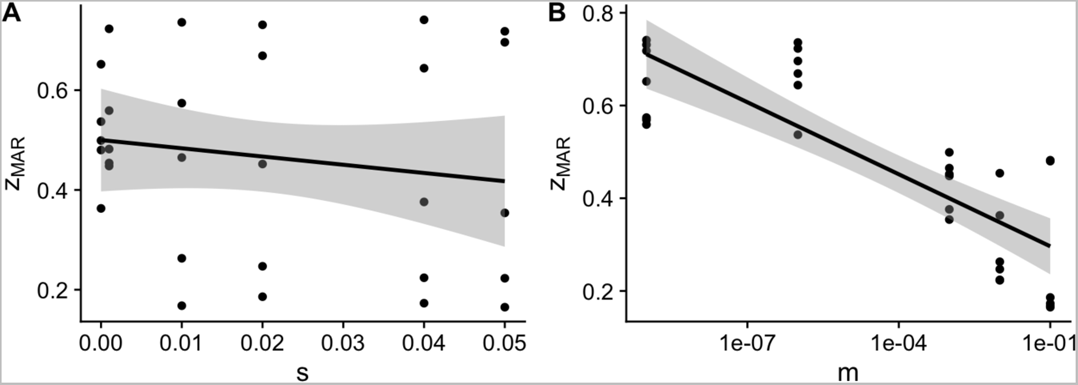
SLiM population genetic simulations in 2D with selection and local adaptation. Simulations were carried out with different combinations of migration rates and strength of antagonistic pleiotropic selection at 12 QTLs. (A) Marginal relationship between z_MAR_ with the strength of spatially-varying selection s. (B) Marginal relationship between z_MAR_ with the migration rate m.

These results, together with individual-based simulations, corroborate what we had observed with coalescent simulations, i.e. that *z_MAR_* is lowest with a high migration rate. The simulations also appear to show a negative effect of selection on *z_MAR_*. Generating a linear model fitting migration rate and selection and their interaction to understand what factors explain the scaling coefficient: *z_MAR_ ~ log_10_(m) + s + log_10_(m) s;* we confirm that both had a significant effect, and that selection significantly reduces *z_MAR_* (Fig. S7, see below summary Table S2). This may seem counterintuitive, as one may expect that locally-adaptive mutations are rare and will be localised only to where they are adaptive. More work is necessary to understand the signatures that spatially-varying natural selection (and its different types) create on *z_MAR_*, but we can think that under migration limited scenarios (where *z* approaches1) adaptive alleles and their linked mutations permeate faster to similar neighbour environments than neutral alleles.

**Table S2.** Linear model explaining z_MAR_ by migration rate and natural selection Summary table of the linear model z_MAR_ ~ mig + s + mig:s.

|  | Estimate | SE | t-value | P-value |
| --- | --- | --- | --- | --- |
| intercept | 0.3385022 | 0.0469174 | 7.214859 | 0.0000001 |
| mig | -0.0419733 | 0.0085804 | -4.891792 | 0.0000407 |
| s | -4.693492 | 1.6290184 | -2.881178 | 0.0076725 |
| mig : s | -0.4998393 | 0.2426463 | -2.059950 | 0.0491621 |

##### II.3.5. Metapopulations in space with purifying selection

To understand the effect of purifying selection on *z_MAR_* we also ran 2D simulations with a fraction of the genome allowed to be globally-deleterious (i.e. independent of the spatially-varying environment). We simulated an increasingly strong purifying selection (|*s|* range from 0.0 to 0.1), simulating roughly that 29% of the genome of Arabidopsis is coding (arabidopsis.org) and mutations can be deleterious. We also varied the degree of recombination. Following our expectation, with stronger purifying selection deleterious mutations are pushed to lower allele frequencies, stopping their geographic spread, which increases *z_MAR_*. Recombination rate appears to have a minor role on *z_MAR_* (Fig. S8).

**Fig. S8.**
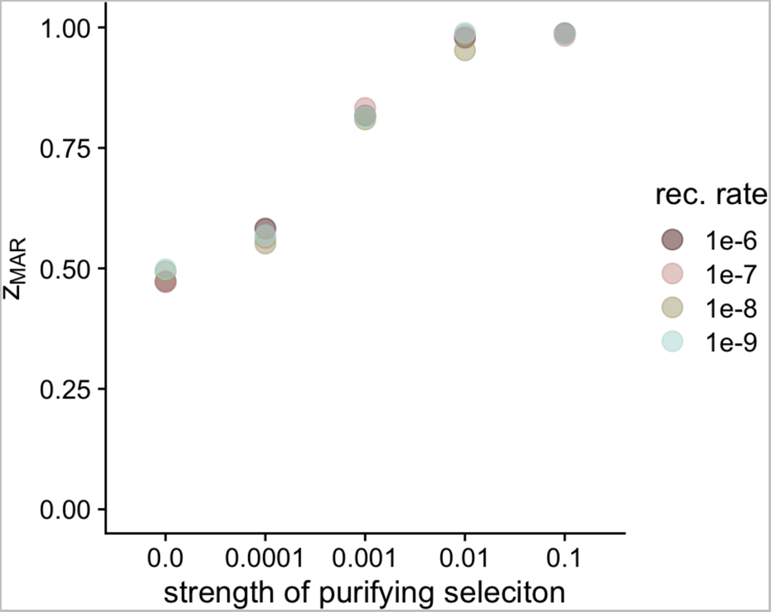
SLiM population genetic simulations in 2D with purifying selection. Simulations were carried out with varying strengths of purifying selection (|s| range from 0.0 to 0.1) at coding positions, representing about 29% of the genome. Different values of recombination rate were also used in all pairwise combinations with |s|.

##### II.3.6 Continuous-space non-Wright-Fisher models

In order to confirm *z_MAR_* generality in highly realistic conditions and its behavior through the population extinction process (II.4), we set up SLiM simulations in continuous space using non-Wright-Fisher dynamics (*24*). Spatial population structure in these simulations was established through individual dispersal, local mate choice and spatial competition, which we chose to lead to realistic values of *F_ST_* across space. Spatial competition also acted as population control, by keeping the total population size below a target carrying capacity through direct effects on individual fitness. In addition to competition, fitness was also affected by individual age as well as by a polygenic trait under stabilising selection. A subset of variants (final proportion ~10%) directly affected this trait with effect sizes drawn from a Gaussian distribution with mean = 0.0 and standard deviation = 0.1, and a fitness penalty was incurred by deviating from the optimal trait value using a Gaussian fitness function centered at the optimum and with a standard deviation = 5.0. We initialised functional variation for SLiM using neutral coalescent simulations with msprime (*23*) to reduce the computational burden of burn-in, and loaded the resulting tree sequences into SLiM (*26, 27*). We drew functional effect sizes for these variants, placed individuals into continuous space, and ran simulations forward-in-time for 5,000 generations. After that, the geographic distribution of the species experienced impacts as expected during global change: every generation, 0.001 of one edge of the species distribution got its carrying capacity reduced to 0. This meant that over 1,000 generations the whole species would disappear (note that this is a reasonable fraction of area reduction given the estimates of yearly deforestation and habitat change in section V). We subsequently overlayed neutral mutations on the tree sequence using msprime, and analysed genomes sampled throughout the extinction process (by tracking them in the tree sequence output) and extracted using tskit.

**Fig. S9.**
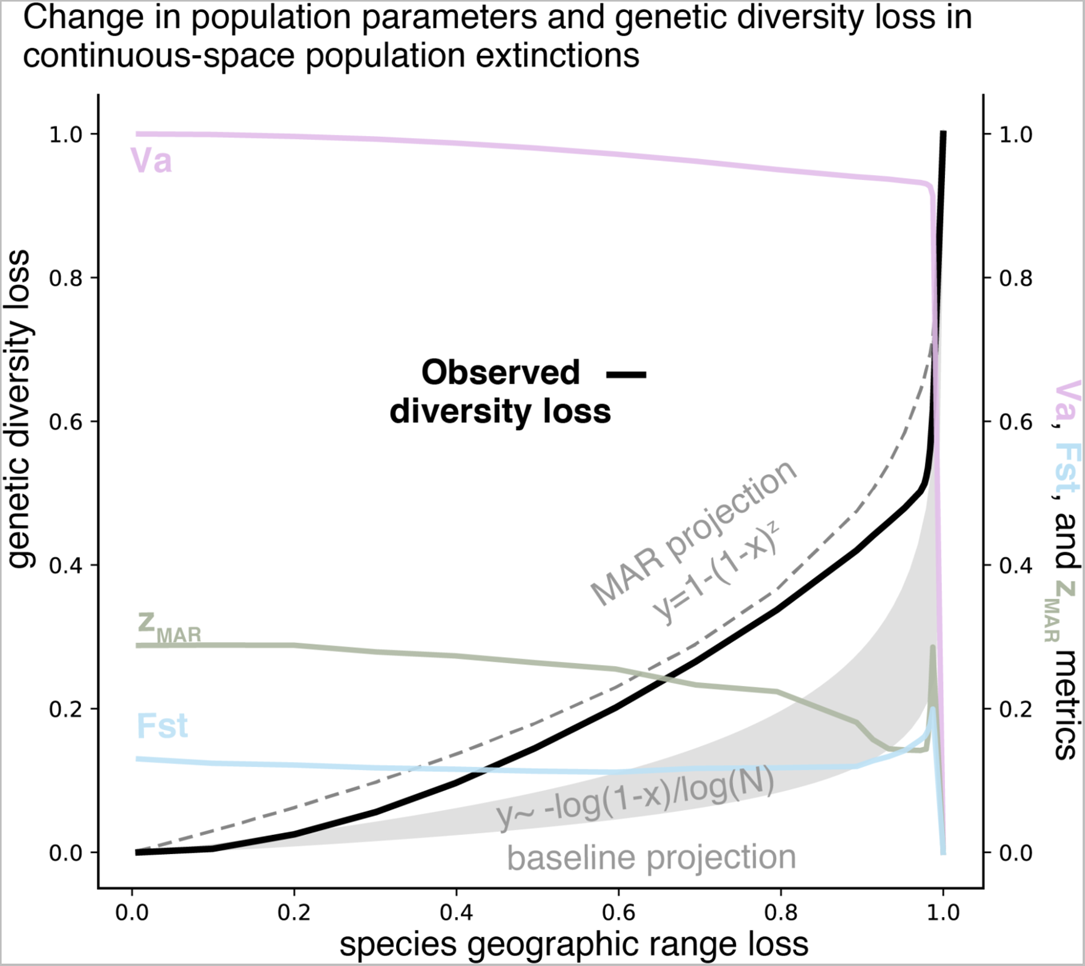
Continuous space SLiM population genetic simulations. At 19 timepoints leading up to extinction, 1,000 individuals were sampled randomly in continuous space to quantify diversity loss (black line). The prediction of MAR (dashed line) using the starting z_MAR_ seemed to follow the real trend better than the baseline of just loss of individuals(dashed line). This suggests that even if z_MAR_ varies during the population extinction process, it is relevant to understand genetic loss by area reduction. We also tracked metrics of population structure (z_MAR_, F_ST_) and a proxy of adaptive capacity (Va), which showed qualitatively similar patterns as the GWA-based trends (Fig S21).

##### II.3.7 Connection of *z_MAR_* with the isolation-by-distance pattern

Ultimately, *z_MAR_* is a complex integrator of evolutionary forces acting in space (mutation, migration, drift, selection) and captures how structured the distribution of a species’ mutations is. Although the isolation-by-distance pattern conceptually resembles *z_MAR_*, we have found no obvious analytical expression that relates both. Note that *F_ST_* is defined based on heterozygosity or π, instead of the number of segregating sites (i.e., mutations *M*). For instance, using Hudson’s estimator (*28*) to compute *F_ST_* across a set of populations we calculate *F_ST_ = 1-(π_w_ / π_b_)*, where *π_w_* is the diversity or heterozygosity within a population and *π_b_* is the same parameter calculated for the meta-population. Plotting *F_ST_* of a metapopulation by the distance of the farthest demes shows the typical non-linear trend of isolation-by-distance, which shows that very close populations have similar allele frequencies whereas populations further away drift apart. A challenge of *F_ST_* is that it requires pre-defining discrete populations, which is straightforward in stepping-stone simulations but hard in real data. Comparing average *F_ST_* of our 14×14 spatial demes and *z_MAR_*, we see that the two parameters correlate (Fig. S10C). However, it appears that for low values of *F_ST_*, *z_MAR_* captures more variation across the simulations (Fig. S10). These patterns were also confirmed in continuous space simulations (not shown).

**Fig. S10.**
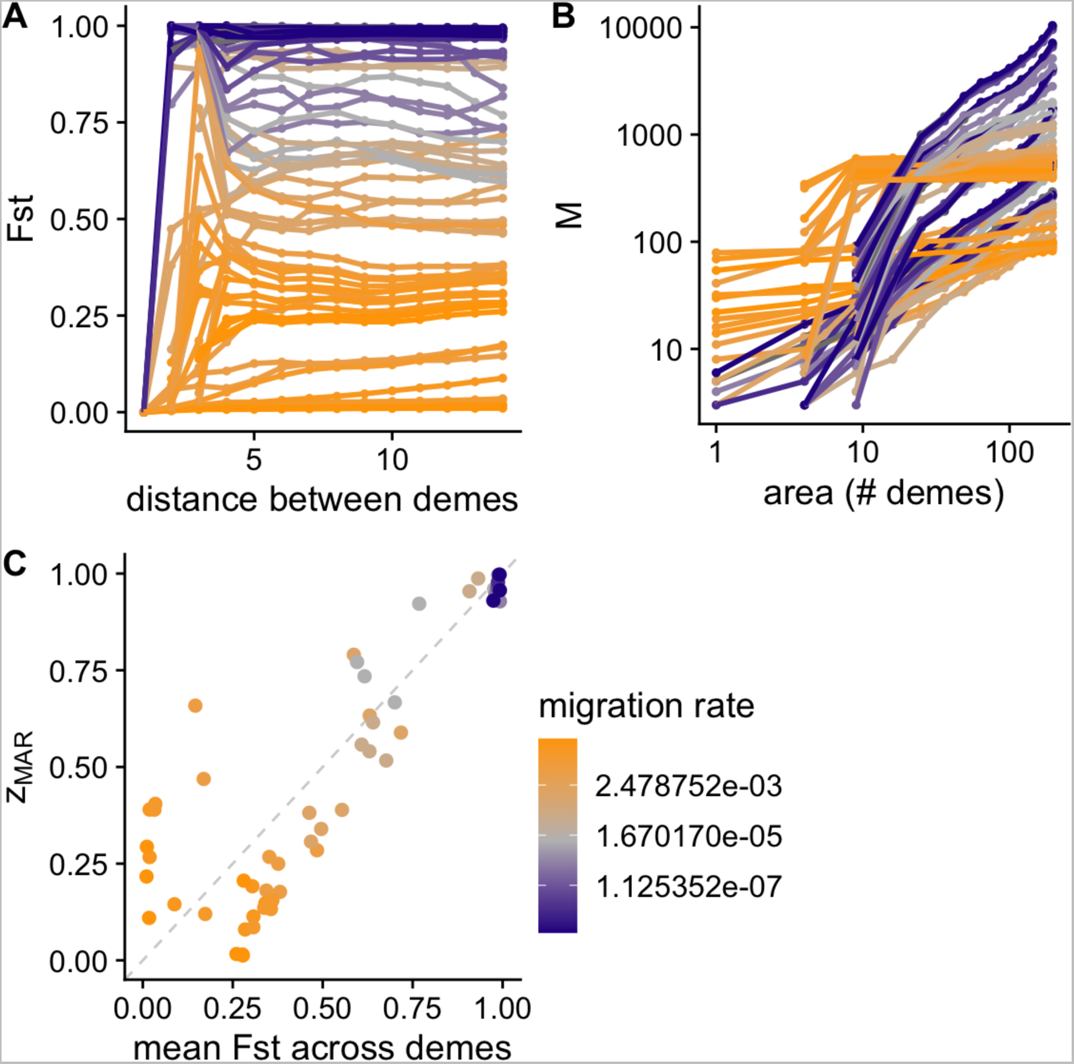
SLiM population genetic simulations in 2D comparing F_ST_ and z_MAR_. Neutral SLiM simulations with different degrees of migration. (A) Hudson’s F_ST_ across populations with different area subsamples. Following the expectation of the isolation-by-distance pattern, as the distance between the farthest demes in the subsample increases, F_ST_ becomes larger and saturates at large distances. (B) The mutations-area relationship. (C) Comparison between F_ST_ and z_MAR_.

##### II.4 The loss of mutations (genetic diversity) in space

The aim is to predict the fraction of genetic diversity loss, *x_M_*, from shrinking of an ecosystem by an area *a.* To define all terms, we then have a past area *A_t-1_* and a present reduced area *A_t_=A_t-1_* − *a*, and a fraction of area extinct *x=a/A_t-1_*

We first think of the loss of genetic diversity *x_M_* through the basic process of losing individuals. From the population genetics’s coalescent theory derivation of the number of mutations or segregating sites from individuals we got the approximation *M~log(N).* Assuming the loss of area is simply the loss of individuals (*A=N*), we can derive the fraction of genetic diversity loss as:

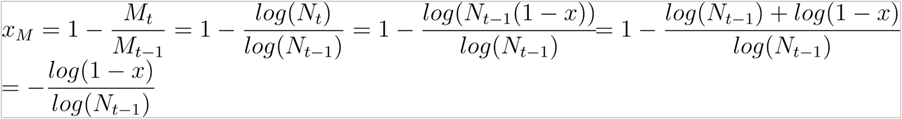

The loss of mutations is then in the scale of: *log(1-x)*; which is very slow, as we expected from having derived the trend that under panmixia *z_MAR_* ≈ 0. A substantial loss of genetic diversity in this case only happens when population extinction is almost complete.

Species do not typically behave perfectly panmictic given different *z_MAR_* values. Under population structure, we can use our relationship to project the number of mutations (genetic diversity) lost as the geographic distribution due to habitat loss or climate change following equation:

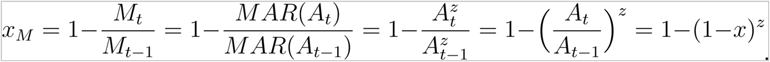

In the most extreme scenario of *z_MAR_* ≈ 1, the fraction loss of geographic area directly translates to the same fraction loss of genetic diversity.

Reality should be in between the panmictic and fully-migration-limited cases. With combinations of environmental selection, non-equilibrium demography, and long-range dispersal, we may get intermediate *z_MAR_* values, and it will be empirical estimates that can inform us how much may be lost (Section III).

##### II.5 Recovery of genetic diversity after a bottleneck or local extinction

**Fig. S11.**
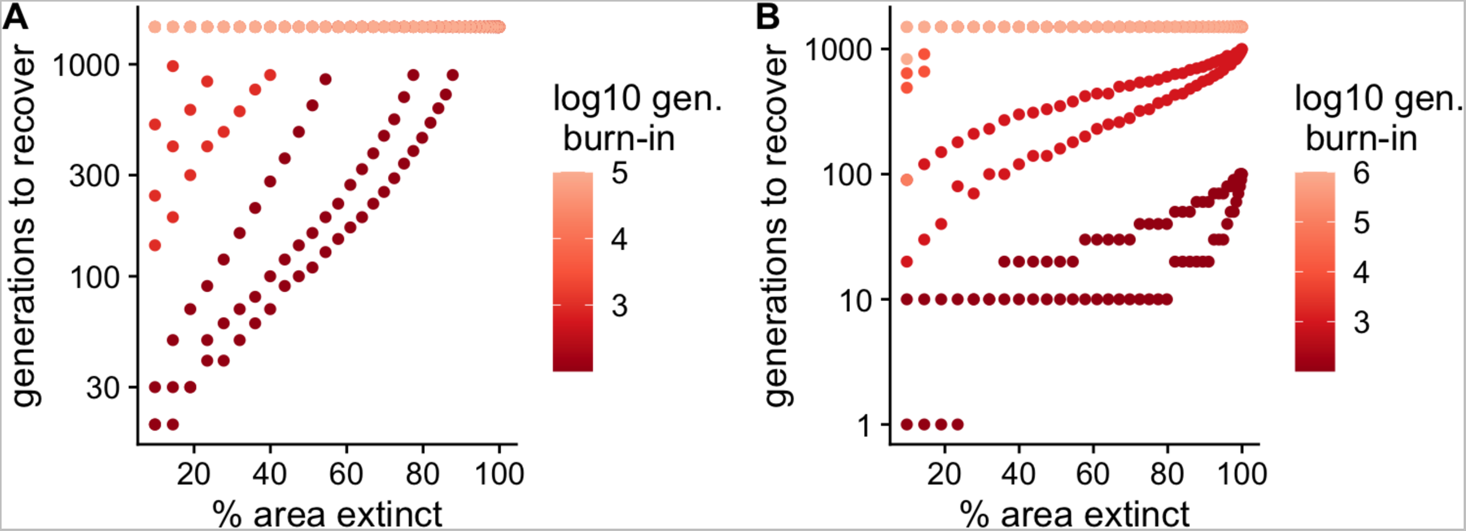
2D stepping-stone msprime simulations with extinction and recovery. (A) Recovery of genetic diversity (number mutations) after loss of a fraction of the population. (B) Recovery of genetic diversity after instantaneous loss of a fraction of the population and consecutive repopulation. *Simulations with number of generations until recovery that are exceedingly large are assigned a value of 1,500, as none are realistic for current conservation timelines.

The intuition that rapid recovery of genetic diversity may be possible is likely flawed. While genetic recovery may be faster than speciation rates, which are on the order of millions of years, the time for a set of populations that went through a simulation burn-in of 1,000 generations (not yet in diversity equilibrium), and that suffer an instantaneous 5% reduction of area and an instantaneous recovery (e.g., through reforestation) would range from 20-90 generations. This number of generations for long-lived species would translate into centuries or millennia of recovery without further impacts. About 49% of simulations – including every simulation that reached equilibrium (burn-in generations >10,000) – have a recovery time of more than a thousand generations (Fig. S11).

### SUPPLEMENTAL RESULTS

#### III. The mutations-area relationship with the 1001 Arabidopsis Genomes

We begin testing the idea of a general mutations-area relationship using the extensive sampling of the model plant species *Arabidopsis thaliana* and the 1001 Arabidopsis Genomes Project (*29*). This section will serve as a case study to explore different approaches and biases when building MAR to then apply the learned lessons across species (section IV).

##### III.1 The Site Frequency Spectrum of the 1001 Arabidopsis Genomes

We began analyzing the frequency distribution of 11,769,920 biallelic genetic variants (i.e., mutations), which is typically called the Site Frequency Spectrum (SFS) in population genetics.

**Fig. S12.**
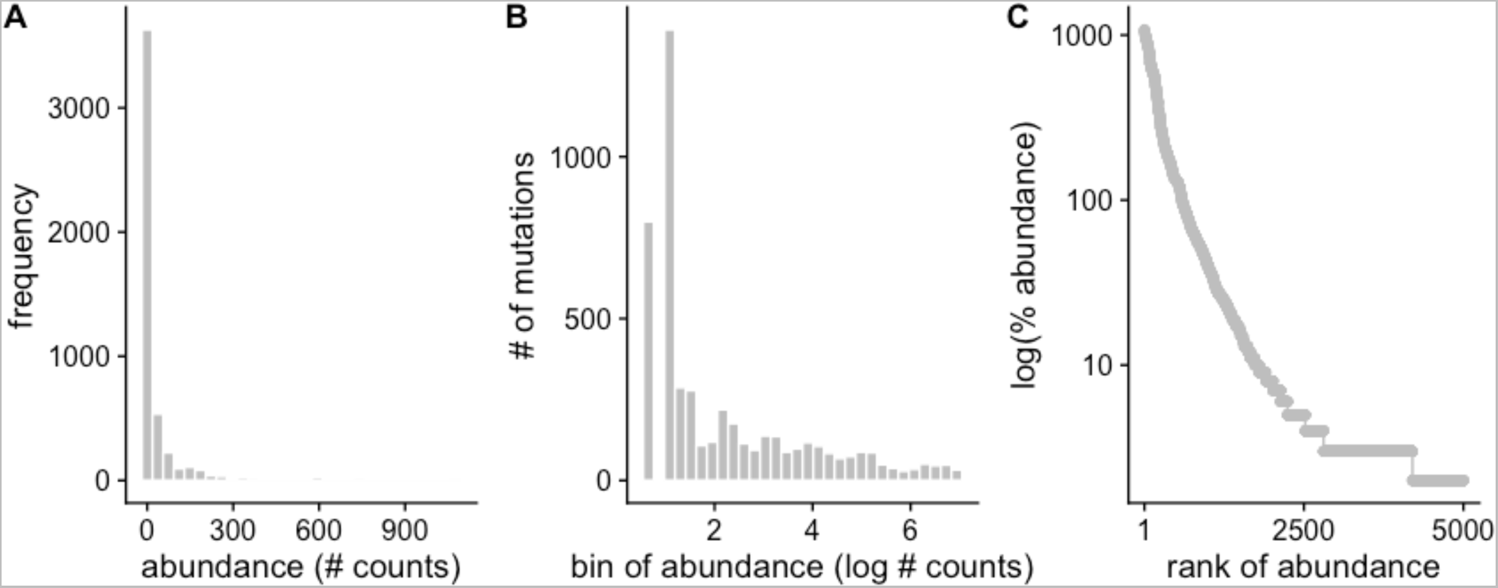
Mutation abundance study in *A. thaliana*. (A) Site Frequency Spectrum (SFS). (B) Preston plot of mutation abundances. (C) Whittaker plot of mutation rank abundances.

To showcase the similarities to the Species Abundance Distributions (SAD), we use the Whittaker plot of mutation rank abundance (Fig. S12) that suggests a log-normal of S-shape may be the best fitting model (Table S3). For a review listing many popular models, see (*30*), and for implementation details of 13 SAD models see the thorough manual of R package SADS (*31*). As we shall see later, the log-normal distribution seems to be the best fit across species.

**Fig. S13.**
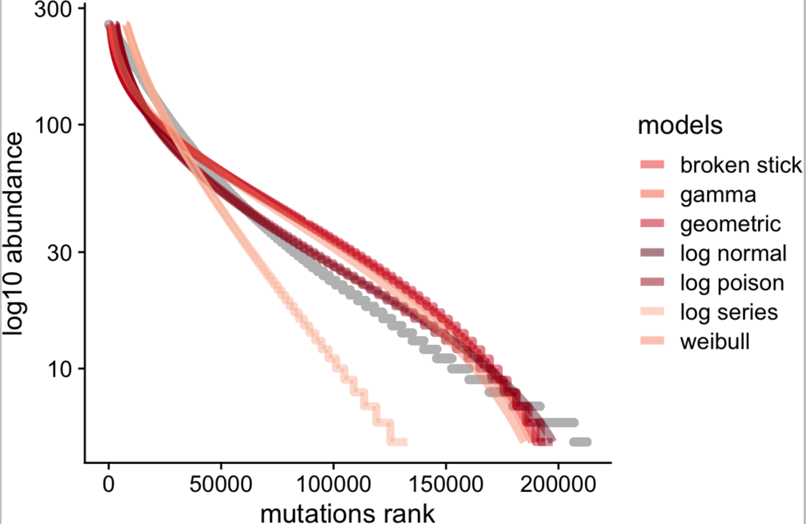
Fit of mutation abundance study in *A. thaliana* with different SAD models. Representative models from Table S3 are plotted along with the observed frequency of 11,769,920 mutations.

Although model AIC captures best the fit of a curve accounting for the difference in parameter complexity of each model and the statistical distributions behind, we often are interested in the variance explained. We then calculated a proxy of predictive accuracy using a pseudo*-R^2^* approach of the difference between the model fit and the observed data as: 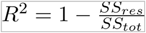. For *A. thaliana*, we used 10,000 SNPs sampled at random to an accuracy of over *R^2^* >0.999 for both the top log-Normal model and the bottom log-Series model, indicating that all “commonness of rarity” models must have a pretty good fit of mutation frequency data.

**Table S3.**
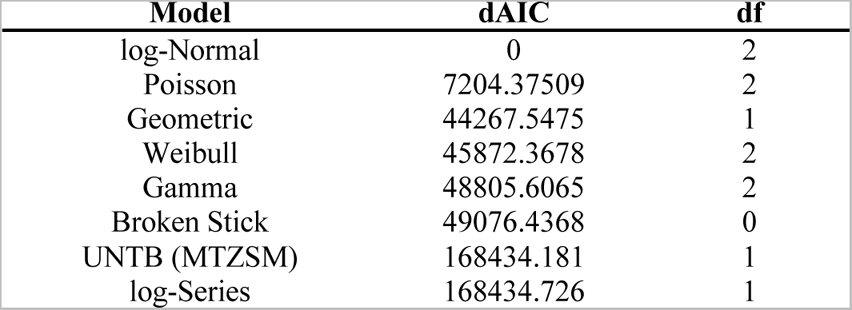
AIC values for model fit of common species distribution curves. For each SAD model, the degrees of freedom and the delta AIC compared to the top model are reported.

The typical SFS from population genetics is of course not implemented in current packages for Species Abundance Distributions like R sads. For comparison, in the main text we also calculate the log likelihood and AIC of this following the standard population genetics likelihood:

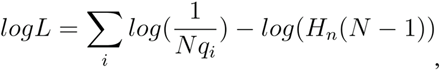

where *N* represents the number of individuals in a sample, and *q_i_* is the minor allele frequency of a SNP in the sample, in the main text calculated for *i*=1…10000 random SNPs (see main text). As before, *H_n_* is the harmonic number function.

##### III.2 Building the Mutations-Area Relationship

In the following, we explain how the area was estimated that was used to compute *z_MAR_* on real world data. In short, we used a grid on the world map, with samples placed on the map based on their geo-coordinates of origin (Fig. 1). We first create square spatial subsamples of the *Arabidopsis thaliana* geographic distribution (Fig. 1, Fig. S15) and quantify diversity *M* as the total segregating sites. Excluding zeros, these two variables are fed to the sars_power function from the R SARS package (*32*).

Although the power law mutations-area relationship was already theoretically motivated (II.3), here we also fit different types of functions typically applied to the Species-Area Relationship. Doing this, we reach the conclusion that multiple models perform very similarly, and the classic power law is among the top models, see Table S4. Although small marginal fitting accuracy could be achieved with other models, for mathematical convenience and historical continuity, we use the power law for later sections and the study of MAR across species (Sections IV and V).

**Table S4.**
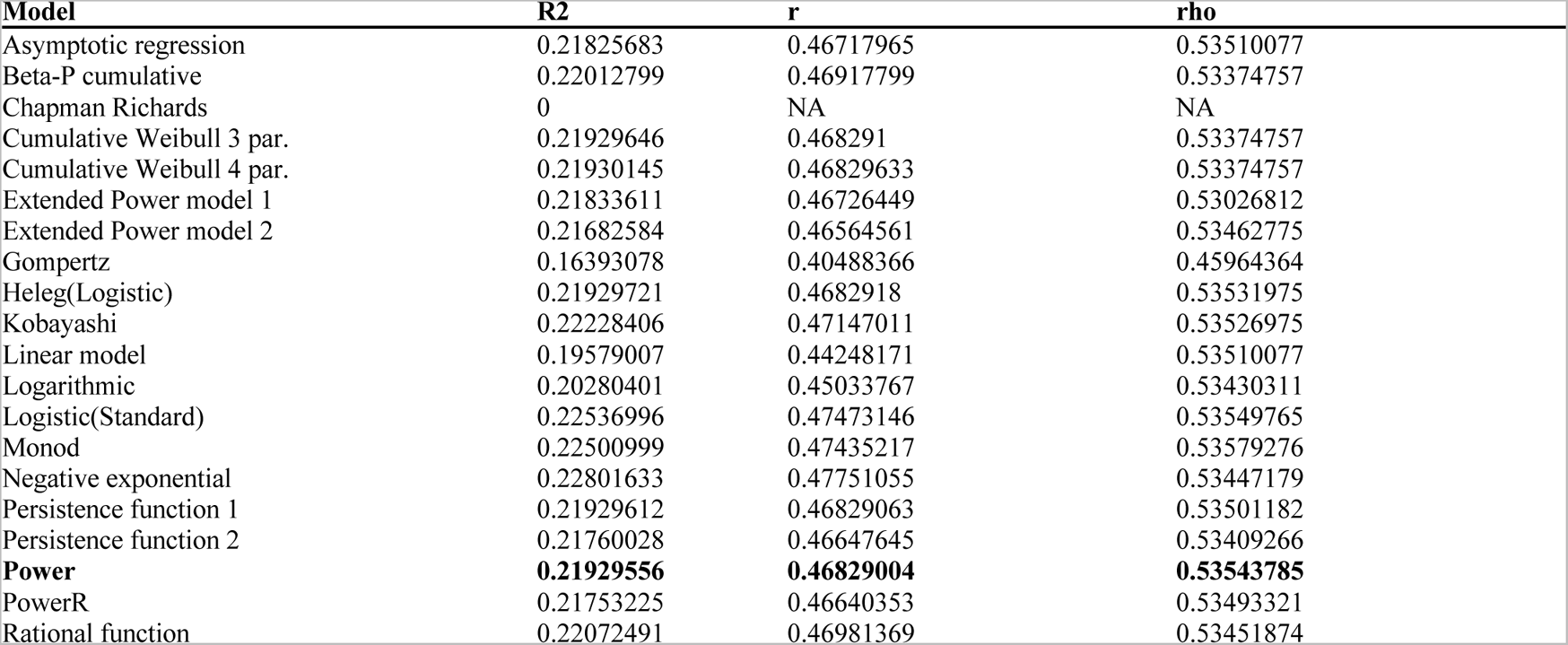
Different SAR curves fit to mutations. We fit 20 different functions and calculated the variance explained (R2), Pearson’s r, and Spearman’s rho.

| Model | $R^2$ | $r$ | $\rho$ |
| --- | --- | --- | --- |
| Asymptotic regression | 0.21825683 | 0.46717965 | 0.53510077 |
| Beta-P cumulative | 0.22012799 | 0.46917799 | 0.53374757 |
| Chapman Richards | 0 | NA | NA |
| Cumulative Weibull 3 par. | 0.21929646 | 0.468291 | 0.53374757 |
| Cumulative Weibull 4 par. | 0.21930145 | 0.46829633 | 0.53374757 |
| Extended Power model 1 | 0.21833611 | 0.46726449 | 0.53026812 |
| Extended Power model 2 | 0.21682584 | 0.46564561 | 0.53462775 |
| Gompertz | 0.16393078 | 0.40488366 | 0.45964364 |
| Heleg(Logistic) | 0.21929721 | 0.4682918 | 0.53531975 |
| Kobayashi | 0.22228406 | 0.47147011 | 0.53526975 |
| Linear model | 0.19579007 | 0.44248171 | 0.53510077 |
| Logarithmic | 0.20280401 | 0.45033767 | 0.53430311 |
| Logistic(Standard) | 0.22536996 | 0.47473146 | 0.53549765 |
| Monod | 0.22500999 | 0.47435217 | 0.53579276 |
| Negative exponential | 0.22801633 | 0.47751055 | 0.53447179 |
| Persistence function 1 | 0.21929612 | 0.46829063 | 0.53501182 |
| Persistence function 2 | 0.21760028 | 0.46647645 | 0.53409266 |
| **Power** | **0.21929556** | **0.46829004** | **0.53543785** |
| PowerR | 0.21753225 | 0.46640353 | 0.53493321 |
| Rational function | 0.22072491 | 0.46981369 | 0.53451874 |

Because in the species literature it is recommended to only quantify richness of endemic species (*33*), we also count segregating sites that are private to the area subsample, creating the equivalent endemic-mutations-area relationship (EMAR) (*33*). The MAR slope and 95% Confidence Interval was *z* = 0.324 (0.238 - 0.41) (Table S5, Fig. S14 A), while the EMAR was *z* = 1.241 (1.208 - 1.274) (Table S6, Fig. S14 B). Interestingly, the endemics-area relationship of *z* ≈ 1 resembles that of endemic species, whereas the total mutation relationship with area is above that of species relationships, which typically follows the canonical *z* ≈ 0.2 − 0.4.

We must note that EMAR, the genetic analogy of the Endemic-(species)-Area Relationship (EAR) may not be that meaningful when analyzing genomic data (we did not find a way to theoretically motivate it in section II), and later we see it overestimates loss in our simulations (Fig. S18)

**Table S5.** The mutations-area relationship (MAR). Fitted values in a log-log power function between area sampled and mutations discovered.

|  | Estimate | Std. Error | t value | P | 2.5% | 97.5% | nls.Est. | nls.2.5% | nls.97.5% |
| --- | --- | --- | --- | --- | --- | --- | --- | --- | --- |
| c | 494.565432 | 135.6314588 | 3.646392 | 0.0003138 | 223.3025141 | 765.8283493 | 494.5531270 | 278.1107276 | 822.829918 |
| z | 0.323727 | 0.0430277 | 7.523681 | 0.0000000 | 0.2376715 | 0.4097824 | 0.3237367 | 0.2430303 | 0.413162 |

**Table S6.** The endemic-mutations-area relationship (EMAR). Fitted values in a log-log power function between area sampled and endemic mutations discovered.

|  | Estimate | Std. Error | t value | P | 2.5% | 97.5% | nls.Est. | nls.2.5% | nls.97.5% |
| --- | --- | --- | --- | --- | --- | --- | --- | --- | --- |
| c | 0.0001001 | 0.0000231 | 4.337758 | 1.98e-05 | 0.0000539 | 0.0001463 | 0.0001001 | 0.0000635 | 0.0001555 |
| z | 1.2411831 | 0.0165268 | 75.101442 | 0.00e+00 | 1.2081296 | 1.2742366 | 1.2412125 | 1.2096087 | 1.2737927 |

**Fig. S14.**
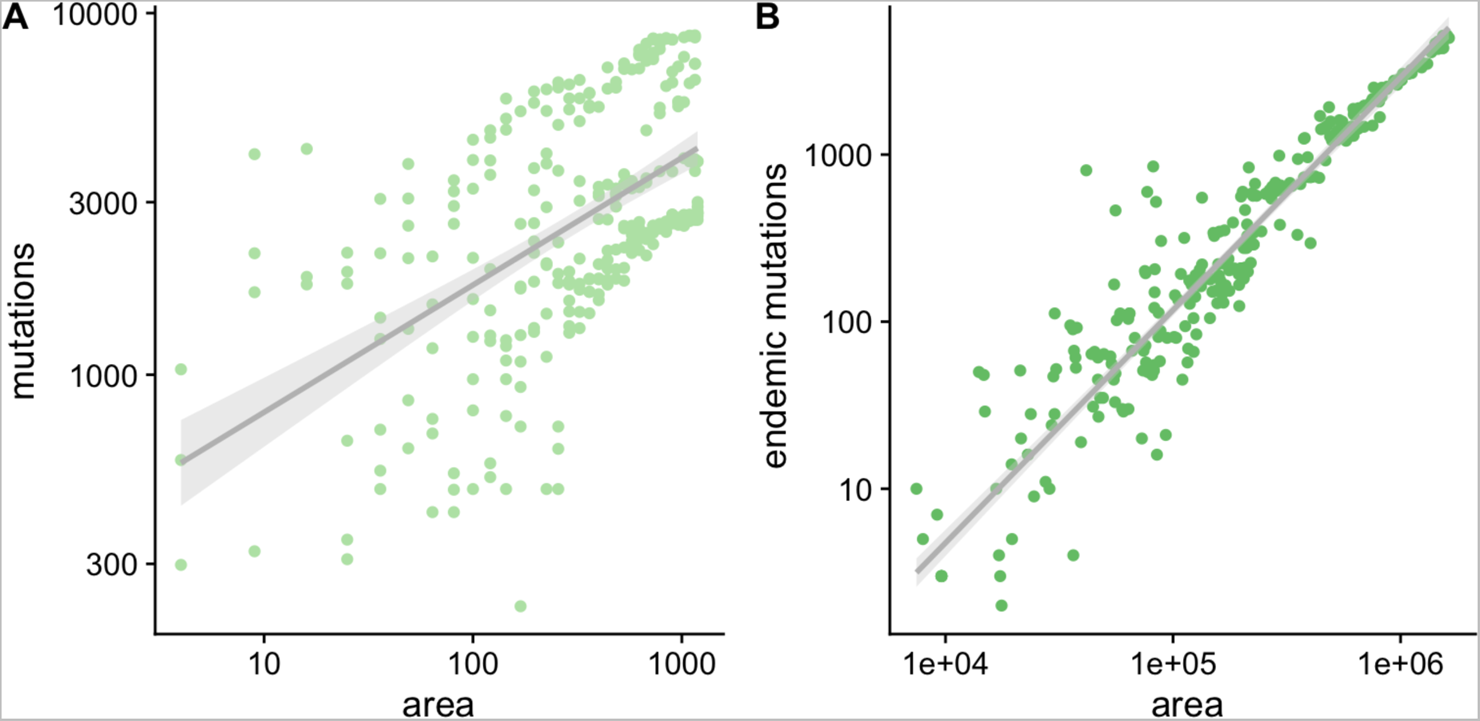
The mutations-area and endemic-mutations-area relationships in *A. thaliana*. Dividing *A. thaliana* native geographic distribution into a 1 degree lat/long grid, square areas with 1 degree side-length to 36 degrees side-length were randomly placed (n=100 for each size) across the distribution, and genetic diversity metrics were computed to produce the (A) Mutations-Area Relationship and (B) Endemic-Mutations Area relationship.

##### III.3 Testing for potential numerical artefacts

We wondered whether MAR estimates may be affected by some numerical artefacts in our software pipeline (available at https://github.com/moiexpositoalonsolab/mar). For instance, real world data may have uneven sampling in space, the spatial resolution of georeferenced samples may vary, projection of samples into gridded maps may have limited resolution, software pipelines may produce biased estimates, etc. To test this, we conducted several experiments:

###### Lower bound of the method for *z_MAR_*

Our first experiment when building the MAR aimed to make sure that spatial sampling, or some unknown bias in genome sequencing, or the number of samples used, are not creating artificially large *z_MAR_*. We then simulated a mock dataset of *A. thaliana* with the same number of mutations, samples, and using the original geographic locations. The number of SNPs were also sampled in a way that we created a canonical 1/q SFS for the whole species. Under no biases, we then expect the MAR to follow the theoretical derivation under panmixia with a *z*~0. This exercise confirmed we get a value approaching zero: *z=*0.033, (−0.095 - 0.162).

**Table S7.** MAR built with different area calculations and grid sizes.

| Grid resolution<br>(deg.) | $z_{MAR}$ [CI95%]<br>(cell area) | $z_{MAR}$ [CI95%]<br>(total area) |
| --- | --- | --- |
| A=N | 0.431 (0.423 - 0.439) | NA |
| 0.1 | 0.435 (0.424 - 0.446) | 0.367 (0.281 - 0.454) |
| 0.25 | 0.454 (0.449 - 0.459) | 0.422 (0.376 - 0.467) |
| 0.5 | 0.488 (0.465 - 0.511) | 0.352 (0.152 - 0.551) |
| 1 | 0.543 (0.529 - 0.558) | 0.389 (0.295 - 0.483) |
| 2.5 | 0.644 (0.6 - 0.688) | 0.388 (0.251 - 0.526) |
| 5 | 0.617 (0.205 - 1.029) | 0.403 (-0.204 - 1.011) |

###### Grid sizes, area calculations, and non-random spatial sampling

In order to streamline geospatial operations, we implemented the MAR relationship calculations in this project using R raster objects (*34*). This required projecting the collected samples of a species and the observations of any given mutation into a world map (i.e., each mutation’s geographic distribution). Necessarily, in order to be able to assign areas to sets of samples or mutations on the map, the projection requires the choice of a grid size. The larger the grid size (e.g., lower spatial resolution), the faster the spatial operations can be performed. Further, for larger grid sizes, we expect the slope of MAR to be more influenced by larger-scale patterns, while for smaller grid sizes, the MAR will be influenced by smaller-scale patterns. To test this, we repeated the subsampling of *A. thaliana* distribution with grid sizes ranging 0.1 degrees latitude/longitude (roughly 10km side-length in temperate regions) to 10 degrees (roughly 1,000 km side-length). The estimates were roughly consistent between 0.4-0.6, but increases in value at larger grid sizes (row in Table S7 for large grid size values), a scale-dependent pattern that resembles results of SAR of species in ecosystems fitted at different scales (*10*).

Because we often have sparse samples of individuals in space, we devised two strategies to calculate areas during the subsampling of MAR (see cartoon in Fig. S15): (A) the total square area of the minimum and maximum latitude/longitude values of all the samples analyzed. That is, simply the area of the red box in the figure. (B) the sum of areas of grid cells that contain at least one sample. That is, the sum of the grey squares within the red box in the figure. In addition, we also calculated the MAR relationship assuming the total area is equal to the number of individuals (*A=N*) (which should be theoretically equivalent to a grid of very high resolution where we end up with a maximum of one individual sampled at any grid cell).

Table S7 values suggest there is a dependency of *z_MAR_* with the grid size when areas are calculated as the sum of grid cells with at least one sample. Our intuition for this pattern is that lower resolution grids (e.g., 5 degrees side) lead to some grid cells having many samples, which would increase the number of mutations discovered when discovering the area. On the other hand, the calculation of *z_MAR_* using the total area does not seem to affect the *z_MAR_* estimate; however, because large areas often do not have samples (limiting the potential to find new mutations), it creates a higher variance in the estimate of *z_MAR_* (see confidence intervals in Table S7 and Fig. S16). Here, we favored consistency of *z* at the expense of broader, more conservative confidence intervals. All the estimates reported below and in the main text therefore use the total area approach.

**Fig. S15.**
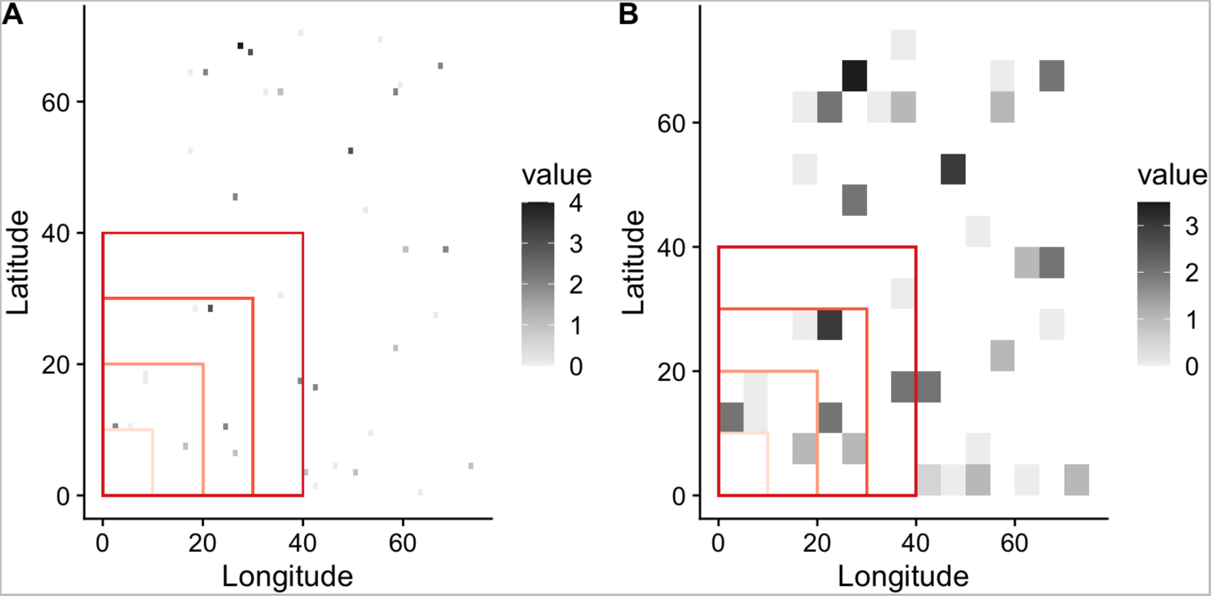
Cartoon of raster sampling to build the MAR. Map of mock samples of a species projected into a raster. Grey scale indicates the number of samples per grid cell. Red boxes exemplify the process of spatial subsampling of increasing area to build the MAR relationship. Two example grid sizes were created for illustrative purposes: (A) Small grid size or high spatial resolution. (B) Large grid size or low spatial resolution.

**Fig. S16.**
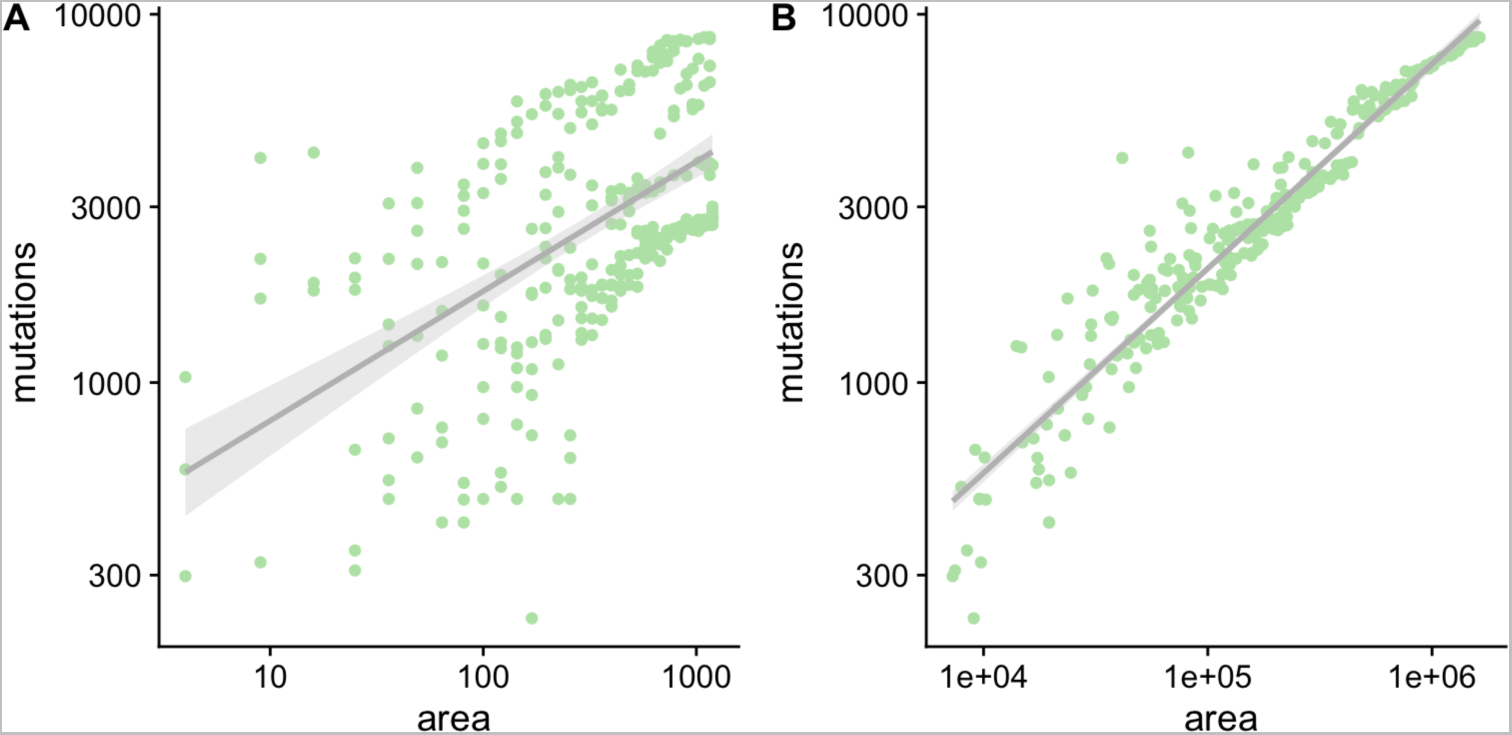
MAR comparison with different area calculations. (A) Using total area, (B) using grid cell sum with at least one sample. For 1 degree latitude/longitude grid cell.

###### Geographic subsampling strategy (inwards, outwards, random)

It has been indicated that the way the Species-Area Relationship (SAR) and Endemics-Area Relationship (EAR) are created may create differences in the scaling parameter *z*. The plots and estimates above were produced by randomly placing boxes of different size or area across the distribution of the species. Often, however, either discovery of species or extinction happen in certain patterns. For instance, we often imagine sampling an ecosystem concentrically outwards from a focal point, whereas we may think of the extinction process of species area reductions being concentrically inwards (*33*). Because these patterns seem of importance, we also calculated the MAR and EMAR outwards from the latitude and longitude median of all the samples in the map, moving outwardly until the map is filled (Fig. S17, Table S8). Likewise, the inward pattern is conducted in an inverse manner.

**Fig. S17.**
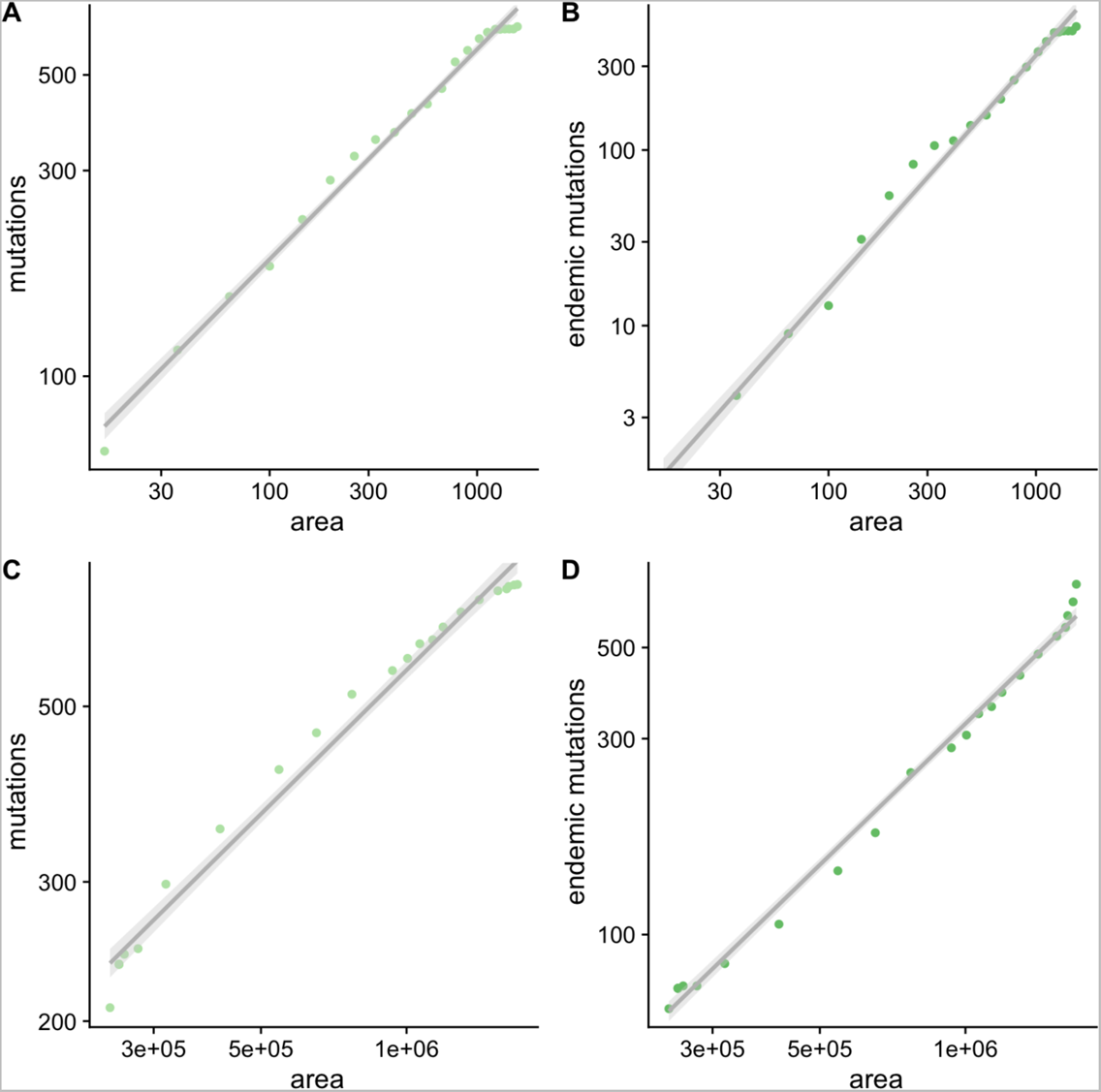
MAR and EMAR in Arabidopsis thaliana using outward and inward sampling. Dividing *A. thaliana* native distribution in 1 degree lat/long grid, a square area of 1 degree was placed at the median of the sampling range and was expanded iteratively by 1 degree lat/long until all the area of the distribution was covered. (A-B) MAR and EMAR using a typical outward sampling. (C-D) MAR and EMAR using an inward sampling. The latter may not be a common process of sample collection, but it is common for extinction progress.

**Table S8.** Outward and inward MAR and EMAR. The MAR and EMAR relationship computed with inward or outward nested subsampling, calculating area only as those cells with samples.

| Relationship | z |
| --- | --- |
| MAR outwards | 0.444 (0.412 - 0.476) |
| EMAR outwards | 1.086 (0.982 - 1.189) |
| MAR inwards | 0.561 (0.524 - 0.597) |
| EMAR inwards | 1.295 (1.192 - 1.399) |

###### Incomplete sampling of the species

To check whether the relationship holds with few individuals of a species or limited geographic distributions, we compared the species-wide MAR with that of subset populations. Downsampling the native distribution of *A. thaliana* to a region within North-East Spain (−2.00–4.25 degrees East, 36.52–42.97 degrees North), or to a region within Germany (2.69–13.73 degrees East, 50.0–52.0 degrees North), and using only 1,000 SNPs, we recovered *z_MAR_*= 0.423(0.233-0.614) for Spain and 0.525(0.242-0.807) for Germany, which were close to the estimate based on the whole distribution (Table 1). This result is reassuring in that if migratory patterns are relatively homogeneous, one may be able to estimate this parameter from a subset of the species distribution. For heterogeneous population structure cases, we expect incomplete sampling to produce unreliable estimates.

###### Number of genome-wide SNPs used

To check whether different numbers of SNPs used for the analyses would lead to different *z_MAR_*, we conducted analyses with random subsets consisting of 100, 1,000, and 10,000 SNPs, replicated 3 times. Estimates had a coefficient of variation of 4.7%, which is way below the standard error of typical estimates (Table 1).

###### Locally-adaptive variants

We then aimed to understand the effect of utilizing SNPs that appear to be related to adaptation. To study this, we utilized an outdoor climate-manipulated experiment that recorded fitness data (survivorship and reproduction output of seeds) for 515 *Arabidopsis thaliana* ecotypes part of the 1001 Genomes set in 8 environments (Exposito-Alonso, 2019). We devised two sets of alleles: 10,000 that were negatively correlated with fitness in a Genome-Wide Association across 8 different environments, and 10,000 alleles that were associated positively with fitness in one environment but negatively in another (antagonistic pleiotropic). The MAR relationship was computed as before and compared to the original random (putatively neutral) set of alleles from the previous sections (Table S9). Although we see a trend that locally-adaptive alleles have a slightly higher *z*, estimates overlap. The effects seen here of having smaller *z* for adaptive alleles than neutral variation could, however, be due to top GWA SNPs often being ascertained to higher frequency than background SNPS.

**Table S9.**
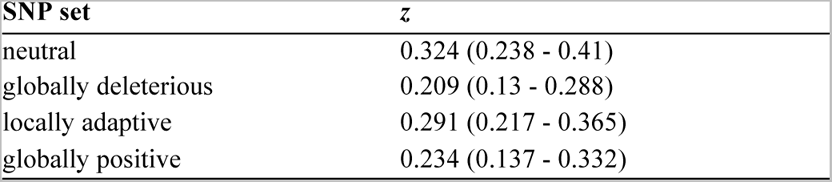
MAR for putatively neutral, deleterious, and locally adaptive alleles in Arabidopsis thaliana.

##### III.4 Local population extinction in Arabidopsis

Using the MAR framework, we can make projections of loss of mutations (or its inverse, the remaining genetic diversity. By doing this, the known intuition is that with *z >1* (as from EMAR) the decrease of diversity is much faster than the decrease of habitat, but with *z < 1* (as from MAR), there is a (desirable) slower dynamics of genetic loss. In the latter, despite habitats disappearing, reservoirs of mutations distributed across different locations enable conservation of certain variation. To study which one is more likely and to observe the stochastic nature of genetic diversity loss, we simulated in silico population extinctions of map cells from the Arabidopsis map (Fig. 1) and directly estimated from the genome matrix of remaining individuals the remaining genetic diversity. These simulations were implemented to capture different hypothesised patterns of extinction (see main text). All, however, agree with the more hopeful estimate of *z_MAR_* ≈ 0.3.

**Fig. S18.**
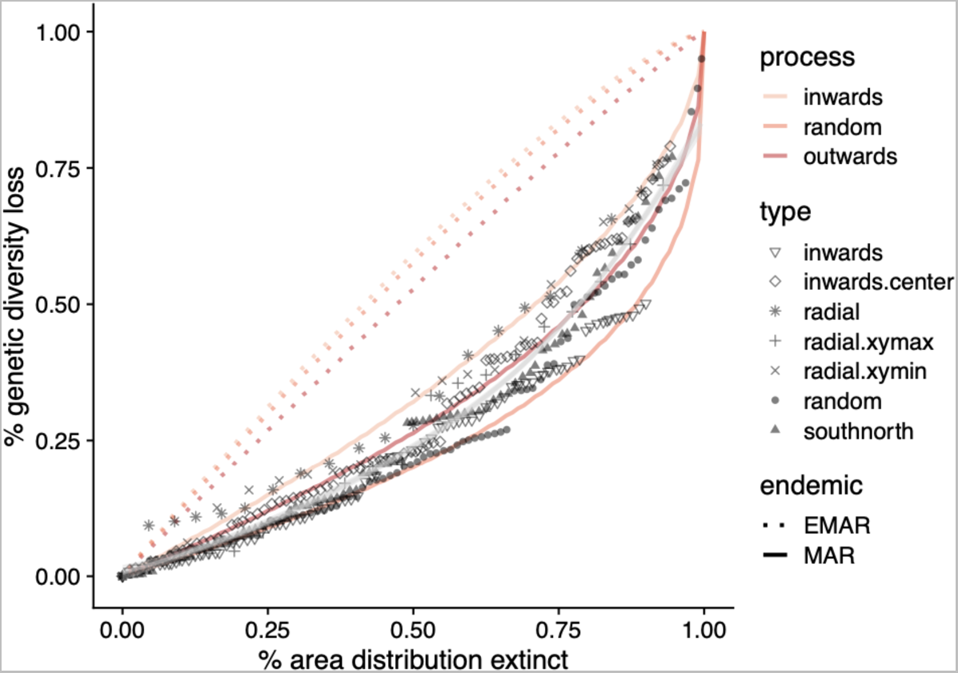
Loss of mutations with habitat loss in *A. thaliana*. Predictions based on MAR and EMAR functions and in silico extinction stochastic simulations in *A. thaliana*.

To study the fit of the genetic loss predictions based on MAR relationships and the results from computer simulations, we calculated a pseudo-*R^2^* based on the squared differences between the predicted line and the “observed” genetic loss as: 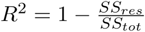. This results in a high fit *R^2^=*0.872 of the MAR, built from random samples of distribution areas, while the EMAR had a poor fit due to overestimation of genetic loss: *R^2^=*-0.710 (negative values indicate predictions are worse than the mean of the data).

##### III.5 Potential impacts of genetic loss in adaptability

Although likely imperfect, Genome-Wide Associations could help to understand the relevance of mutations in different frequency classes in model organisms such as *Arabidopsis thaliana*. Fig. S19 shows the site frequency spectrum and a metric of the “total accumulated effect in fitness” of the alleles in every bin. Effect sizes were retrieved from GWA on lifetime fitness of 515 ecotypes in outdoor experiments (*35*). The average effect size across 8 fitness GWA from 8 experimental combinations were used: high/low precipitation, high/low latitude of outdoor stations, and high/low plant density. This exercise showcases the phenomenon that low frequency variants often have strong effect sizes, which is expected under a stabilising selection quantitative model (*36*). Because low frequency alleles will be the first to be lost during a bottleneck (as would happen with the rapid extinction of populations of a species), we may expect to lose variants that are related to fitness and thus potentially lose diversity that could be advantageous in some environments. Alternatively, deleterious mutations are also expected to be at low frequency, in which case would also make them more easily lost.

**Fig. S19.**
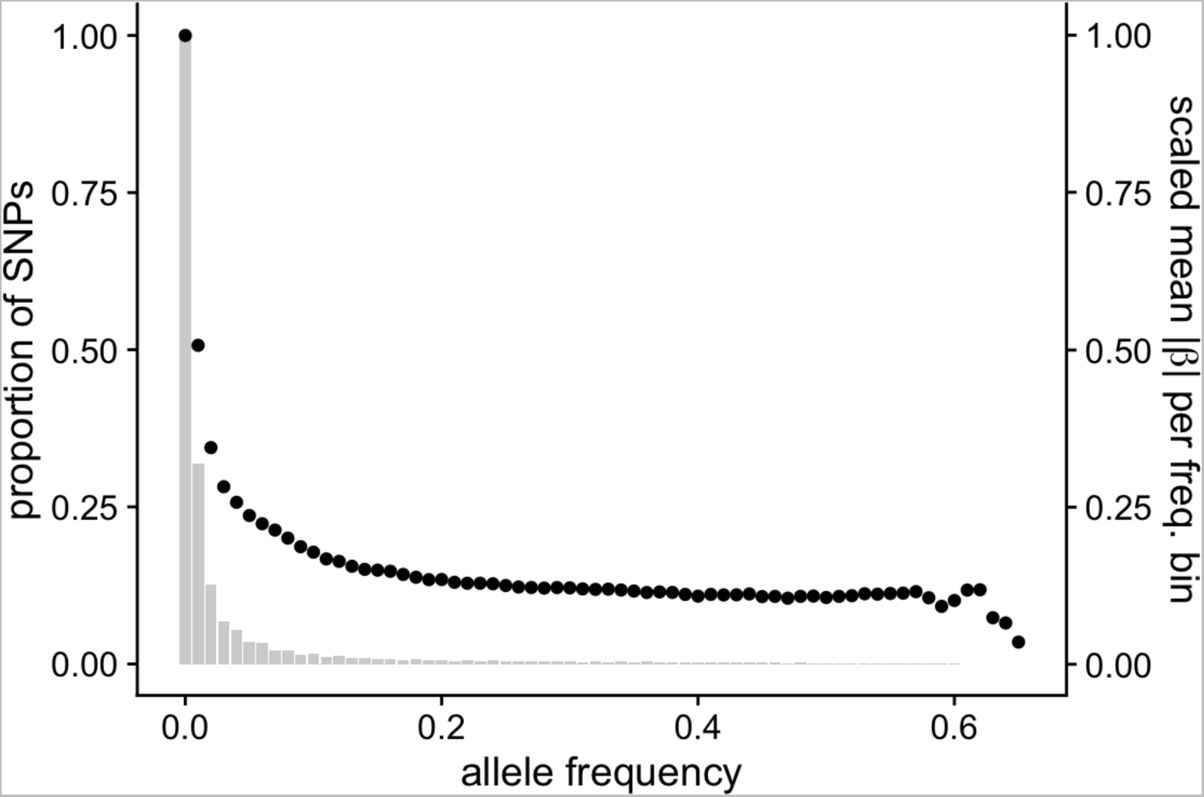
Bias of low frequency mutations and effect size for fitness traits in *A. thaliana*. Grey bars represent the site frequency spectrum (scaled for visualisation purposes). The black dots represent the mean absolute effects of alleles as estimated from GWAs with 515 accessions scored for fitness traits in 8 outdoor experiments.

To further build intuition on the progress of extinction in relation to loss of genetic diversity that is not neutral, we repeated warm edge extinction simulations with several subsets of alleles: randomly selected SNPs, SNPs that were associated positively in 2 environments (low precipitation Spain and high precipitation Germany) (labelled globally positive), and SNPs that were associated positively in one environment and negatively in the other (labelled antagonistic pleiotropic or putatively locally-adaptive). This (Fig. S20) supports our intuition that although putatively functional alleles (or alleles tightly linked to such functional ones) may have slower loss dynamics than neutral variants due to a high frequency and *z_MAR_*, certain population extinction patterns may actually lead to rapid loss of potentially-adaptive genetic diversity. The complexity of these patterns, together with the evolutionary feedback created by lowering genetic standing variation that affects fitness, make the inference of adaptive capacity loss even more difficult than just inferring the loss of genetic diversity itself.

**Fig. S20.**
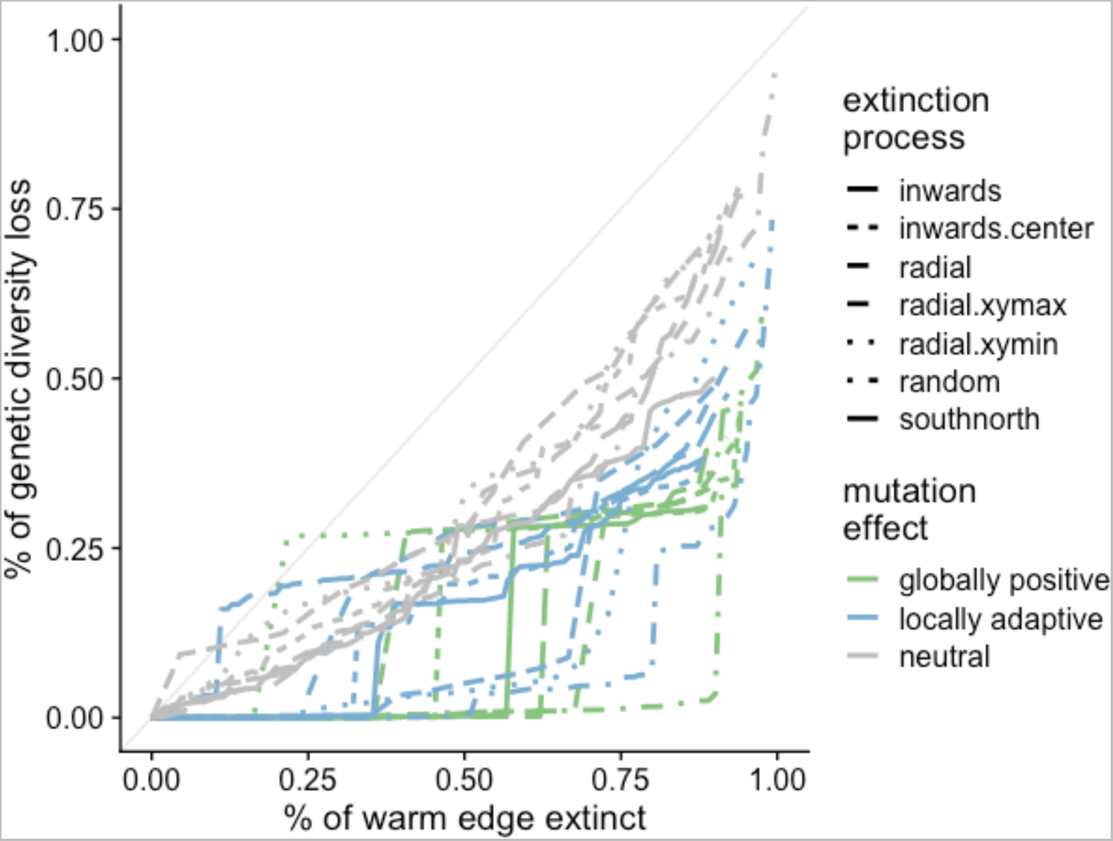
Simulations illustrating the potential loss of locally-adaptive mutations in *A. thaliana*. Simulations of extinction using multiple patterns of population losses with different subsets of alleles ascertained to show positive associations in fitness GWA in two outdoor experiments (green), positive associations in one environment (e.g. low precipitation) but negative in a second environment (e.g. high precipitation) or vice versa (green). These were compared to a random set (grey).

**Fig. S21.**
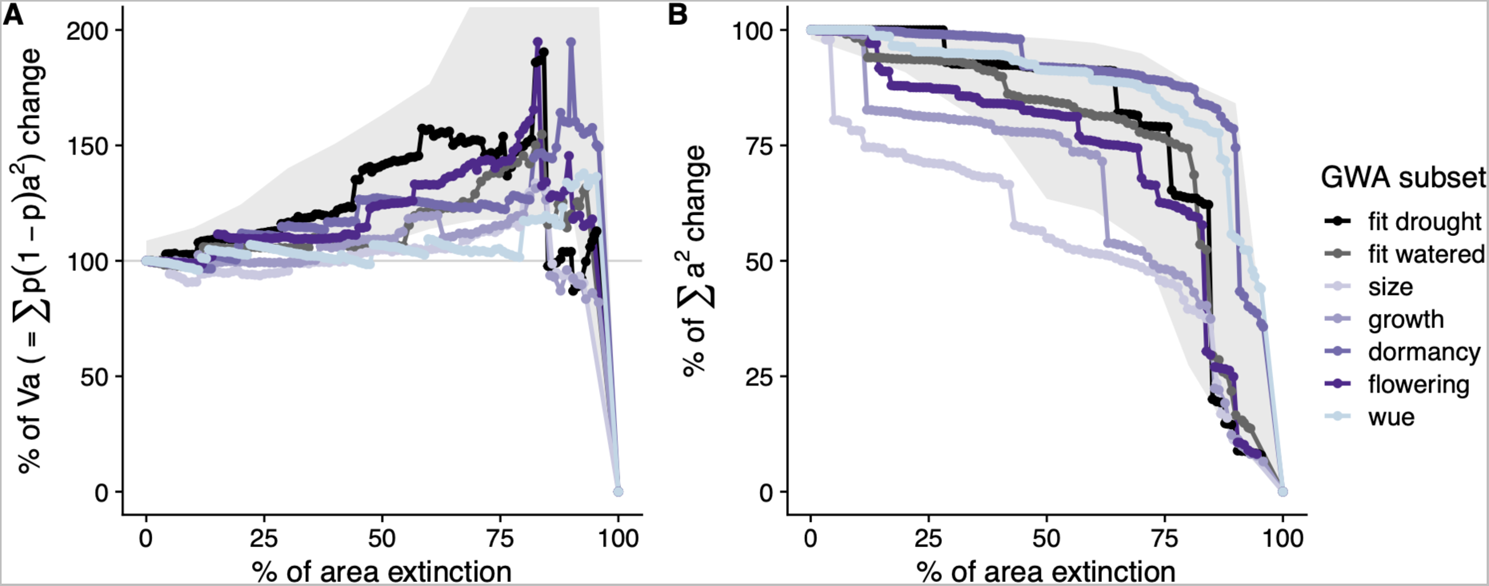
Extinction simulations showing proxies of adaptive capacity of *A. thaliana*. Using estimated allele effect sizes from 10,000 SNPs in the 1% P-value tails of several Genome-Wide Associations, we show (A) Percentage of change of Va as a proxy of adaptive potential and (B) raw square sum of allele effects to showcase the inflating effect of intermediate frequency alleles. Grey background shape indicates the minimum and maximum boundaries of trajectories created by replicated frequency-matched non-effect sets of SNPs (one per GWA). The trajectories of some effect alleles appear to show faster loss than the non-effect background trajectories.

##### III.6 Case study of a massive natural bottleneck

A recent colonisation of North America by *Arabidopsis thaliana* can help us understand the recovery of genetic variation. Whole-genome sequencing of 100 specimens of North American *A. thaliana* indicates that it migrated from its native range of Europe to North America in the 17th century, and began spreading across the continent from a genetically-homogeneous population (*37*). Despite ideal conditions to re-gain genetic diversity—a continental population expansion aided by human travel (*38, 39*)—only ~8,000 new mutations were detected through spontaneous accumulation, equivalent to only ~0.067% of the species-wide native genetic diversity. Because most of these mutations are at very low frequency, as expected during population expansion, the scaling of genetic diversity with area is approximately 1 (*z_MAR_* = 1.025 [CI95%: 0.878 - 1.173]).

#### IV. The mutations-area relationship in diverse species

Every dataset was retrieved online either from the published article in the form of VCF or fastq files, or provided by the study authors upon request. All datasets were first transformed into PLINK files using PLINK v1.9 (*40*). For computational efficiency, and since we showed random subsampling does not appear to affect calculations of *z_MAR_* (Section III.3), we conducted all analyses with up to 10,000 randomly selected SNPs for each species sampled genome-wide, or in the largest chromosome for those species with large genomes. We aim to use mostly unfiltered SNP datasets to avoid ascertainment biased toward intermediate frequency SNPs, and therefore we did not apply a MAF filter for any analyses. By default, PLINK transforms SNP matrices into biallelic (if multiallelic, it takes the two most common alleles). Although the preservation of structural genetic variation may also be relevant and may have important consequences in adaptation (*41*), we do not expect dramatic differences in their scaling relationship compared to biallelic SNPs, as their SFS are relatively similar (Structural variants may show a skew to lower frequency, resulting in steeper *z_MAR_*. By excluding those, our analyses may be conservative). In order to properly characterise the geographic distribution of a mutation using all available geo-tagged individuals, we filtered for genotyping rate (plink --geno), and the final value is reported per dataset.

Details for dataset processing or homogenization are described below.

- The 1001 Arabidopsis Genomes Consortium (*29*) generated a WGS Illumina sequencing dataset of *Arabidopsis thaliana* comprising 1,135 individuals and 11,769,920 SNPs. The VCF with the data is available at: https://1001genomes.org. The raw sequencing data is available at https://www.ncbi.nlm.nih.gov/bioproject/PRJNA273563. These included recently colonised regions such as North America or Japan. Analyses of *z_MAR_* were calculated only for the native range, which comprises most of the species diversity (>99%) and 1001 individuals. For computational efficiency, we conducted analyses using randomly sampled SNPs from chromosome 1, as we did not observe any difference when sampling from other chromosomes. A number of MAR approaches were tested in this species (section III). For homogeneity, the final reported estimate (Table 1) was conducted following the same procedures as other species with a random sample of 10,000 SNPs.
- Lucek & Willi (*42*) recently published a dataset of WGS Illumina sequencing 108 *Arabidopsis lyrata* individuals from North America, which the authors directly shared as a VCF. The raw data is available at https://www.ncbi.nlm.nih.gov/bioproject/?term=PRJEB30473. We retrieved the latitude/longitude data from the supplemental material. We applied a genotyping rate filter ending with a dataset of 0.955431 genotyping rate. 10,000 SNPs were subsetted at random from the genome-wide data.
- Kreiner et al. (*43*) WGS Illumina sequenced 165 individuals of *Amaranthus tuberculatus*. The raw data is available in the link https://www.ebi.ac.uk/ena/browser/view/PRJEB31711. The authors provided a VCF. Overall, 155 individuals contained latitude and longitude information and were kept for the analyses. The genotyping rate was 0.98162 and we subsetted randomly 10,000 SNPs.
- Supple et al. (*44*) generated a dataset of *Eucalyptus melliodora* of 275 individuals from 36 broadly distributed populations. The dataset was produced by Illumina sequence Genotyping-by-Sequencing (GBS) libraries digested with ApeKI as in Elshire et al. (2011). The raw data is available at https://www.ncbi.nlm.nih.gov/bioproject/PRJNA413429/. The authors provided the dataset in PLINK format. Genotyping rate was 0.769807 but we did not apply a further filter to avoid reducing the total number of variants. We conducted analyses with all 9378 SNPs. The genotyping rate in this dataset is likely not problematic as the total number of GPS locations is 36, with multiple individuals sampled closely. This sampling scheme probably allows to characterise an allele’s distribution correctly despite the lower genotyping rate.
- Vallejo-Marin et al. (*45*) generated a GBS dataset of 521 Mimulus plants, with 286 samples being *Mimulus guttatus* from its native distribution. Libraries for Genotyping-By-Sequencing were prepared with PstI enzyme as described in Twyford & Friedman (2015) and sequenced using Illumina. The VCF of this dataset is available at http://hdl.handle.net/11667/168 and was also directly shared by the authors. After applying a filtering for missingness, we ended up with a genotyping rate of 0.904192 and 1,498 SNPs, which were used for the analyses.
- Lovell & MacQueen (*46*) generated a WGS Illumina sequencing dataset of Switchgrass, *Panicum virgatum*, of a collection of 732 individuals and 33,905,044 variants. The raw data is available at: https://www.ncbi.nlm.nih.gov/bioproject/PRJNA622568. The authors provided a VCF file and latitude/longitude tables. 576 individuals were from natural collections. The dataset contains also other collections such as cultivars, which were not used to build the MAR. The genotyping rate was 0.976393 and analyses were conducted with 10,000 SNPs drawn from the largest chromosome.
- MacLachlan et al. (*47*) generated a SNP chip dataset of *Pinus contorta* comprising 929 trees with latitude and longitude information and 32,449 SNPs. Genotyping was conducted with the AdapTree lodgepole pine Affymetrix Axiom 50,298 SNP array and data was provided in the supplemental material of the paper along with custom scripts to parse the data. The database is available at https://datadryad.org/stash/dataset/doi:10.5061/dryad.ncjsxkstp. The genome matrix was transformed into PLINK. The genotyping rate was 0.959146, and analyses were conducted with 10,000 randomly drawn SNPs. The fact that this dataset was created with ascertained SNPs likely generates a frequency bias. In Fig. S22, one can see that this may be a problem to calculate *z_MAR_*, as the mutations~area graph appears nonlinear and rapidly saturates. This confirms the expectation that SNPs are ascertained to be common, as they are discovered immediately with very few samples.
- Tuskan et al. (*48*) WGS Illumina sequenced 882 *Populus trichocarpa* trees. The dataset includes 28,342,826 SNPs. The data is available under this DOI https://doi.ccs.ornl.gov/ui/doi/55 which redirects to a globus data sharing platform. The authors provided the dataset as a VCF along with latitude/longitude coordinates. This dataset was downsampled to the first chromosome. The genotyping rate was 0.921191, and 10,000 SNPs were randomly sampled for analyses.
- The Anopheles gambiae 1000 Genomes Consortium (*49*) (Phase 2) produced Whole-Genome Illumina sequencing data for 1142 wild-caught mosquitoes of *Anopheles gambiae*. All raw and processed data are available through https://www.malariagen.net/data. We downloaded a VCF and latitude/longitude coordinate files. The VCF was filtered for genotyping rate ending up at a 0.998895 rate. For efficiency, 10,000 randomly-selected SNPs from the VCF of the largest chromosome 2L were used for analyses downstream.
- Fuller et al. (*50*) WGS Illumina sequenced 253 coral individuals of *Acropora millepora* in 12 reefs. The dataset was downloaded as fastq files from the published online material from https://www.ncbi.nlm.nih.gov/bioproject/?term=PRJNA593014, and SNPs were called as described in the supplemental material ending with 17,931,448, which were filtered to achieve a genotyping rate of 0.935709 for a total of 2,512 SNPs, which were used in the analyses.
- Ruegg et al. (*51*) generated a dataset of 219 birds *Empidonax traillii*, for which 199 could be matched with geographic coordinates. SNPs were ascertained from several publications using RAD seq and Fluidigm 96.96 IFC described and available in their repository https://github.com/eriqande/ruegg-et-al-wifl-genoscape. A total of 349,014 SNPs were parsed using their custom scripts and we transformed them into PLINK files. A genotyping rate filter was applied ending with a 0.96061 rate and 195,700 SNPs. 10,000 SNPs were selected at random for downstream analyses. Similarly, as with the *Pinus contorta*, the incorporation of some ascertained SNPs in the dataset based on Fluidigm technology could lead to quick saturation of the MAR curve (Fig. S22).
- Bay et al. (*52*) generated a dataset of 199 *Setophaga petechia* birds using a Restriction site–associated DNA sequencing (RAD-Seq). The raw data is available at https://www.ncbi.nlm.nih.gov/bioproject/421926. The authors shared a VCF file, with a genotyping rate of 0.962419 and a total of 104,711 SNPs. 10,000 SNPs were selected at random for downstream analyses.
- Kingsley et al. (*53*) produced a dataset of 80 *Peromyscus maniculatus* deermice, for which 78 could be matched with geographic locations. The SNP dataset was produced using MY-select capture followed by Illumina sequencing. The VCF and PLINK files are available via Figshare at https://doi.org/10.6084/m9.figshare.1541235. The dataset included a total of 14,076 variants which were filtered to achieve a genotyping rate of 0.940411 for 2,946 SNPs, which were used in subsequent analyses.
- We identified two published datasets for wolves. Smeds et al. (*54*) produced a WGS Illumina sequencing dataset and combined it with pre-existing datasets for a total of 349 local dog breeds and wolves, of which 230 were *Canis lupus* from natural populations. However, these samples did not have GPS locations assigned. The second dataset we identified was from Schweizer et al. (*55*), which contained 107 geo-tagged grey wolves from North America using a capture and resequencing approach for 1040 genes. The raw data is available at https://trace.ncbi.nlm.nih.gov/Traces/sra/?study=SRP065570, and meta-data along with a VCF area available at https://doi.org/10.1111/mec.13467. This data contained 13,092 SNPs at 0.993061 calling rate, and a better geographic resolution. We report data for the second dataset.
- The 1000 Genome Consortium (*56*) created WGS Illumina sequencing for over 2,504 humans and 24 unique geographic locations. We downloaded chromosome 1 from http://ftp.1000genomes.ebi.ac.uk/vol1/ftp/datacollections/ <u>1000G2504highcoverage/working/20190425NYGCGATK/</u> and gathered the population locations from https://www.internationalgenome.org/data-portal/population. To conduct analyses, we subsampled 10,000 SNPs at genotping rate 0.991069.
- Palacio-Mejia (*57*) used WGS for 591 *Panicum hallii* individuals to sequence at low coverage. The raw data is available at https://www.ncbi.nlm.nih.gov/bioproject/PRJNA390994. The authors shared an unfiltered VCF of 45,589 SNPs. Because of the low-coverage, stringent filters of calling rates as used for other species would lead to removing all SNPs, and we settled on a genotyping rate of 0.825824 for 242 variants, all of which were used for downstream analyses.
- Royer et al. (*58*) produced a SNP dataset using RAD-Seq based Genotyping-By-Sequencing of 290 *Yucca brevifolia* (Joshua Tree) individuals. A total of 10,695 SNPs with a genotyping rate of 0.897501 wre used for the analyses. The data was available at Dryad https://datadryad.org/stash/dataset/doi%253A10.5061%252Fdryad.7pj4t.
- Kapun et al. (*59*) produced a WGS dataset of pooled *Drosophila melanogaster*, sequencing ~80 pooled individuals from each of 271 populations as part of the European “Drosophila Evolution over Space and Time” (DEST) project. A total of 5,019 shared SNPs with a genotyping rate of 0.937697 were used for analyses. The dataset, both raw and processed, is available through https://dest.bio.
- Di Santo et al. (*60*) studied the highly-threatened species *Pinus torreyana*. They used Genotyping-by-Sequencing of 242 individuals of the last remaining populations. The dataset is not yet available through NCBI but the authors kindly shared a VCF directly with us. From a total set of 166,564 SNPs with a genotyping rate of 0.964632, 10,000 were randomly selected for our analyses.
- von Seth et al. (*61*) studied the highly-threatened species *Dicerorhinus sumatrensis*. They used Illumina WGS of 16 individuals of the last remaining populations. The raw data is available at https://www.ebi.ac.uk/ena/browser/view/PRJEB35511. The authors shared a VCF. In total, this comprises a set of 8,870,513 SNPs, with a genotyping rate of 0.854862, which we did not further filter due to the small number of individuals. For computational efficiency we selected 10,000 SNPs from the largest chromosome.

Information and results per species are gathered in Table 1 and its extended version, Table S10, and the average *z_MAR_* across species are provided in Table S11.

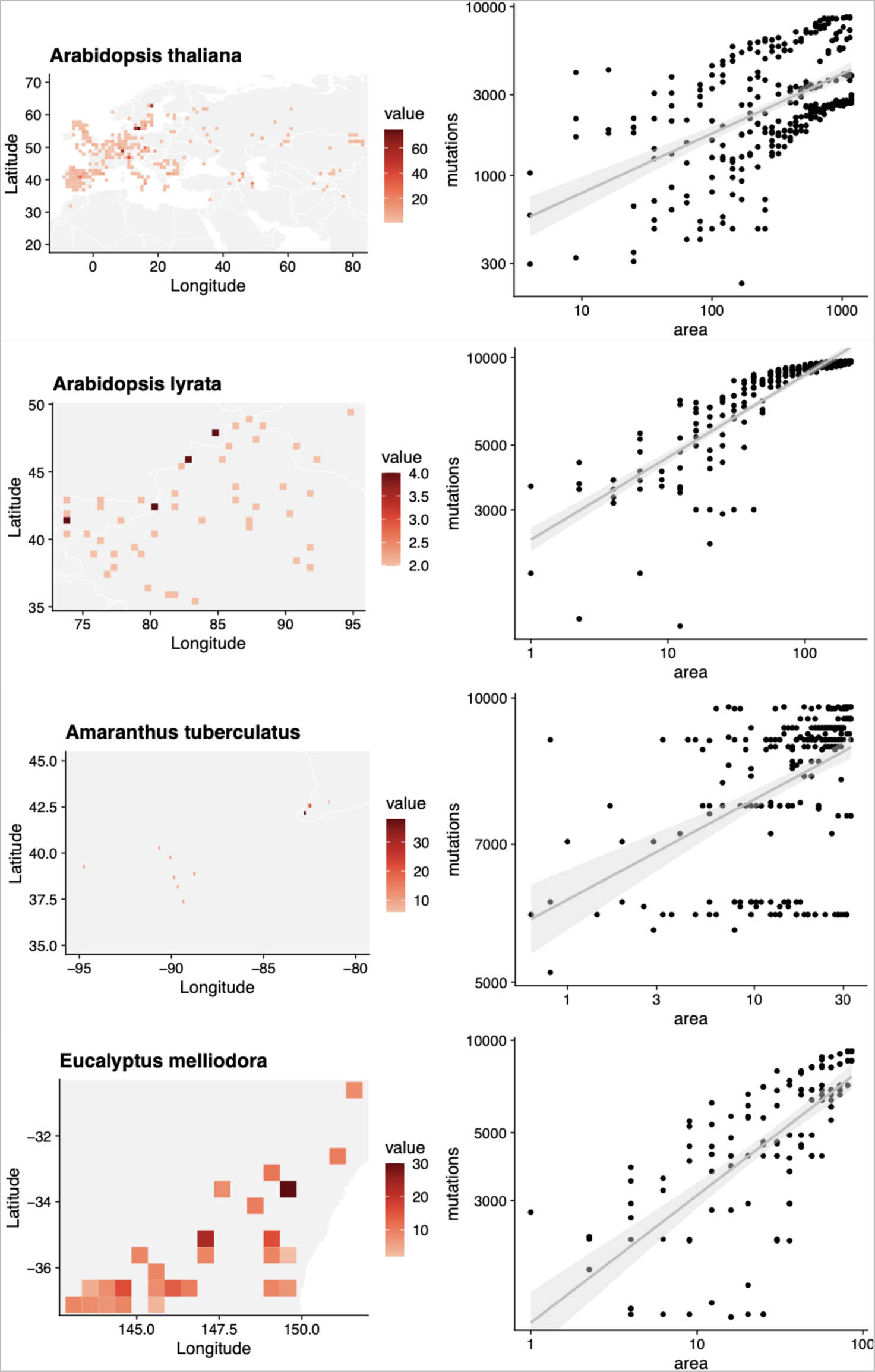

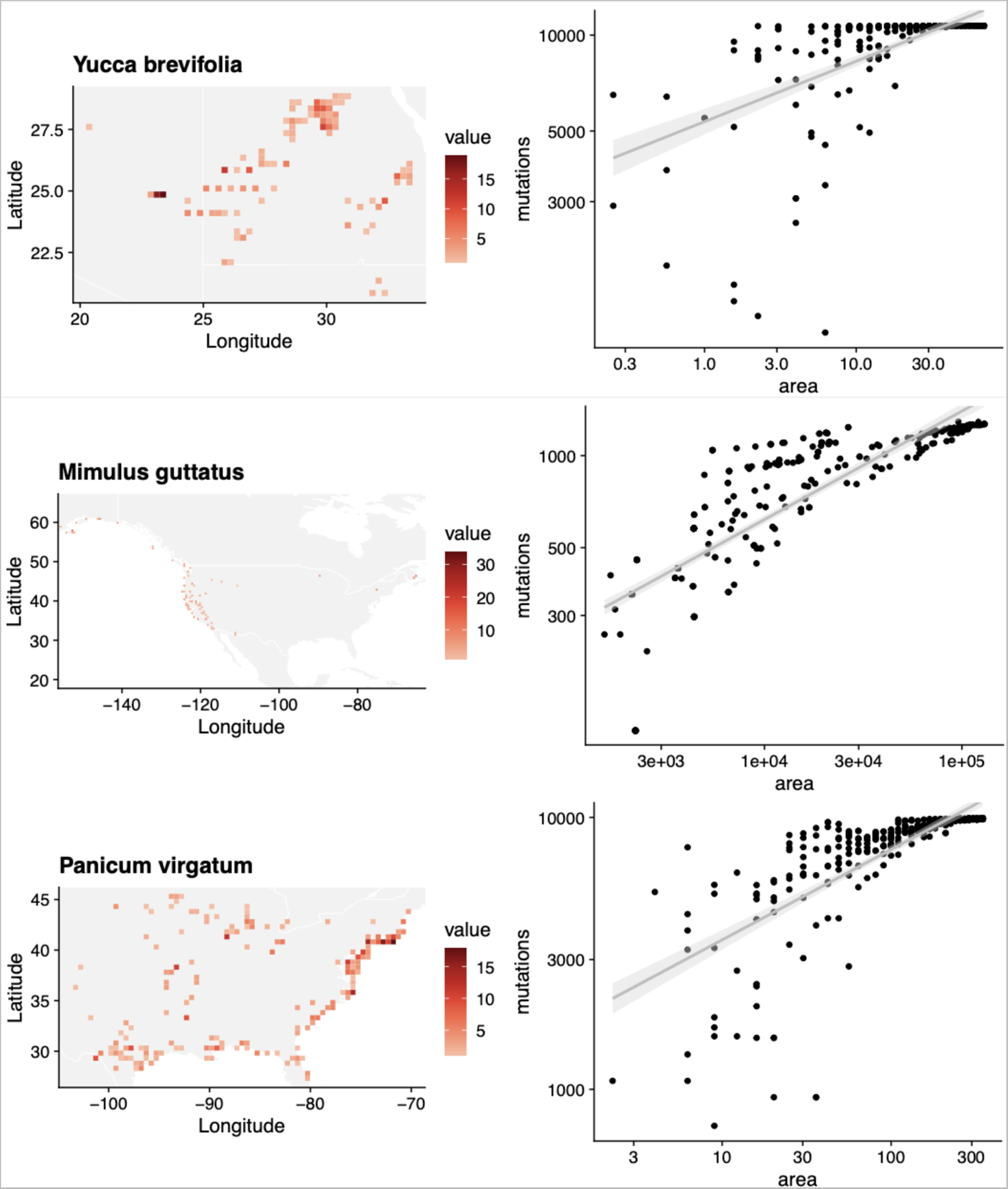

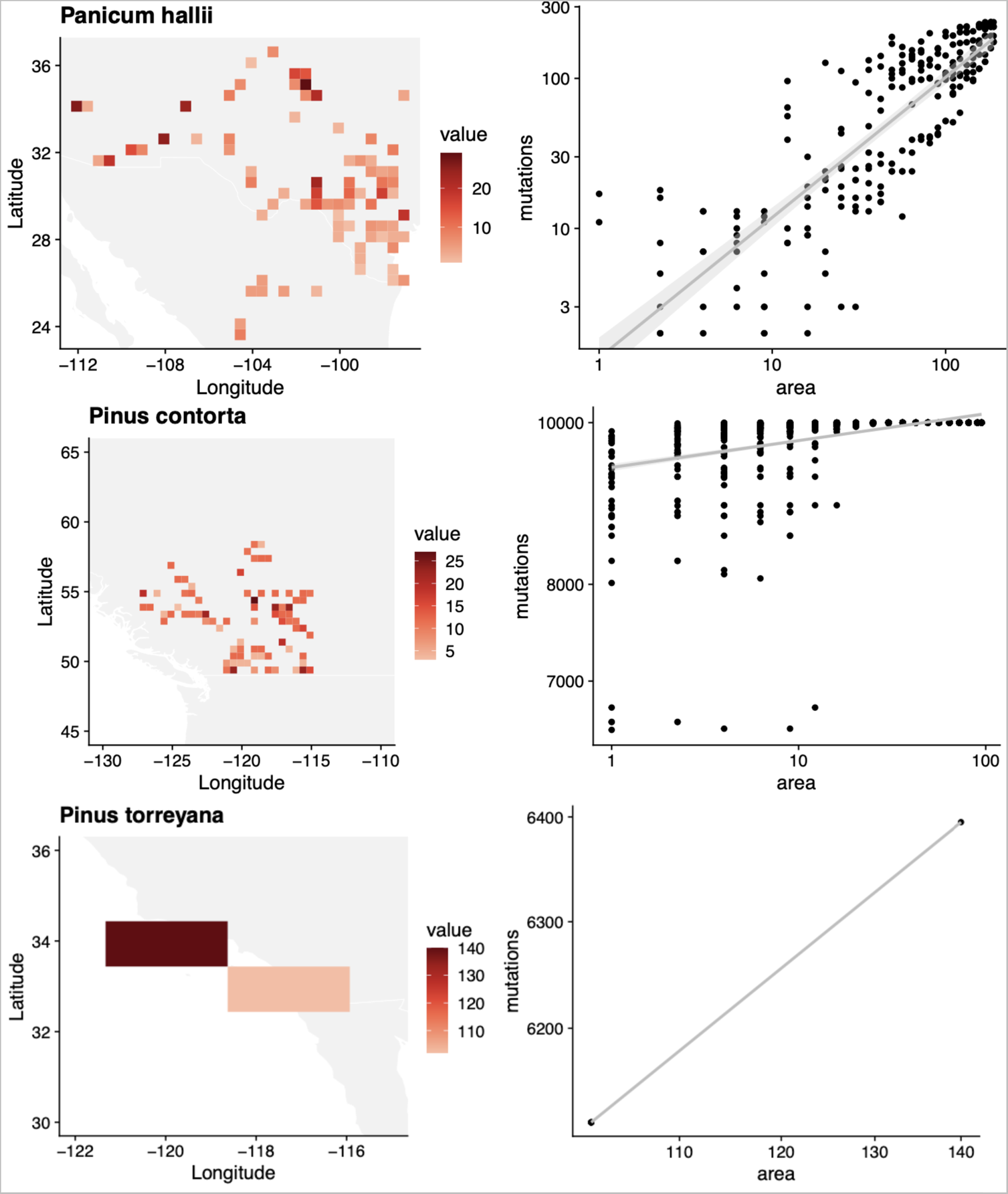

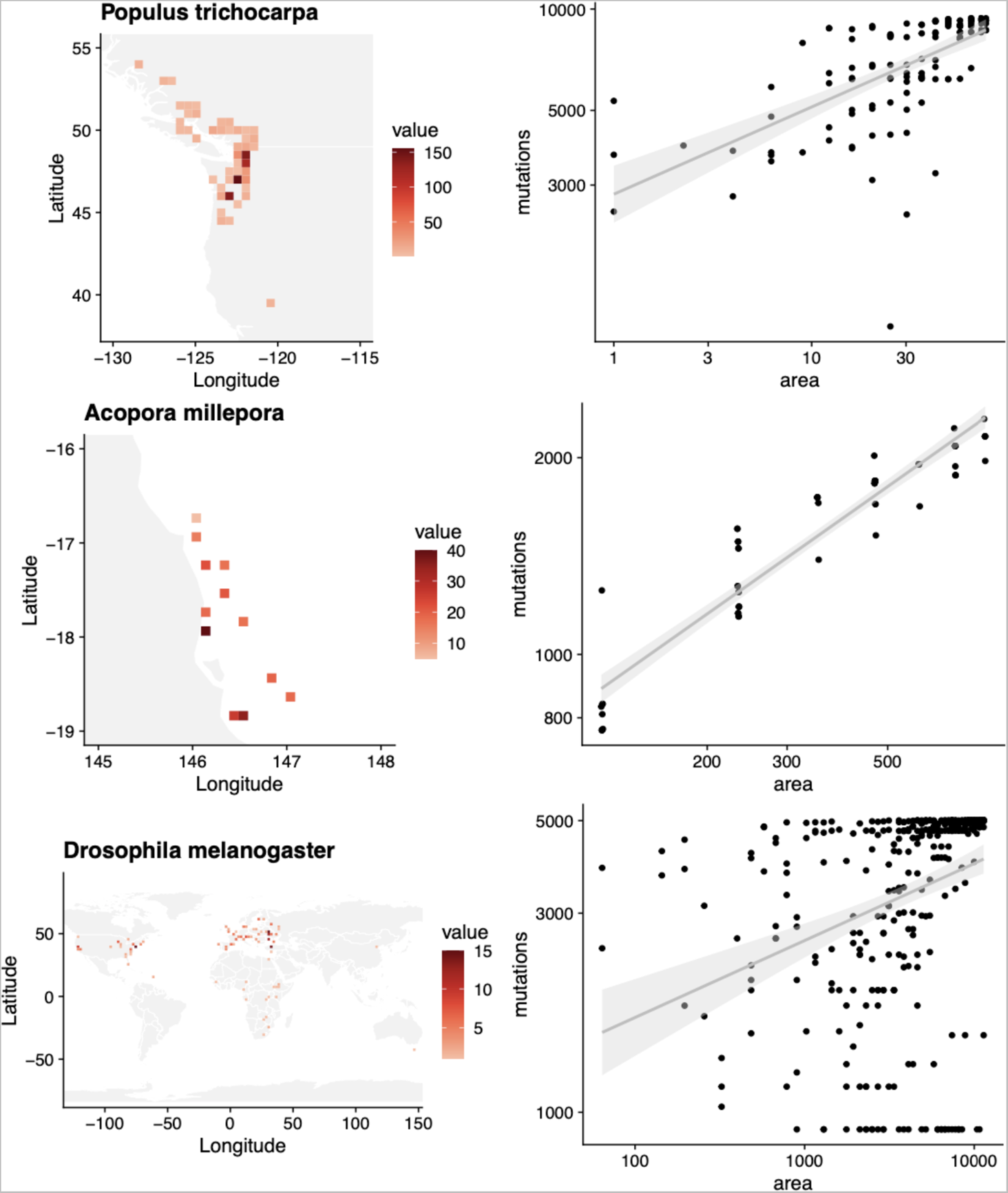

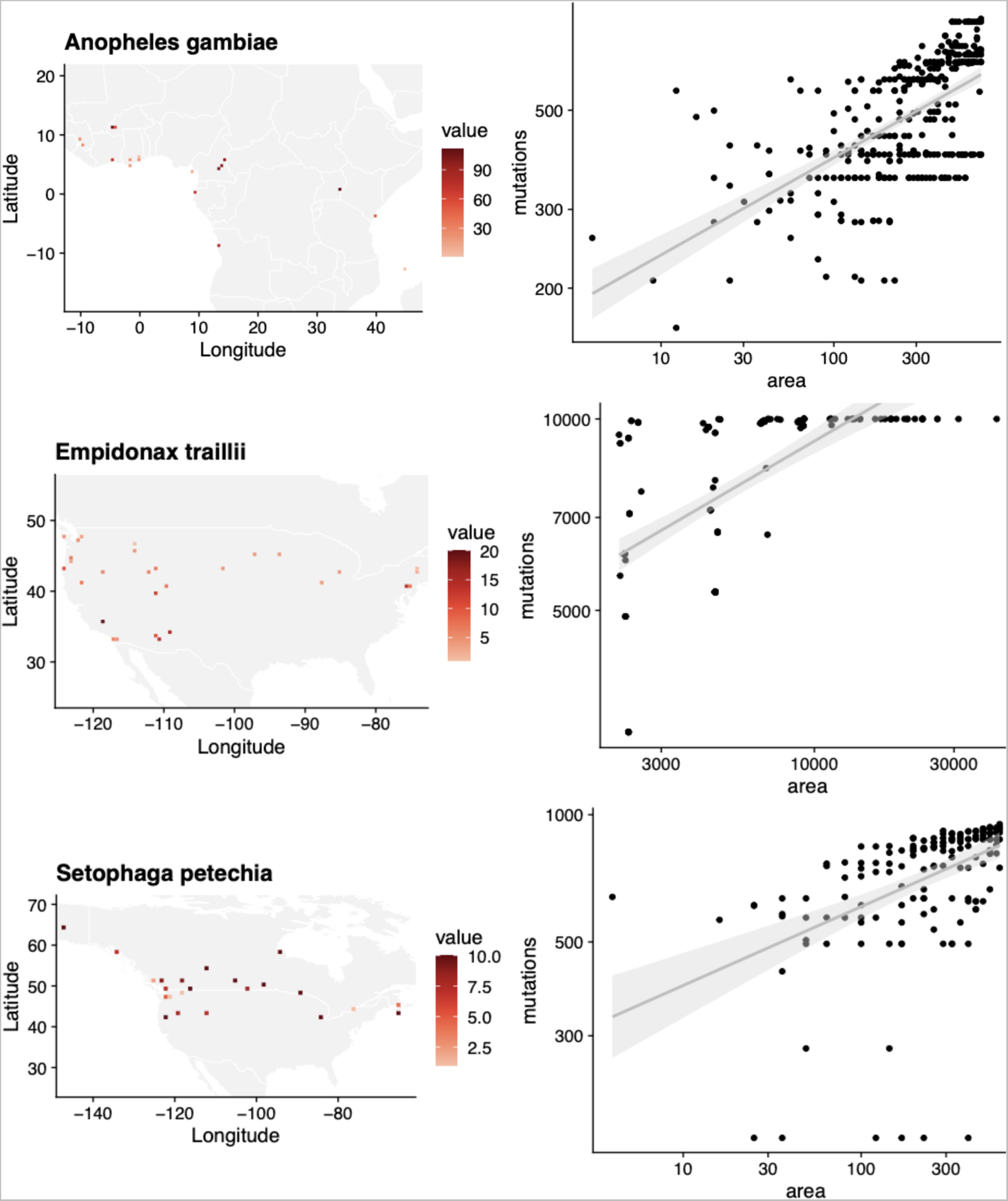

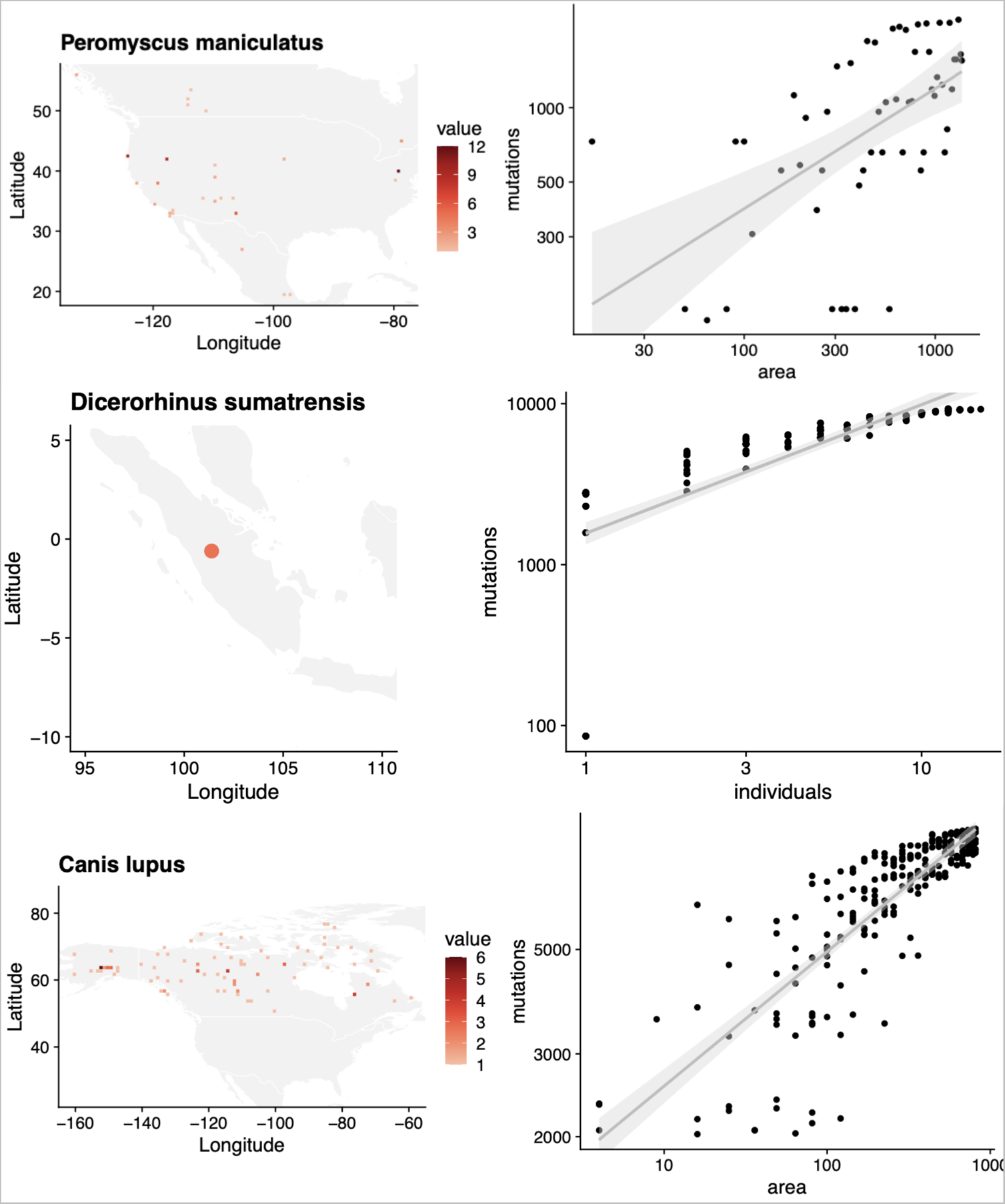

**Fig. S22.**
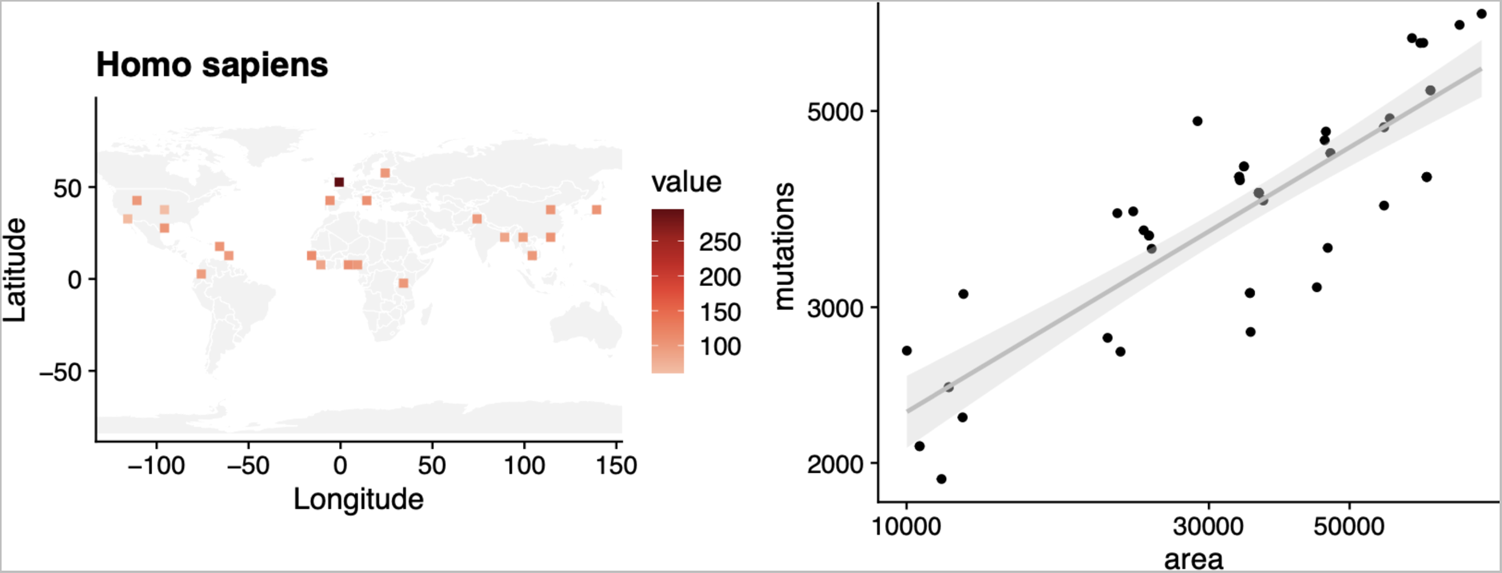
MAR summaries across species. For each species we plot (left) the map of sample density in space and (right) the mutations-area relationship. (The locations of 16 Dicerorhinus sumatrensis are unknown so only Sumatra is shown. Pinus torreyana was only found in two extant populations.)

**Table S10.**
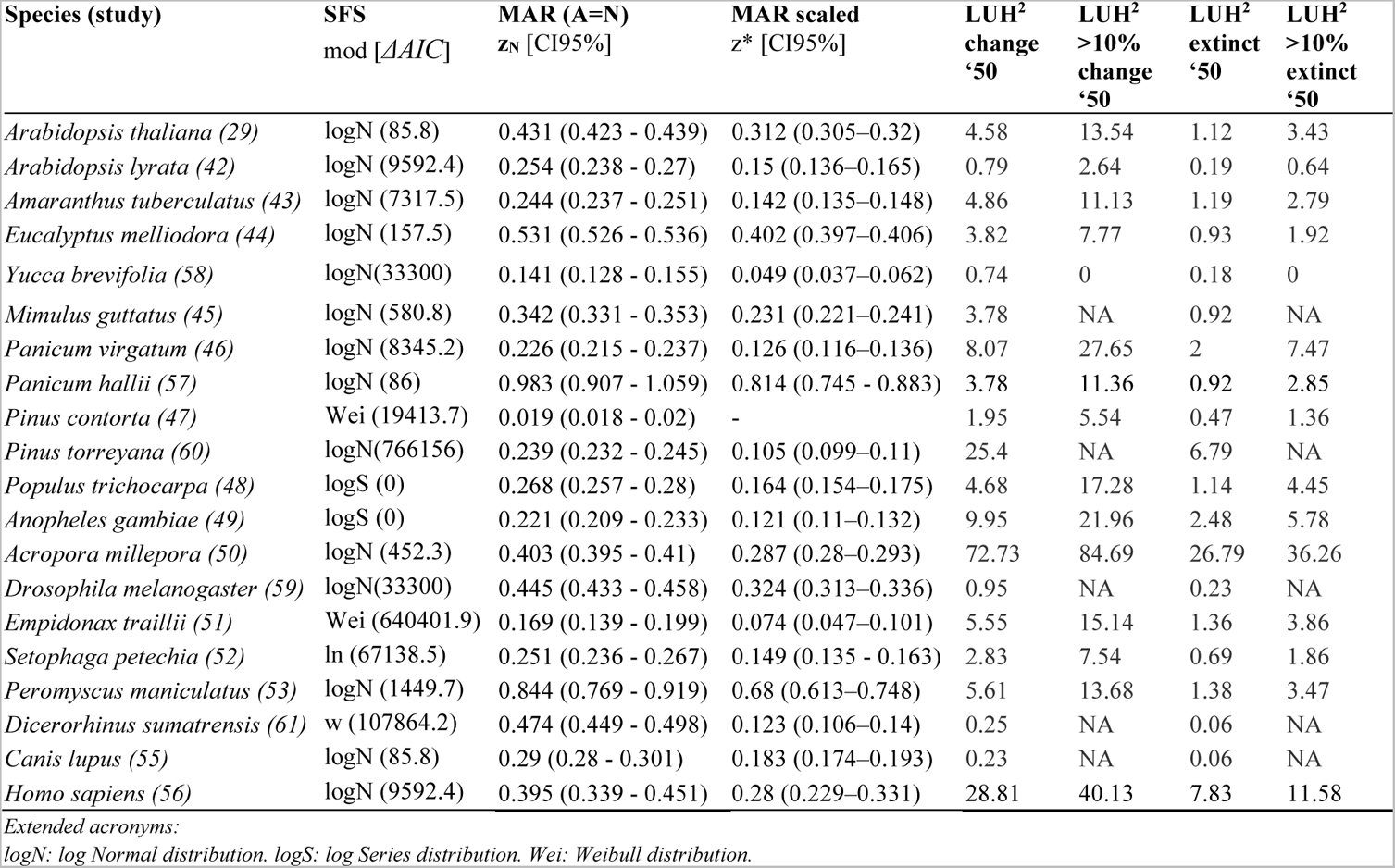
The mutations-area relationship across species. Extended Table 1. The Mutations-Area Relationship (MAR) fitted with Area = Individuals and the scaled version. In the main text areas to protect 90% of genetic diversity per species are provided given the scaled z*. Here, we also provide the average estimated area based on % of grid cells per species to be transformed from 2015 to 2050 using the LUH^2^ dataset, the area where at least 10% of grid cells will be transformed, and the genetic loss corresponding to those area transformations (see section V.2).

| Species (study) | SFS<br>mod [AIC] | MAR (A=N)<br>$z_N$ [CI95%] | MAR scaled<br>$z^*$ [CI95%] | LUH<sup>2</sup><br>change<br>'50 | LUH<sup>2</sup><br>>10%<br>change<br>'50 | LUH<sup>2</sup><br>extinct<br>'50 | LUH<sup>2</sup><br>>10%<br>extinct<br>'50 |
| --- | --- | --- | --- | --- | --- | --- | --- |
| *Arabidopsis thaliana* (29) | logN (85.8) | 0.431 (0.423 - 0.439) | 0.312 (0.305–0.32) | 4.58 | 13.54 | 1.12 | 3.43 |
| *Arabidopsis lyrata* (42) | logN (9592.4) | 0.254 (0.238 - 0.27) | 0.15 (0.136–0.165) | 0.79 | 2.64 | 0.19 | 0.64 |
| *Amaranthus tuberculatus* (43) | logN (7317.5) | 0.244 (0.237 - 0.251) | 0.142 (0.135–0.148) | 4.86 | 11.13 | 1.19 | 2.79 |
| *Eucalyptus melliodora* (44) | logN (157.5) | 0.531 (0.526 - 0.536) | 0.402 (0.397–0.406) | 3.82 | 7.77 | 0.93 | 1.92 |
| *Yucca brevifolia* (58) | logN(33300) | 0.141 (0.128 - 0.155) | 0.049 (0.037–0.062) | 0.74 | 0 | 0.18 | 0 |
| *Mimulus guttatus* (45) | logN (580.8) | 0.342 (0.331 - 0.353) | 0.231 (0.221–0.241) | 3.78 | NA | 0.92 | NA |
| *Panicum virgatum* (46) | logN (8345.2) | 0.226 (0.215 - 0.237) | 0.126 (0.116–0.136) | 8.07 | 27.65 | 2 | 7.47 |
| *Panicum hallii* (57) | logN (86) | 0.983 (0.907 - 1.059) | 0.814 (0.745 - 0.883) | 3.78 | 11.36 | 0.92 | 2.85 |
| *Pinus contorta* (47) | Wei (19413.7) | 0.019 (0.018 - 0.02) | - | 1.95 | 5.54 | 0.47 | 1.36 |
| *Pinus torreyana* (60) | logN(766156) | 0.239 (0.232 - 0.245) | 0.105 (0.099–0.11) | 25.4 | NA | 6.79 | NA |
| *Populus trichocarpa* (48) | logS (0) | 0.268 (0.257 - 0.28) | 0.164 (0.154–0.175) | 4.68 | 17.28 | 1.14 | 4.45 |
| *Anopheles gambiae* (49) | logS (0) | 0.221 (0.209 - 0.233) | 0.121 (0.11–0.132) | 9.95 | 21.96 | 2.48 | 5.78 |
| *Acropora millepora* (50) | logN (452.3) | 0.403 (0.395 - 0.41) | 0.287 (0.28–0.293) | 72.73 | 84.69 | 26.79 | 36.26 |
| *Drosophila melanogaster* (59) | logN(33300) | 0.445 (0.433 - 0.458) | 0.324 (0.313–0.336) | 0.95 | NA | 0.23 | NA |
| *Empidonax traillii* (51) | Wei (640401.9) | 0.169 (0.139 - 0.199) | 0.074 (0.047–0.101) | 5.55 | 15.14 | 1.36 | 3.86 |
| *Setophaga petechia* (52) | ln (67138.5) | 0.251 (0.236 - 0.267) | 0.149 (0.135 - 0.163) | 2.83 | 7.54 | 0.69 | 1.86 |
| *Peromyscus maniculatus* (53) | logN (1449.7) | 0.844 (0.769 - 0.919) | 0.68 (0.613–0.748) | 5.61 | 13.68 | 1.38 | 3.47 |
| *Dicerorhinus sumatrensis* (61) | w (107864.2) | 0.474 (0.449 - 0.498) | 0.123 (0.106–0.14) | 0.25 | NA | 0.06 | NA |
| *Canis lupus* (55) | logN (85.8) | 0.29 (0.28 - 0.301) | 0.183 (0.174–0.193) | 0.23 | NA | 0.06 | NA |
| *Homo sapiens* (56) | logN (9592.4) | 0.395 (0.339 - 0.451) | 0.28 (0.229–0.331) | 28.81 | 40.13 | 7.83 | 11.58 |

Extended acronyms:
logN: log Normal distribution. logS: log Series distribution. Wei: Weibull distribution.

##### IV.1 Exclusion of species from global averages

To avoid contaminating across-species averages of *z_MAR_* with estimates of species whose data we do not fully trust, we conducted global averages excluding species for which we are not confident *z_MAR_* reflects the correct species diversity-area relationships.

*Pinus contorta* showed a lower *z_MAR_* than what is expected in a theoretical baseline from individual sampling (section II). This is most likely due to this being the only species for which SNPs were previously ascertained to be intermediate frequency (i.e. the genome technology was a SNP chip). This alters SFS, so we are not confident the *z_MAR_* is the true parameter of the species.

*Yucca brevifolia* was a dense sampling of several local populations within a constrained area that is a hybrid zone. Since this species was not sampled range-wide we do not feel confident to include it in downstream analyses. The species also has a lower *z* than expected (Fig. S5)

*Pinus torreyana* only has two wild populations left, and therefore the MAR is based on two area sizes (Fig. S22). Because this is such a threatened species with already most of its range loss, we do not have confidence in the *z* parameter.

*Dicerorhinus sumatrensis* has only ~30 estimated adult individuals in the wild. Again we do not have confidence in the z parameter in such extinction-edge cases.

*Homo sapiens*. We exclude our own species.

**Table S11.** Mean z_MAR_ and other summary statistics across species. We selected those species that did not show artefacts in Fig. S22 or whose z_MAR_ overlapped with 0 to calculate a species-wide mean. See section IV.1.

| | $z_{MAR}$ | $z_{MAR} (A=N)$ | $z^*_{MAR} \text{ scaled}$ |
| --- | --- | --- | --- |
| mean | 0.31 | 0.39 | 0.27 |
| mean se | 0.038 | 0.053 | 0.048 |
| median | 0.25 | 0.29 | 0.18 |
| IQR | 0.15 | 0.19 | 0.17 |

Although we could not see any obvious patterns relating *z_MAR_* with certain groups of species (Table 1), we wondered whether any life history trait of the species analysed could explain the variation we observed (see Table S12 of traits). An ANOVA did not show any significant relationship. Because we know theoretically this parameter must be related to the degree of dispersal ability of genotypes of a species relative to the whole species geographic range, we expect traits involved in determining these to be good predictors. Future work will be necessary to validate this, as the sample size (n=19) may not permit enough power to detect these expected patterns.

**Table S12.** Traits, life history, and other characteristics of the analyzed species.

| Species | RedList | Known Decline | Kingdom | Reproduction | Pollination | Mobility | AreaRange |
| --- | --- | --- | --- | --- | --- | --- | --- |
| *Arabidopsis thaliana* | NO | NO | Plantae | Selfing | Selfing | Sessile | 27337467.4 |
| *Arabidopsis lyrata* | NO | NO | Plantae | Outcrossing | Vector | Sessile | 2791301.4 |
| *Amaranthus tuberculatus* | LC | NO | Plantae | Outcrossing | Vector | Sessile | 804124.8 |
| *Eucalyptus melliodora* | VU | NO | Plantae | Outcrossing | Wind | Sessile | 948699.3 |
| *Yucca brevifolia* | LC | YES | Plantae | Outcrossing | Vector | Sessile | 1213454.4 |
| *Mimulus guttatus* | LC | NO | Plantae | Outcrossing | Vector | Sessile | 25138310.6 |
| *Panicum virgatum* | LC | NO | Plantae | Outcrossing | Wind | Sessile | 6291400.2 |
| *Panicum hallii* | NO | NO | Plantae | Outcrossing | Wind | Sessile | 2188807.4 |
| *Pinus contorta* | LC | NO | Plantae | Outcrossing | Wind | Sessile | 886182.2 |
| *Pinus torreyana* | CR | YES | Plantae | Outcrossing | Wind | Sessile | 30781.95 |
| *Populus trichocarpa* | LC | NO | Plantae | Outcrossing | Wind | Sessile | 1119664.1 |
| *Drosophila melanogaster* | NO | NO | Animalia | Outcrossing | Activemating | Fly | 115208408 |
| *Anopheles gambiae* | NO | NO | Animalia | Outcrossing | Activemating | Fly | 19959809.9 |
| *Acropora millepora* | NT | YES | Animalia | Outcrossing | Activemating | Fly | 26725.9 |
| *Empidonax traillii* | LC | YES | Animalia | Outcrossing | Activemating | Fly | 7027395.2 |
| *Setophaga petechia* | LC | NO | Animalia | Outcrossing | Activemating | Fly | 15172431.15 |
| *Peromyscus maniculatus* | LC | NO | Animalia | Outcrossing | Activemating | Mobile | 22609152.6 |
| *Dicerorhinus sumatrensis* | CR | YES | Animalia | Outcrossing | Activemating | Mobile | 3335605.58 |
| *Canis lupus* | LC | NO | Animalia | Outcrossing | Activemating | Mobile | 19102403.5 |
| *Homo sapiens* | NA | NA | NA | NA | NA | NA | 80763121.8 |

**Table S13.** Association of traits, life history, and other characteristics with z_MAR_. Acronyms: NO=not assessed but likely non-threatened, LC=low concern, VU=vulnerable, CR=critically endangered

|  | Df | Sum Sq | Mean Sq | F value | Pr(>F) |
| --- | --- | --- | --- | --- | --- |
| RedList | 4 | 0.0952396 | 0.0238099 | 0.5580988 | 0.7040464 |
| KnownDecline | 1 | 0.0275537 | 0.0275537 | 0.6458527 | 0.4580865 |
| Kingdom | 1 | 0.0011684 | 0.0011684 | 0.0273876 | 0.8750400 |
| Reproduction | 1 | 0.0003238 | 0.0003238 | 0.0075890 | 0.9339612 |
| Pollination | 1 | 0.0375975 | 0.0375975 | 0.8812784 | 0.3909509 |
| Mobility | 1 | 0.1600627 | 0.1600627 | 3.7518370 | 0.1104995 |
| AreaRange | 1 | 0.0174745 | 0.0174745 | 0.4095989 | 0.5503439 |
| Residuals | 5 | 0.2133125 | 0.0426625 | NA | NA |

While no association between life history and *z_MAR_* was found (Table S13), this may be due to limited power, as the sample size of species analysed here is still small, n=20. Further studies expanding the numbers of species will be necessary to confirm or reject this expected association.

#### V. An estimate of global genetic diversity loss

Using the approach described in section II.4, we generated a number of estimates either per ecosystem or per species. All estimates below tried to be conservative, and thus we always used the scaled *z_MAR_* values (section II.3.2.)

##### V.1 Estimates of ecosystem area losses

**Table S14.**
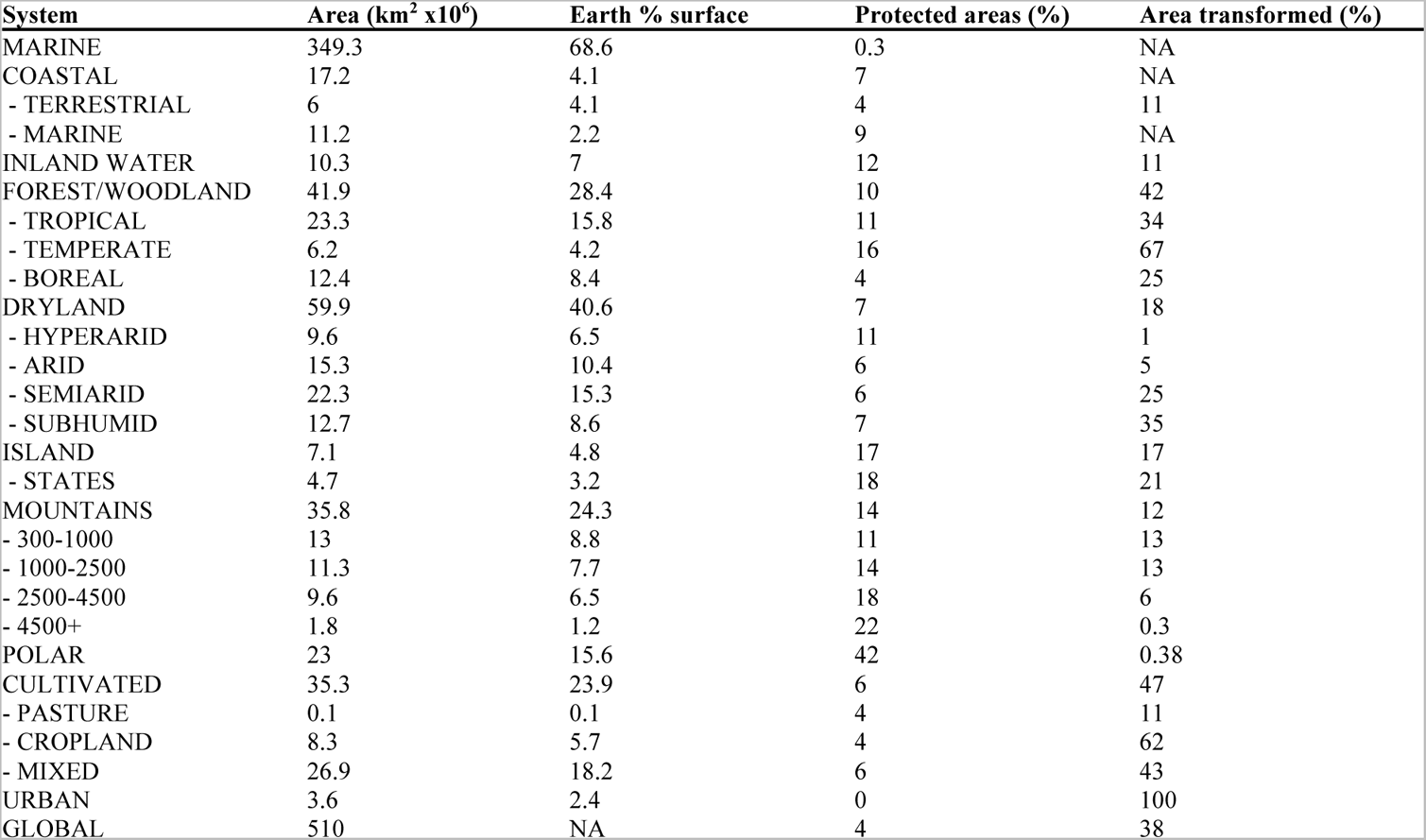
Millennium Ecosystem Assessment land cover transformation. Changes of ecosystem area pre-21st century. Ecosystem names are repeated for ecosystem sub-classes. Source: https://www.millenniumassessment.org

| System | Area (km<sup>2</sup> x10<sup>6</sup>) | Earth % surface | Protected areas (%) | Area transformed (%) |
| --- | --- | --- | --- | --- |
| MARINE | 349.3 | 68.6 | 0.3 | NA |
| COASTAL | 17.2 | 4.1 | 7 | NA |
| - TERRESTRIAL | 6 | 4.1 | 4 | 11 |
| - MARINE | 11.2 | 2.2 | 9 | NA |
| INLAND WATER | 10.3 | 7 | 12 | 11 |
| FOREST/WOODLAND | 41.9 | 28.4 | 10 | 42 |
| - TROPICAL | 23.3 | 15.8 | 11 | 34 |
| - TEMPERATE | 6.2 | 4.2 | 16 | 67 |
| - BOREAL | 12.4 | 8.4 | 4 | 25 |
| DRYLAND | 59.9 | 40.6 | 7 | 18 |
| - HYPERARID | 9.6 | 6.5 | 11 | 1 |
| - ARID | 15.3 | 10.4 | 6 | 5 |
| - SEMIARID | 22.3 | 15.3 | 6 | 25 |
| - SUBHUMID | 12.7 | 8.6 | 7 | 35 |
| ISLAND | 7.1 | 4.8 | 17 | 17 |
| - STATES | 4.7 | 3.2 | 18 | 21 |
| MOUNTAINS | 35.8 | 24.3 | 14 | 12 |
| - 300-1000 | 13 | 8.8 | 11 | 13 |
| - 1000-2500 | 11.3 | 7.7 | 14 | 13 |
| - 2500-4500 | 9.6 | 6.5 | 18 | 6 |
| - 4500+ | 1.8 | 1.2 | 22 | 0.3 |
| POLAR | 23 | 15.6 | 42 | 0.38 |
| CULTIVATED | 35.3 | 23.9 | 6 | 47 |
| - PASTURE | 0.1 | 0.1 | 4 | 11 |
| - CROPLAND | 8.3 | 5.7 | 4 | 62 |
| - MIXED | 26.9 | 18.2 | 6 | 43 |
| URBAN | 3.6 | 2.4 | 0 | 100 |
| GLOBAL | 510 | NA | 4 | 38 |

Ecosystem transformation has been tracked over decades. We extracted ecosystem transformations from the Millennium Ecosystem Assessment (*62*), which estimated ecosystem transformations from presumably native systems to cultivated or urban areas by GLC2000 land cover dataset (Table S14). The forest/woodland is calculated as percentage change between potential vegetation from WWF ecoregions to the current actual forest/woodland areas from GLC2000. These provide bulk ecosystem reductions, not for a given species, but may be a good proxy for an average across species.

**Table S15.** IPBES land cover transformation, Source: https://ipbes.net

| Region | Area(Mkm2) | MSA 2010 | MSA 2050 SSP2 | MSA 2050 SSP1 | MSA 2050 SSP3 |
| --- | --- | --- | --- | --- | --- |
| North America | 20 | 65 | 56 | NA | NA |
| Central and South America | 18 | 65 | 53 | NA | NA |
| Middle East and Northern Africa | 11 | 81 | 77 | NA | NA |
| Sub-Saharan Africa | 24 | 70 | 56 | NA | NA |
| Western and Central Europe | 6 | 37 | 29 | NA | NA |
| Russian region and Central Asia | 21 | 73 | 65 | NA | NA |
| South Asia | 5 | 44 | 35 | NA | NA |
| China region | 11 | 56 | 49 | NA | NA |
| Southeast Asia | 7 | 55 | 43 | NA | NA |
| Japan, Korea and Oceania | 8 | 71 | 57 | NA | NA |
| Polar | 2 | 96 | 91 | NA | NA |
| World | 132 | 66 | 56 | 62 | 54 |

The Intergovernmental Science-Policy Platform on Biodiversity and Ecosystem Services (IPBES) recently used a PBL satellite product from the Netherlands Environmental Assessment Agency (https://www.pbl.nl/en/natureand-biodiversity) to study the % of area ecosystem transformation in the world (Table S15). This provides an updated estimate to the Millennium Assessment as well as projections under several Shared Socioeconomic Pathways (1-3) for 2050. These were reported per region as of 2010, and for projections to 2050 (scenario SSP2). Instead of direct area, the metric is a composite of land use information to predict Mean Species Abundance (MSA), a measure of the size of populations of wild organisms as a percentage of their inferred abundance in their natural state (% MSA).

A global transformation metric can also be captured by the most updated land use transformation data, the Land Use Harmonization 2 (release v2e for 2015-2011 and release v2h for baseline 1850-2015) (*63*). Baseline transformation of primary ecosystems was calculated subtracting the total area covered by primary forest (primf) and primary non-forest (primn) variables between year 1850 layer (roughly pre-industrial baseline) and the present, 2015, as *1-A_2015_ / A_1850_* (Table S16). Analyses that use projections to mid-21^st^ century were conducted similarly as in (*64*), summing over all transitions from primary forest (primf), primary non-forest (primn), secondary forest (secdf) and secondary non-forest (secdn) lands to any other category for all years within the 2015-2050 period (see Table S10).

**Table S16.** Land Use Harmonization 2 from 1850 to 2015. Source: https://luh.umd.edu/data.shtml

|  | Area % |
| --- | --- |
| Primary forest transformed | 43 |
| Primary non-forest transformed | 50 |

Finally, we searched for timely estimates of forest reduction (based on vegetation cover) reported in the Global Forest Watch website: https://www.globalforestwatch.org/dashboards/global/ (accessed June 2021). From 2002 to 2020, there has been a global tree cover loss of 10%, with an annual tree cover loss of 0.6-1.1%.

Although these are not direct area transformations, we also used the IUCN Red List resource (https://www.iucnredlist.org, Table S12 shows status of the species analysed here), which includes guides to categorise species as vulnerable, endangered, critically endangered, and extinct, and has conducted extensive assessments across thousands of species (Table S17).

**Table S17.** IUCN Red List categories of extinction risk and number of species. Source: www.iucnredlist.org, January 2021

| IUCN Red List | Description | Criterion of area or pop. | # plant species |
| --- | --- | --- | --- |

| Category |  | reduction (>%) |  |
| --- | --- | --- | --- |
| EX | Extinct | 100 | 164 |
| EW | Extinct in the Wild | 100 |  |
| CR | Critically Endangered | 80 | 4674 |
| EN | Endangered | 50 | 8593 |
| VU | Vulnerable | 30 | 8459 |
| NC, LR, NT, DD, LC | No Concern, Low Risk, Near Threatened,<br>Data Deficient, Least Concern, Other | 0 | 32237 |

##### V.2 A global estimate of genetic loss

Taking the estimates and standard error of *z_MAR_* across species, and the world’s reduction of ecosystems we can calculate the fraction of genetic diversity reduction following the MAR equation (section II.4), giving a range of estimates (Table S18).

**Fig. S23.**
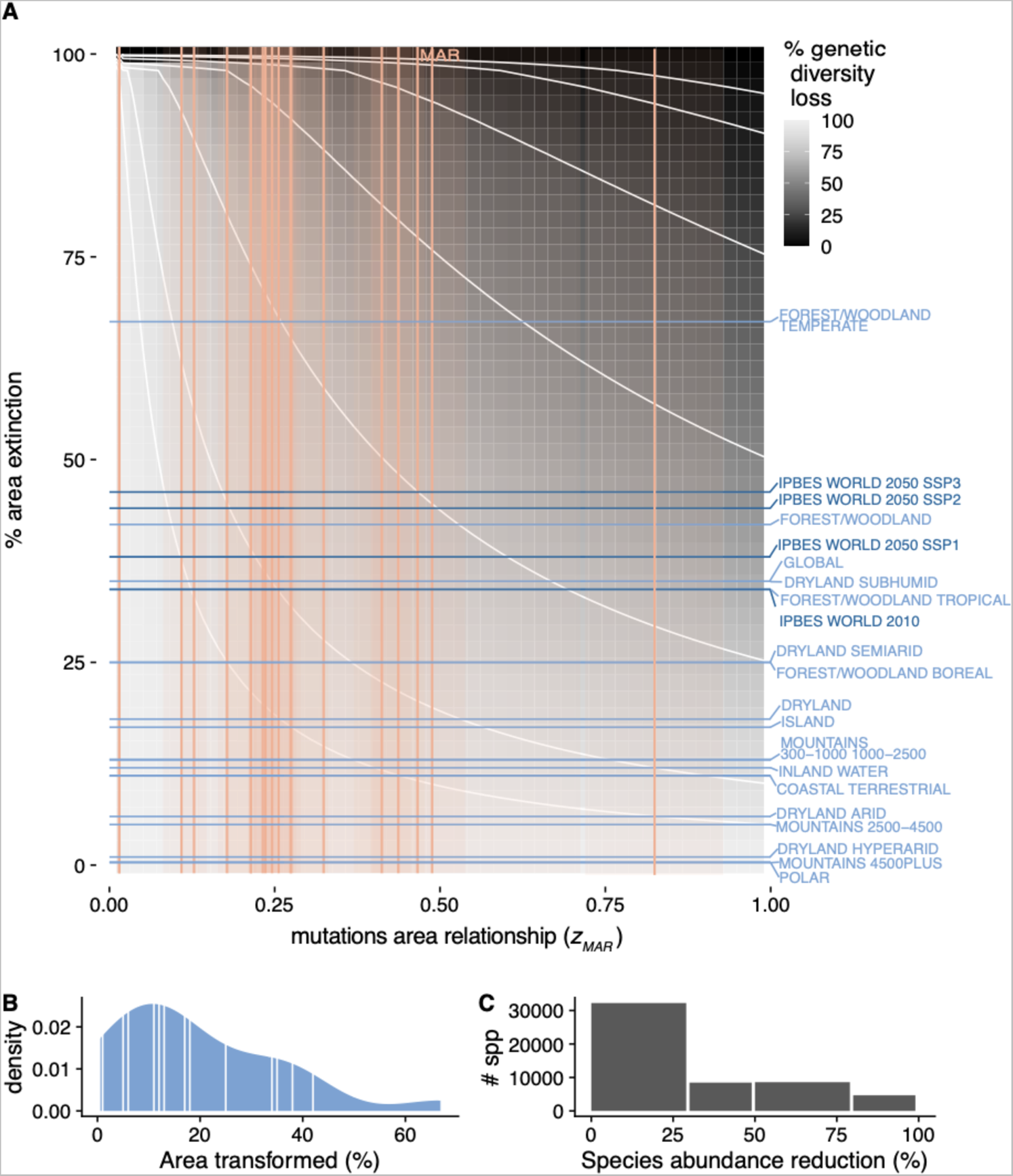
The parameter space of genetic diversity loss, extended. (A) The theoretical space of genetic diversity loss. z_MAR_ values (using area, unscaled for samples, differently from Fig. 32) computed for species analyzed here are marked as orange vertical lines, with confidence intervals as orange shading. Blue horizontal lines correspond to ecosystem transformations from the Millennium Assessment (light blue) and IPBES Assessment (dark blue) (B) Density histogram of percentages of area transformed across ecosystems from the MA, with averages per ecosystem marked in the distribution as well as horizontal lines in (A). (C) The number of species of each of the IUCN categories and the most optimistic range of area or abundance reduction for each of the category brackets.

**Table S18.**
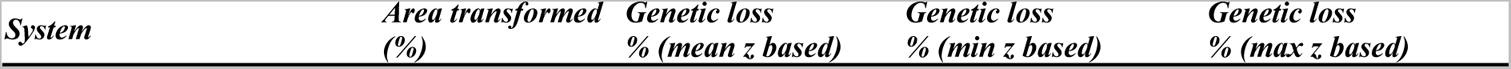
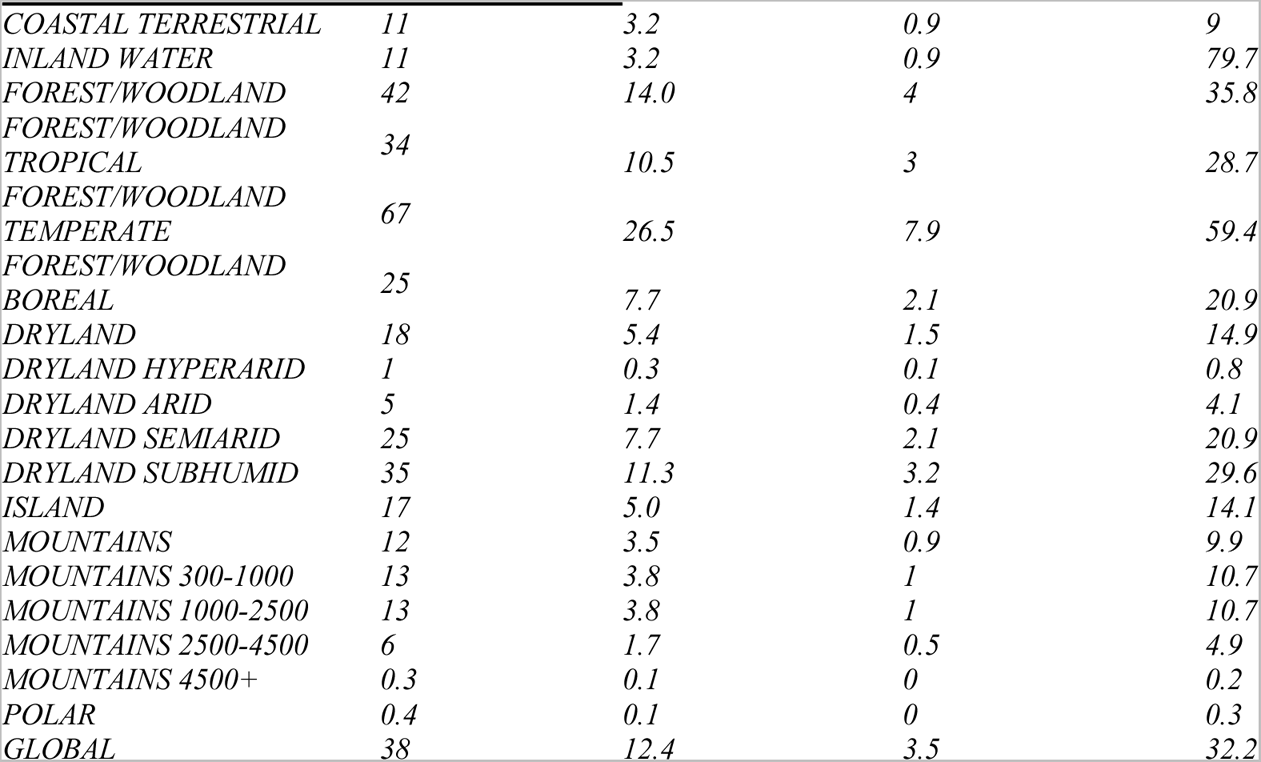
Estimates of average expected genetic loss for different ecosystems. Assuming ecosystem transformation approximately translates into average species distribution reduction, and using the ranges of z_MAR_ from Table 1 of the main text, we project the average genetic loss using the Mutations Area Relationship.

Assuming the average *z_MAR_*, and utilising tree cover from the Global Forest Watch (https://www.globalforestwatch.org), which estimates 0.6-1.1% of transformation per year across Canada, United States and Australia, we extrapolated genetic diversity loss in the next 50 years for tree species to be 8-15% genetic diversity loss.

Assuming that the calculated *z_MAR_* estimates (Table 1) are representative of plant species, we conducted an experiment to create a distribution of % of genetic diversity loss in threatened species. We used the number of species in each IUCN category (Table S17) for a total of 54,127 plant species. For plant species, one of the evaluation criteria of percentage of population loss likely translates faithfully to area reduction in the species. Thus, the proportion of species per category gives a discrete probability distribution of the ranges of percentage of area loss: P(0-29%)=0.596, P(30-49%)=0.156, P(50-79%)=0.159, P(80-99%)=0.086, P(99%-100%)=0.003. Using a simulation-based sampling approach, we drew 350,000 random area reductions *A_t_ / A_t-1_* from the previous distribution and a *z_MAR_* from the mean and variance of our estimates from Table 1 for plants. These were plugged into the MAR equation (Section II.4) to calculate the percentage of genetic diversity loss of these 350,000 random draws. The resulting distribution had a median and interquartile range of 17.53 % [7.51-31.82].

Using the Land Use Harmonization 2 dataset, we also create per-species predictions based on the % transformation of each of the sampled regions per species (Table S14). As before, the land use transformations that merit be considered area losses are all transitions from primary forest (primf), primary non-forest (primn), secondary forest (secdf) and secondary non-forest (secdn) lands to any other category. Taking all the locations where each species has been sampled, we extracted the predicted % of land use change per cell and summed over all cells where individuals had been sampled (we call this LUH^2^ change ‘50, see column in Table S10). We also produced the alternative area loss estimate taking that at least 10% predicted habitat transformation for a grid cell renders the entire area of that grid cell as impacted or lost (we call this LUH^2^ >10% change ‘50). These per-species area losses, in combination with the matched *z_MAR_*, provided a range of potential loss estimates to 2050 ranging 0-36% depending on the species (Table S10).

##### V.3 Community ecology simulations and MAR

To test whether intermediate levels of MAR would be expected across species in entire ecosystems, we conducted community assembly simulations of ~100-500 species following the Neutral Theory of Biodiversity (*1, 41*) and coalescent simulations (*23*) using the software MESS (*65*). These simulations are computationally demanding and could not run in a complete 2D spatial grid. Instead, they were simulated in a mainland-island system, with islands of increasing areas. The community forms by species colonising an empty island according to Hubbell’s Unified Neutral Theory of Biodiversity and Biogeography (UNTB), where all species are equally likely to colonise and persist in the local community. Continued colonisation and migration to the local community continues to bring in new species that may or may not survive, while also continuously bringing in individuals of species already in the local community. The community assembly process ends when the community has reached an equilibrium denoted as the balance between local extinction and new species dispersing into the area (Hubbell 2001). Once the forward-time process has ended, we simulate the coalescent history of each species backward in time. For this, MESS considers the population size, divergence time, and migration rates of the meta and local communities. These coalescent simulations provide us with genetic data and ultimately diversity estimates for each species in the community.

We simulated 100 MESS communities, and for each community the size of the local community was varied from 1K to 100K. We varied the size of communities to emulate variation in area occupied by a given community because we assume as the number of individuals in a community increases from 1,000 to 100,000, so does the area occupied. All other parameters were kept consistent across each of these community simulations, and most remained at their default value. The parameters changed were the length of the sequences simulated for the coalescent-based simulations, which was fixed at 10,000 bp, and the migration rate, which was fixed at 0.01.

The simulation output was used to then compute a single *z_SAR_* for the system as *S=cA^zSAR^,* and one *z_MAR_* for each species in the same way, *M=cA^zMAR^*. This resulted in the distribution of *z_MAR_* from Fig. S24. This confirmed that we can recover typical *z_SAR_* and *z_MAR_* values from completely stochastic neutral yet spatially structured systems such as species in communities and mutations in populations of a species.

**Fig. S24.**
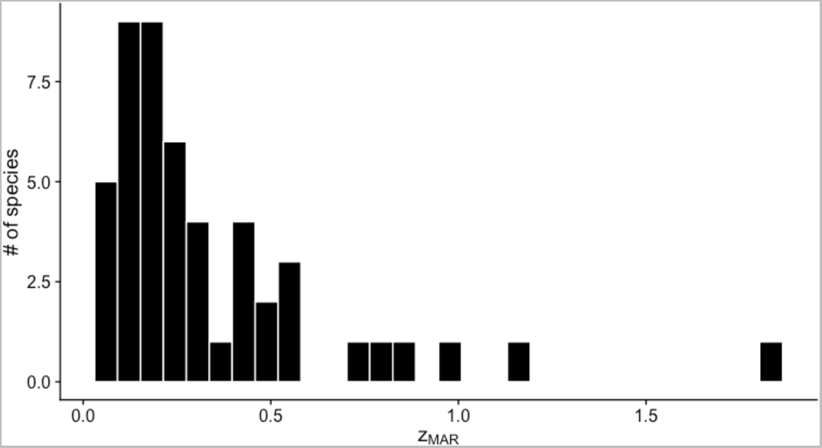
z_MAR_ calculated from MESS eco-evolutionary simulations. Using the MESS framework of a mainland-island model with different island sizes, z_MAR_ per species is recovered. The stochastic nature of the simulations results in each species having different abundances and migration histories that change the scaling value. Values were typically around 0.3. Rarely some species had values above 1, which appear could be noisy estimates from recently colonising species in the simulations.

##### V.4 The nested species extinction and genetic diversity loss processes

Finally, we worried that our estimates of V.2 would be mistaken as overestimates. In fact, we believe these may be underestimated. Recent policy proposals for the United Nations’ Sustainability Goals emphasize that the target of protecting 90% of species genetic diversity for all species cannot leave the already-extinct species behind (*66*) (That is, one cannot protect 90% of species and leave 10% to become extinct to meet this goal). This clearly exemplifies a problem in conservation biology that what researchers can study is (most of the time) what has escaped extinction, and therefore if we do not account for extinct species in our overall estimates of genetic diversity loss we may naively think ecosystems have not suffered genetic diversity loss (i.e. in the extreme scenario, an ecosystem that has lost all but one abundant species may not really appear genetically eroded if such species is in good shape).

We then created spatial simulations in R where 1,000 species are distributed in 100×100 grid cells following a UNTB abundance distribution and then proceeded with an edge extinction of the ecosystem (see Fig. S25 for a cartoon).

**Fig. S25.**
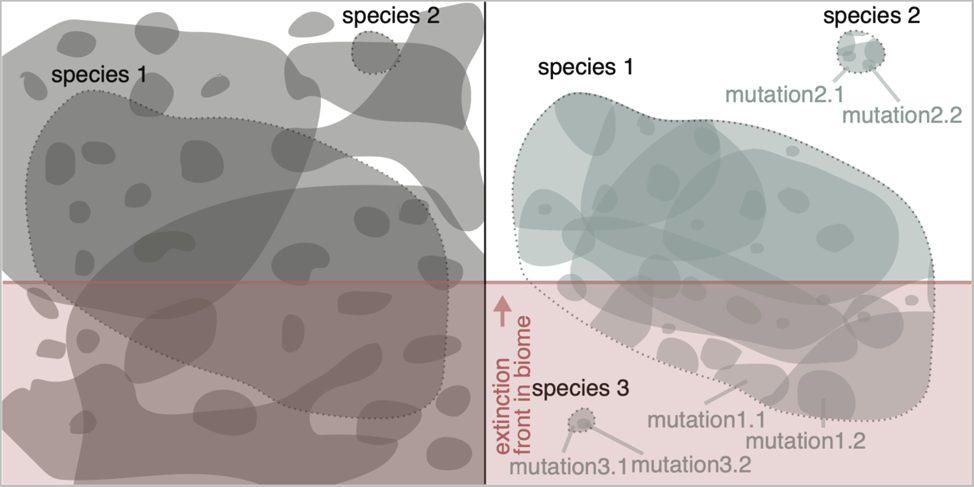
Cartoon of nested extinction of species and genetic diversity loss. An ecosystem with multiple species within it (left), distributed in space, with few species broadly distributed and many narrowly distributed. Moving one level of biological organization lower, mutations within species (right) are also spatially distributed with many narrowly distributed. As extinction happens (red line moving bottom to top), all species below the red line go extinct, but only the mutations within species 1 below the line are lost, while mutations above the line remain. Species 3 has already become extinct, and therefore also all the mutations within it.

Two extreme types of distributions of species can be imagined: species are randomly placed in space, or species are found mostly in perfectly contiguous ranges (We ended up using as an example a simulation with 85% of the individuals of a species found in a core square continuous distribution and 15% found outside that core in fragmented observations, as this scenario produced the canonical SAR of *z*~0.3). Spatial structure interestingly creates two extreme distributions of area reductions across species (Fig. S26): random placement of cell habitats essentially show that the average area reduction per ecosystem is followed by most species, while autocorrelated placement of cell habitats create a U distribution in area reductions, where at the beginning of the extinction process most species have not experienced any impact (Fig. S26B left) but at the end of ecosystem reduction virtually all species are already extinct (Note we may be at the beginning of S26B process given the data from IUCN, Fig 3C).

**Fig. S26.**
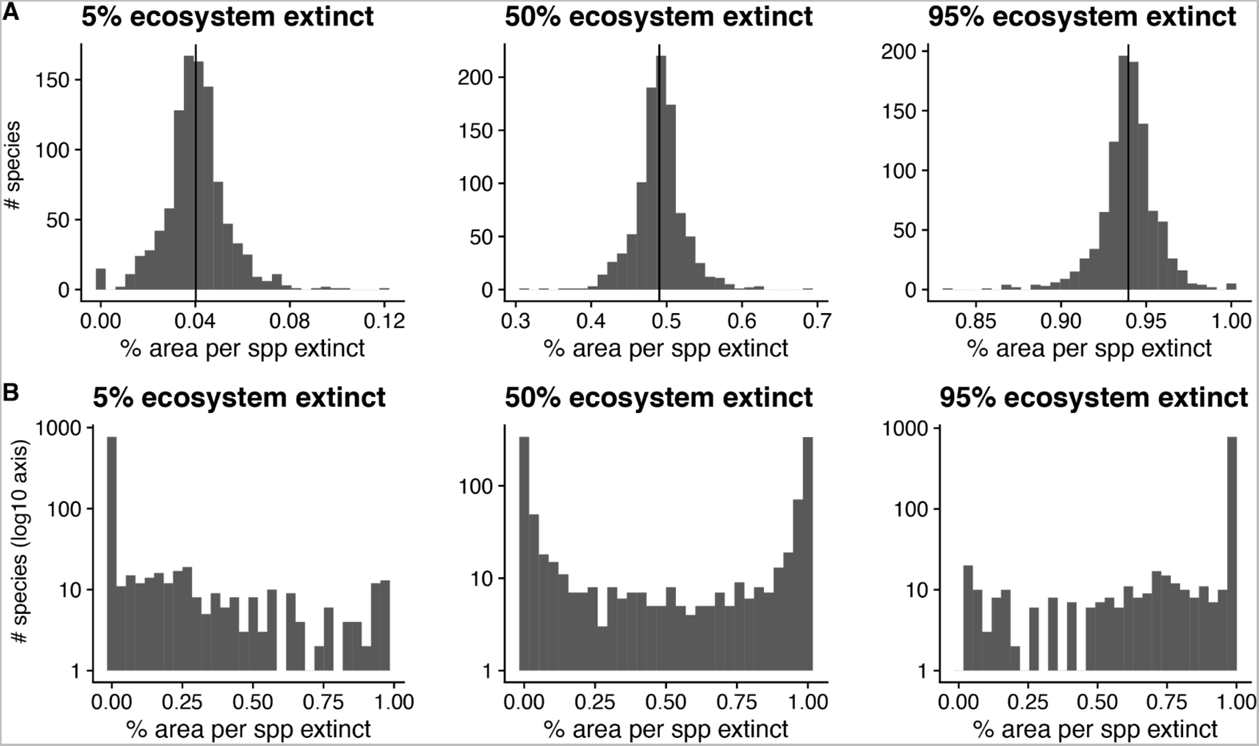
The distribution of per-species area lost and total ecosystem extinction with 1000 species. Two ecosystems of 100×100 cells with 1000 species. Species are either randomly distributed in cells (A) or spatially autocorrelated with occupying mostly contiguous cells (B). As the extinction process wipes out part of the ecosystem (snapshots are provided at 5%, 50%, and 95%), the area loss per species (and hence genetic diversity lost) is tracked. In (A) the average area lost per species is roughly the total reduction of the ecosystem, whereas in (B) the distribution is U shaped (note the log-scaled y-axis). While in (B) the mean area lost in the distribution correctly captures the area loss of the ecosystem, per species losses are highly uneven.

To study the consequence of the above differential area loss and the effect of some species going extinct on the total ecosystem genetic diversity, we conducted the next analysis: For extant species, we assumed they would lose genetic diversity following the MAR relationship (section II.4), with all species having *z_MAR_ = 0.3* for simplicity (i.e. all species lose genetic diversity at the same rate). For extinct species (100% of their area reduced), we considered genetic diversity loss was 100%. The compound total genetic diversity loss would then just be the sum of those 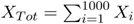 (Of course, in reality species may vary in their genome-wide diversity average, and we could for instance use Watterson’s Θ*_W_* (see section II.2) to scale the total loss of genetic diversity in the ecosystem accounting for different basal level of diversity per species: 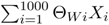). Interestingly, if we calculate the *z* of the slope of compound genetic diversity across species in an ecosystem it is much larger than MAR or SAR alone: *z_compunded_ =* 0.6 (Fig. S27).

**Fig. S27.**
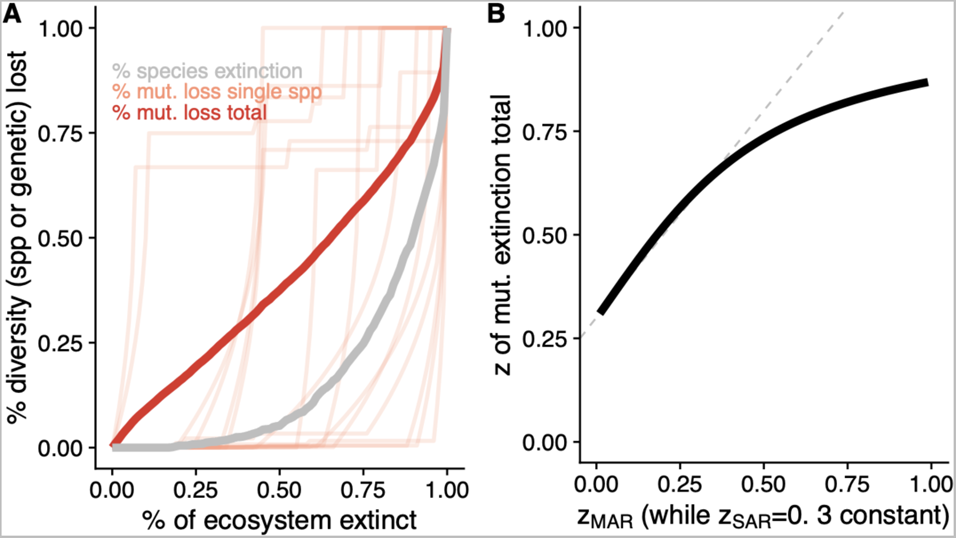
Numeric simulation of nested species and genetic diversity loss. (A) Simulating the extinction of an ecosystem with 1,000 species that follow a log-normal species abundance curve. Extinction of the ecosystem creates a curve of species loss of z~0.3 (grey). Likewise, each species trajectory (light red, 15 species drawn randomly) follows a simulated genetic diversity loss of z_MAR_~0.3 as they lose area. Because species’ geographic distributions are by construction smaller than the whole ecosystem area, those distributed closer to the start of the extinction front lose area first, while those distributed farthest from the extinction front only lose area when the ecosystem is almost completely destroyed. Because genetic diversity loss is both due to complete extinction of species as well as area reduction of extant species, the compound genetic diversity loss curve (red) follows the faster loss dynamics. (B) Holding z_SAR_=0.3 constant, and varying z_MAR_ in independent simulations shows that the compound genetic diversity across species is close to the sum of both z slopes (the SAR and the MAR), but it saturates at ca. 0.85 (grey dotted line shows z_MAR_ + z_SAR_).

#### VI. Limitations and outlook

In this last section we list some potential limitations of an inherently simple scaling law, and what approaches could be used to address those and improve genetic diversity loss projections.

##### VI.1 Reasons for overestimations

Many researchers have posited that SAR likely overestimates species extinction (*33, 67*). For instance:

- Ignoring that a diversity-area relationship can be defined outwards, inwards, or focusing on endemisms can have an impact (*10, 33, 67*). To address this, we confirmed relative consistency between inward, outward, and random placement MAR, and proposed that the EMAR may not be that appropriate to study genetic diversity loss (or at least EMAR does not show predictability in our simulation).
- Species may persist in altered habitats, like some animals are known to do (*68*). We have focused some of the estimates in this study on plants, for which area loss should equate to population loss and vice versa, but further extensions could be applied in the future as described by Pereira and Daily (*68*).
- SAR is not a mechanistic model (*69*). We have derived its ranges of possible values and averages analytically and are beginning to understand how evolutionary forces shape MAR. Realistic simulations can help understand in a process-based framework how populations (and their MAR) react to partial population extinction (continuous space simulations with progressive area reductions appear to fit well with the MAR predictions calculated before the extinction process starts, section III.2.6).
- There is a scale dependence in the SAR slope, with slight increase in the slope at large scales (*10*). Since power laws are typically fit with large-scale datasets and used to predict local scale extinctions, predictions could be overestimated at local scales.

##### VI.2 Reasons for underestimations

While the simplicity of power laws to make predictions of species extinction may lead to overestimations, there are also important reasons to believe MAR would underestimate genetic loss.

- Perhaps even more so than in species list datasets and census, the discovery of low frequency genetic variants is highly underpowered (*70*). These are highly prevalent, but genomic pipelines, with the aim to be conservative, often filter out rare variants. This would underestimate *z_MAR_* and therefore the degree of genetic diversity loss with area shrinkage. This is clear in the pre-selected-only marker dataset of *Pinus contorta*.
- Related to the previous: Although sequencing methods have an error rate that misreads true nucleotide sequences, this rate is typically extremely low (many sequencing projects described here used Illumina HiSeq series, which has a 0.112% error rate, or about 1 misread nucleotide in 1000). This could intuitively lead to overestimates in mutations in space but in fact, the mis-reading of DNA ends up causing an underestimation. This is because bioinformatic software that transforms raw data into SNP variant tables errs towards the conservative direction, often not calling mutations that have been observed very few times, and thus likely under-representing rare mutations (*71*).
- The use of scaled *z_MAR_* proposed in section II.3.2. accounts for that the minimum *z_MAR_* is rarely exactly 0, especially when sample sizes are limited. We use this correction scaling down *z_MAR_* to be conservative. However, *z_MAR_* could only in very exceptional circumstances be 1, but we do not correct for this, again, to have a conservatively low *z_MAR_.* Hence, our conservative approach would generally lead to underestimates of genetic diversity loss.
- When species shrink in area, the effective population size of the remaining population decreases, increasing drift and moving towards a lower diversity equilibrium. This reactive process is not captured by the phenomenological MAR relationship.
- The nested extinction of species and genetic diversity loss (section V.3) would lead us, by the right of “survival bias”, to underestimate how much genetic diversity has been lost cumulative in an ecosystem.

##### VI.3 Final notes

Ultimately, to make accurate predictions of genetic diversity loss and increased extinction risk of species, very detailed data and expert assessment per species will be required: census sizes, genome size, migration in metapopulations, mating system, detailed maps of genetic makeups, and finescale area transformations. This could enable mechanistic models projected forward-in-time such as discussed in section II.3.6. The production of new genomic datasets across entire ecosystems should further help create maps of genetic diversity at high resolution to track losses (*72–74*).

Our philosophy in this work has been to err on the conservative side when projecting genetic diversity loss (e.g. using area calculations that produce lower *z_MAR_* values, scaling them for low sample bias, using lower estimates of ecosystem transformation, etc.). However, this conservative approach can also lead us into under-estimating loss. As described in V.4., the phenomenon of survival bias likely leads us to underestimate what has been lost given we do not observe it. A phenomenon also highlighted as a possible explanation for the relatively shy difference in genetic diversity between threatened and non-threatened species (*75, 76*)

Because to our knowledge, no other approaches exist to project genetic diversity, we believe that MAR is a quantitative and scalable first-approximation of genetic diversity that would just require accurate understanding of abundance or area reductions and minimal information about population structure or mating/dispersal/range relationships. Given that scaling relationships are already applied by conservation policy (*77*), and given that assumptions and limitations are understood, we expect MAR to become a relevant tool to project losses of a dimension of biodiversity so far mostly invisible or unaddressable in large conservation projections.

## REFERENCES

1. IUCN, The IUCN Red List of Threatened Species. https://www.iucnredlist.org.

2. S. Díaz, J. Settele, E. S. Brondízio, H. T. Ngo, J. Agard, A. Arneth, P. Balvanera, K. A. Brauman, S. H. M. Butchart, K. M. A. Chan, L. A. Garibaldi, K. Ichii, J. Liu, S. M. Subramanian, G. F. Midgley, P. Miloslavich, Z. Molnár, D. Obura, A. Pfaff, S. Polasky, A. Purvis, J. Razzaque, B. Reyers, R. R. Chowdhury, Y.-J. Shin, I. Visseren-Hamakers, K. J. Willis, C. N. Zayas, Pervasive human-driven decline of life on Earth points to the need for transformative change. Science. 366, eaax3100 (2019).

3. IPBES, Global Assessment Report on Biodiversity and Ecosystem Services (IPBES Secretariat, Bonn, Germany, 2019; https://www.ipbes.net/global-assessment-report-biodiversity-ecosystem-services).

4. J. J. Wiens, Climate-Related Local Extinctions Are Already Widespread among Plant and Animal Species. PLoS Biol. 14, e2001104 (2016).

5. W. Thuiller, S. Lavorel, M. B. Araújo, M. T. Sykes, I. C. Prentice, Climate change threats to plant diversity in Europe. Proc. Natl. Acad. Sci. U. S. A. 102, 8245–8250 (2005).

6. M. Exposito-Alonso, F. Vasseur, W. Ding, G. Wang, H. A. Burbano, D. Weigel, Genomic basis and evolutionary potential for extreme drought adaptation in Arabidopsis thaliana. Nat Ecol Evol. 2, 352–358 (2018).

7. C. Parmesan, Ecological and Evolutionary Responses to Recent Climate Change. Annu. Rev. Ecol. Evol. Syst. 37, 637–669 (2006).

8. T. Capblancq, M. C. Fitzpatrick, R. A. Bay, M. Exposito-Alonso, S. R. Keller, Genomic Prediction of (Mal)Adaptation Across Current and Future Climatic Landscapes. Annu. Rev. Ecol. Evol. Syst. 51, 245–269 (2020).

9. M. Lynch, J. Conery, R. Burger, Mutation Accumulation and the Extinction of Small Populations. Am. Nat. 146, 489–518 (1995).

10. D. Spielman, B. W. Brook, R. Frankham, Most species are not driven to extinction before genetic factors impact them. Proceedings of the National Academy of Sciences. 101 (2004), pp. 15261–15264.

11. W. Steffen, K. Richardson, J. Rockström, S. E. Cornell, I. Fetzer, E. M. Bennett, R. Biggs, S. R. Carpenter, W. de Vries, C. A. de Wit, C. Folke, D. Gerten, J. Heinke, G. M. Mace, L. M. Persson, V. Ramanathan, B. Reyers, S. Sörlin, Planetary boundaries: Guiding human development on a changing planet. Science. 347 (2015), doi:10.1126/science.1259855.

12. S. Díaz, N. Zafra-Calvo, A. Purvis, P. H. Verburg, D. Obura, P. Leadley, R. Chaplin-Kramer, L. De Meester, E. Dulloo, B. Martín-López, M. R. Shaw, P. Visconti, W. Broadgate, M. W. Bruford, N. D. Burgess, J. Cavender-Bares, F. DeClerck, J. M. Fernández-Palacios, L. A. Garibaldi, S. L. L. Hill, F. Isbell, C. K. Khoury, C. B. Krug, J. Liu, M. Maron, P. J. K. McGowan, H. M. Pereira, V. Reyes-García, J. Rocha, C. Rondinini, L. Shannon, Y.-J. Shin, P. V. R. Snelgrove, E. M. Spehn, B. Strassburg, S. M. Subramanian, J. J. Tewksbury, J. E. M. Watson, A. E. Zanne, Set ambitious goals for biodiversity and sustainability. Science. 370, 411–413 (2020).

13. CBD, 1st Draft of The Post-2020 Global Biodiversity Framework (2021), (available at https://www.cbd.int/doc/c/abb5/591f/2e46096d3f0330b08ce87a45/wg2020-03-03-en.pdf).

14. D. M. Leigh, A. P. Hendry, E. Vázquez-Domínguez, V. L. Friesen, Estimated six per cent loss of genetic variation in wild populations since the industrial revolution. Evol. Appl. 12, 1505–1512 (2019).

15. K. L. Millette, V. Fugère, C. Debyser, A. Greiner, F. J. J. Chain, A. Gonzalez, No consistent effects of humans on animal genetic diversity worldwide. Ecol. Lett. 23, 55–67 (2020).

16. I. G. Alsos, D. Ehrich, W. Thuiller, P. B. Eidesen, A. Tribsch, P. Schönswetter, C. Lagaye, P. Taberlet, C. Brochmann, Genetic consequences of climate change for northern plants. Proc. Biol. Sci. 279, 2042–2051 (2012).

17. S. Theodoridis, C. Rahbek, D. Nogues-Bravo, Exposure of mammal genetic diversity to mid-21st century global change. Ecography (Cop.) (2021), doi:10.1111/ecog.05588.

18. O. Arrhenius, Species and Area. J. Ecol. 9, 95–99 (1921).

19. D. Storch, P. Keil, W. Jetz, Universal species-area and endemics-area relationships at continental scales. Nature. 488, 78–81 (2012).

20. C. D. Thomas, A. Cameron, R. E. Green, M. Bakkenes, L. J. Beaumont, Y. C. Collingham, B. F. N. Erasmus, M. F. de Siqueira, A. Grainger, L. Hannah, L. Hughes, B. Huntley, A. S. van Jaarsveld, G. F. Midgley, L. Miles, M. A. Ortega-Huerta, A. Townsend Peterson, O. L. Phillips, S. E. Williams, Extinction risk from climate change. Nature. 427, 145–148 (2004).

21. V. Buffalo, Quantifying the relationship between genetic diversity and population size suggests natural selection cannot explain Lewontin’s paradox. Elife. 10 (2021), doi:10.7554/eLife.67509.

22. R. A. Fisher, XVII.—The Distribution of Gene Ratios for Rare Mutations. Proceedings of the Royal Society of Edinburgh. 50, 204–219 (1931).

23. 1001 Genomes Consortium, 1,135 Genomes Reveal the Global Pattern of Polymorphism in Arabidopsis thaliana. Cell. 166, 481–491 (2016).

24. F. W. Preston, The canonical distribution of commonness and rarity: Part I. Ecology. 43, 185 (1962).

25. A. Hampe, R. J. Petit, Conserving biodiversity under climate change: the rear edge matters. Ecol. Lett. 8, 461–467 (2005).

26. V. Buffalo, Why do species get a thin slice of π? Revisiting Lewontin’s Paradox of Variation. bioRxiv (2021), p. 2021.02.03.429633.

27. Millennium Ecosystem Assessment, Millennium ecosystem assessment (Millennium Ecosystem Assessment, 2005; http://chapter.ser.org/europe/files/2012/08/Harris.pdf).

28. Ipbes, Global assessment report of the Intergovernmental Science-Policy Platform on Biodiversity and Ecosystem Services (IPBES Secretariat, Bonn, Germany, 2019; https://www.ipbes.net/news/ipbes-global-assessment-summary-policymakers-pdf).

29. G. C. Hurtt, L. Chini, R. Sahajpal, S. Frolking, B. L. Bodirsky, K. Calvin, J. C. Doelman, J. Fisk, S. Fujimori, K. Klein Goldewijk, T. Hasegawa, P. Havlik, A. Heinimann, F. Humpenöder, J. Jungclaus, J. O. Kaplan, J. Kennedy, T. Krisztin, D. Lawrence, P. Lawrence, L. Ma, O. Mertz, J. Pongratz, A. Popp, B. Poulter, K. Riahi, E. Shevliakova, E. Stehfest, P. Thornton, F. N. Tubiello, D. P. van Vuuren, X. Zhang, Harmonization of global land use change and management for the period 850–2100 (LUH2) for CMIP6. Geosci. Model Dev. 13, 5425–5464 (2020).

30. G. Gibson, Rare and common variants: twenty arguments. Nat. Rev. Genet. 13, 135–145 (2011).

31. M. V. Rockman, The QTN program and the alleles that matter for evolution: all that’s gold does not glitter. Evolution. 66, 1–17 (2012).

32. M. J. Harms, J. W. Thornton, Evolutionary biochemistry: revealing the historical and physical causes of protein properties. Nat. Rev. Genet. 14, 559–571 (2013).

33. M. Exposito-Alonso, 500 Genomes Field Experiment Team, H. A. Burbano, O. Bossdorf, R. Nielsen, D. Weigel, Natural selection in the *Arabidopsis thaliana* genome in present and future climates. Nature. 573, 126–129 (2019).

34. H. R. Taft, D. A. Roff, Do bottlenecks increase additive genetic variance? Conserv. Genet. 13, 333–342 (2012).

35. P. Ehrlich, B. Walker, Rivets and redundancy. Bioscience. 48, 387 (1998).

36. C. C. Kyriazis, R. K. Wayne, K. E. Lohmueller, Strongly deleterious mutations are a primary determinant of extinction risk due to inbreeding depression. Evolution Letters. n/a (2020), doi:10.1002/evl3.209.

37. M. Kardos, E. Armstrong, S. W. Fitzpatrick, S. Hauser, P. Hedrick, J. Miller, D. Tallmon, W. Chris Funk, The crucial role of genome-wide genetic variation in conservation. bioRxiv (2021), p. 2021.07.05.451163.

38. R. Lande, Risks of Population Extinction from Demographic and Environmental Stochasticity and Random Catastrophes. Am. Nat. 142, 911–927 (1993).

39. M. Lynch, R. Lande, The critical effective size for a genetically secure population. Anim. Conserv. 1, 70–72 (1998).

## SUPPLEMENTAL REFERENCES

1. S. P. Hubbell, The Unified Neutral Theory of Biodiversity and Biogeography (Monographs in Population Biology, 2001).

2. M. Tokeshi, Niche Apportionment or Random Assortment: Species Abundance Patterns Revisited. J. Anim. Ecol. 59, 1129–1146 (1990).

3. M. Tokeshi, in Advances in Ecological Research, M. Begon, A. H. Fitter, Eds. (Academic Press, 1993; https://www.sciencedirect.com/science/article/pii/S0065250408600422), vol. 24, pp. 111–186.

4. MOTOMURA, I, A statistical treatment of ecological communities. Zoological Magazine. 44, 379–383 (1932).

5. R. H. MacArthur, On the relative abundance of bird species. Proc. Natl. Acad. Sci. U. S. A. 43, 293–295 (1957).

6. R. A. Fisher, A. S. Corbet, C. B. Williams, The Relation Between the Number of Species and the Number of Individuals in a Random Sample of an Animal Population. J. Anim. Ecol. 12, 42–58 (1943).

7. F. W. Preston, The canonical distribution of commonness and rarity: Part I. Ecology. 43, 185 (1962).

8. F. W. Preston, The commonness, and rarity, of species. Ecology. 29, 254–283 (1948).

9. D. Alonso, A. J. McKane, Sampling Hubbell’s neutral theory of biodiversity. Ecol. Lett. 7, 901–910 (2004).

10. D. Storch, P. Keil, W. Jetz, Universal species-area and endemics-area relationships at continental scales. Nature. 488, 78–81 (2012).

11. S. L. Pimm, G. J. Russell, J. L. Gittleman, T. M. Brooks, The future of biodiversity. Science. 269, 347–350 (1995).

12. C. D. Thomas, A. Cameron, R. E. Green, M. Bakkenes, L. J. Beaumont, Y. C. Collingham, B. F. N. Erasmus, M. F. de Siqueira, A. Grainger, L. Hannah, L. Hughes, B. Huntley, A. S. van Jaarsveld, G. F. Midgley, L. Miles, M. A. Ortega-Huerta, A. Townsend Peterson, O. L. Phillips, S. E. Williams, Extinction risk from climate change. Nature. 427, 145–148 (2004).

13. J. F. C. Kingman, The coalescent. Stochastic Process. Appl. 13, 235–248 (1982).

14. R. C. Griffiths, S. Tavaré, The age of a mutation in a general coalescent tree. Communications in Statistics. Stochastic Models. 14, 273–295 (1998).

15. R. A. Fisher, XVII.—The Distribution of Gene Ratios for Rare Mutations. Proceedings of the Royal Society of Edinburgh. 50, 204–219 (1931).

16. M. W. Hahn, Molecular Population Genetics (Oxford University Press, 2018; https://play.google.com/store/books/details?id=3BDkswEACAAJ).

17. V. Buffalo, Quantifying the relationship between genetic diversity and population size suggests natural selection cannot explain Lewontin’s paradox. Elife. 10 (2021), doi:10.7554/eLife.67509.

18. D. R. Marshal, A. D. H. Brown, in Crop genetic resources for today and tomorrow, O. H. Frankel, J. G. Hawkes, Eds. (Cambridge University Press, 1975; https://www.researchgate.net/publication/280057199), vol. 2.

19. S. Wright, Isolation by Distance. Genetics. 28, 114–138 (1943).

20. J. Novembre, M. Stephens, Interpreting principal component analyses of spatial population genetic variation. Nat. Genet. 40, 646–649 (2008).

21. H. Fan 樊海英, Q. Zhang 张清臣, J. Rao 饶娟娟, J. Cao 曹静文, X. Lu 卢欣, Genetic Diversity-Area Relationships across Bird Species. Am. Nat. 194, 736–740 (2019).

22. P. R. A. Campos, V. M. de Oliveira, A. Rosas, Epistasis and environmental heterogeneity in the speciation process. Ecol. Modell. 221, 2546–2554 (2010).

23. J. Kelleher, A. M. Etheridge, G. McVean, Efficient Coalescent Simulation and Genealogical Analysis for Large Sample Sizes. PLoS Comput. Biol. 12, e1004842 (2016).

24. B. C. Haller, P. W. Messer, SLiM 3: Forward Genetic Simulations Beyond the Wright–Fisher Model. Mol. Biol. Evol. 36, 632–637 (2019).

25. T. R. Booker, S. Yeaman, M. C. Whitlock, The WZA: A window-based method for characterizing genotype-environment association. bioRxiv (2021), p. 2021.06.25.449972.

26. J. Kelleher, K. R. Thornton, J. Ashander, P. L. Ralph, Efficient pedigree recording for fast population genetics simulation. PLoS Comput. Biol. (2018), p. e1006581.

27. B. C. Haller, J. Galloway, J. Kelleher, P. W. Messer, P. L. Ralph, Tree-sequence recording in SLiM opens new horizons for forward-time simulation of whole genomes. Mol. Ecol. Resour. 19, 552–566 (2019).

28. R. R. Hudson, M. Slatkin, W. P. Maddison, Estimation of levels of gene flow from DNA sequence data. Genetics. 132, 583–589 (1992).

29. 1001 Genomes Consortium, 1,135 Genomes Reveal the Global Pattern of Polymorphism in Arabidopsis thaliana. Cell. 166, 481–491 (2016).

30. B. J. McGill, B. J. Enquist, E. Weiher, M. Westoby, Rebuilding community ecology from functional traits. Trends Ecol. Evol. 21, 178–185 (2006).

31. P. I. Prado, M. Miranda, A. Chalom, sads: R package for fitting species abundance distributions (Github, 2018; https://github.com/piLaboratory/sads).

32. T. J. Matthews, K. A. Triantis, R. J. Whittaker, F. Guilhaumon, sars: an R package for fitting, evaluating and comparing species–area relationship models. Ecography. 42, 1446–1455 (2019).

33. F. He, S. P. Hubbell, Species-area relationships always overestimate extinction rates from habitat loss. Nature. 473, 368–371 (2011).

34. R. J. H. &. J. van Etten, raster: Geographic analysis and modeling with raster data (2012), (available at http://CRAN.R-project.org/package=raster).

35. M. Exposito-Alonso, 500 Genomes Field Experiment Team, H. A. Burbano, O. Bossdorf, R. Nielsen, D. Weigel, Natural selection in the *Arabidopsis thaliana* genome in present and future climates. Nature. 573, 126–129 (2019).

36. Y. B. Simons, K. Bullaughey, R. R. Hudson, G. Sella, A population genetic interpretation of GWAS findings for human quantitative traits. PLoS Biol. 16, e2002985 (2018).

37. M. Exposito-Alonso, C. Becker, V. J. Schuenemann, E. Reiter, C. Setzer, R. Slovak, B. Brachi, J. Hagmann, D. G. Grimm, J. Chen, W. Busch, J. Bergelson, R. W. Ness, J. Krause, H. A. Burbano, D. Weigel, The rate and potential relevance of new mutations in a colonizing plant lineage. PLoS Genet. 14, e1007155 (2018).

38. H. Seebens, T. M. Blackburn, E. E. Dyer, P. Genovesi, P. E. Hulme, J. M. Jeschke, S. Pagad, P. Pyšek, M. Winter, M. Arianoutsou, S. Bacher, B. Blasius, G. Brundu, C. Capinha, L. Celesti-Grapow, W. Dawson, S. Dullinger, N. Fuentes, H. Jäger, J. Kartesz, M. Kenis, H. Kreft, I. Kühn, B. Lenzner, A. Liebhold, A. Mosena, D. Moser, M. Nishino, D. Pearman, J. Pergl, W. Rabitsch, J. Rojas-Sandoval, A. Roques, S. Rorke, S. Rossinelli, H. E. Roy, R. Scalera, S. Schindler, K. Štajerová, B. Tokarska-Guzik, M. van Kleunen, K. Walker, P. Weigelt, T. Yamanaka, F. Essl, No saturation in the accumulation of alien species worldwide. Nat. Commun. 8, 14435 (2017).

39. H. Seebens, F. Essl, W. Dawson, N. Fuentes, D. Moser, J. Pergl, P. Pyšek, M. van Kleunen, E. Weber, M. Winter, B. Blasius, Global trade will accelerate plant invasions in emerging economies under climate change. Glob. Chang. Biol. 21, 4128–4140 (2015).

40. S. Purcell, B. Neale, K. Todd-Brown, L. Thomas, M. A. R. Ferreira, D. Bender, J. Maller, P. Sklar, P. I. W. de Bakker, M. J. Daly, P. C. Sham, PLINK: a tool set for whole-genome association and population-based linkage analyses. Am. J. Hum. Genet. 81, 559–575 (2007).

41. C. Mérot, R. A. Oomen, A. Tigano, M. Wellenreuther, A Roadmap for Understanding the Evolutionary Significance of Structural Genomic Variation. Trends Ecol. Evol. 35, 561–572 (2020).

42. K. Lucek, Y. Willi, Drivers of linkage disequilibrium across a species’ geographic range. PLoS Genet. 17, e1009477 (2021).

43. J. M. Kreiner, D. A. Giacomini, F. Bemm, B. Waithaka, J. Regalado, C. Lanz, J. Hildebrandt, P. H. Sikkema, P. J. Tranel, D. Weigel, J. R. Stinchcombe, S. I. Wright, Multiple modes of convergent adaptation in the spread of glyphosate-resistant Amaranthus tuberculatus. Proc. Natl. Acad. Sci. U. S. A. 116, 21076–21084 (2019).

44. M. A. Supple, J. G. Bragg, L. M. Broadhurst, A. B. Nicotra, M. Byrne, R. L. Andrew, A. Widdup, N. C. Aitken, J. O. Borevitz, Landscape genomic prediction for restoration of a Eucalyptus foundation species under climate change. Elife. 7 (2018), doi:10.7554/eLife.31835.

45. M. Vallejo-Marín, J. Friedman, A. D. Twyford, O. Lepais, S. M. Ickert-Bond, M. A. Streisfeld, L. Yant, M. van Kleunen, M. C. Rotter, J. R. Puzey, Population genomic and historical analysis suggests a global invasion by bridgehead processes in Mimulus guttatus. Commun Biol. 4, 327 (2021).

46. J. T. Lovell, A. H. MacQueen, S. Mamidi, J. Bonnette, J. Jenkins, J. D. Napier, A. Sreedasyam, A. Healey, A. Session, S. Shu, K. Barry, S. Bonos, L. Boston, C. Daum, S. Deshpande, A. Ewing, P. P. Grabowski, T. Haque, M. Harrison, J. Jiang, D. Kudrna, A. Lipzen, T. H. Pendergast 4th, C. Plott, P. Qi, C. A. Saski, E. V. Shakirov, D. Sims, M. Sharma, R. Sharma, A. Stewart, V. R. Singan, Y. Tang, S. Thibivillier, J. Webber, X. Weng, M. Williams, G. A. Wu, Y. Yoshinaga, M. Zane, L. Zhang, J. Zhang, K. D. Behrman, A. R. Boe, P. A. Fay, F. B. Fritschi, J. D. Jastrow, J. Lloyd-Reilley, J. M. Martínez-Reyna, R. Matamala, R. B. Mitchell, F. M. Rouquette Jr, P. Ronald, M. Saha, C. M. Tobias, M. Udvardi, R. A. Wing, Y. Wu, L. E. Bartley, M. Casler, K. M. Devos, D. B. Lowry, D. S. Rokhsar, J. Grimwood, T. E. Juenger, J. Schmutz, Genomic mechanisms of climate adaptation in polyploid bioenergy switchgrass. Nature (2021), doi:10.1038/s41586-020-03127-1.

47. I. R. MacLachlan, T. K. McDonald, B. M. Lind, L. H. Rieseberg, S. Yeaman, S. N. Aitken, Genome-wide shifts in climate-related variation underpin responses to selective breeding in a widespread conifer. Proc. Natl. Acad. Sci. U. S. A. 118 (2021), doi:10.1073/pnas.2016900118.

48. G. Tuskan, W. Muchero, J.-G. Chen, D. Jacobson, T. Tschaplinski, D. Rokhsar, W. Schackwitz, J. Schmutz, S. DiFazio, Populus Trichocarpa Genome-Wide Association Study (GWAS) Population SNP Dataset Released, (available at https://doi.ccs.ornl.gov/ui/doi/55).

49. Anopheles gambiae 1000 Genomes Consortium, Data analysis group, Partner working group, Sample collections—Angola:, Burkina Faso:, Cameroon:, Gabon:, Guinea:, Guinea-Bissau:, Kenya:, Uganda:, Crosses:, Sequencing and data production, Web application development, Project coordination, Genetic diversity of the African malaria vector Anopheles gambiae. Nature. 552, 96–100 (2017).

50. Z. L. Fuller, V. J. L. Mocellin, L. A. Morris, N. Cantin, J. Shepherd, L. Sarre, J. Peng, Y. Liao, J. Pickrell, P. Andolfatto, M. Matz, L. K. Bay, M. Przeworski, Population genetics of the coral Acropora millepora: Toward genomic prediction of bleaching. Science. 369 (2020), doi:10.1126/science.aba4674.

51. K. Ruegg, E. C. Anderson, M. Somveille, R. A. Bay, M. Whitfield, E. H. Paxton, T. B. Smith, Linking climate niches across seasons to assess population vulnerability in a migratory bird. Glob. Chang. Biol. 27, 3519–3531 (2021).

52. R. A. Bay, R. J. Harrigan, V. Le Underwood, H. Lisle Gibbs, T. B. Smith, K. Ruegg, Genomic signals of selection predict climate-driven population declines in a migratory bird. Science. 359, 83–86 (2018).

53. E. P. Kingsley, K. M. Kozak, S. P. Pfeifer, D.-S. Yang, H. E. Hoekstra, The ultimate and proximate mechanisms driving the evolution of long tails in forest deer mice. Evolution. 71, 261–273 (2017).

54. L. Smeds, J. Aspi, J. Berglund, I. Kojola, K. Tirronen, H. Ellegren, Whole-genome analyses provide no evidence for dog introgression in Fennoscandian wolf populations. Evol. Appl. 14, 721–734 (2021).

55. R. M. Schweizer, J. Robinson, R. Harrigan, P. Silva, M. Galverni, M. Musiani, R. E. Green, J. Novembre, R. K. Wayne, Targeted capture and resequencing of 1040 genes reveal environmentally driven functional variation in grey wolves. Mol. Ecol. 25, 357–379 (2016).

56. 1000 Genomes Project Consortium, A. Auton, L. D. Brooks, R. M. Durbin, E. P. Garrison, H. M. Kang, J. O. Korbel, J. L. Marchini, S. McCarthy, G. A. McVean, G. R. Abecasis, A global reference for human genetic variation. Nature. 526, 68–74 (2015).

57. J. D. Palacio-Mejía, P. P. Grabowski, E. M. Ortiz, G. A. Silva-Arias, T. Haque, D. L. Des Marais, J. Bonnette, D. B. Lowry, T. E. Juenger, Geographic patterns of genomic diversity and structure in the C4 grass Panicum hallii across its natural distribution. AoB Plants. 13, lab002 (2021).

58. A. M. Royer, M. A. Streisfeld, C. I. Smith, Population genomics of divergence within an obligate pollination mutualism: Selection maintains differences between Joshua tree species. Am. J. Bot. 103, 1730–1741 (2016).

59. M. Kapun, J. C. B. Nunez, M. Bogaerts-Márquez, J. Murga-Moreno, M. Paris, J. Outten, M. Coronado-Zamora, C. Tern, O. Rota-Stabelli, M. P. García Guerreiro, S. Casillas, D. J. Orengo, E. Puerma, M. Kankare, L. Ometto, V. Loeschcke, B. S. Onder, J. K. Abbott, S. W. Schaeffer, S. Rajpurohit, E. L. Behrman, M. F. Schou, T. J. S. Merritt, B. P. Lazzaro, A. Glaser-Schmitt, E. Argyridou, F. Staubach, Y. Wang, E. Tauber, S. V. Serga, D. K. Fabian, K. A. Dyer, C. W. Wheat, J. Parsch, S. Grath, M. S. Veselinovic, M. Stamenkovic-Radak, M. Jelic, A. J. Buendía-Ruíz, M. Josefa Gómez-Julián, M. Luisa Espinosa-Jimenez, F. D. Gallardo-Jiménez, A. Patenkovic, K. Eric, M. Tanaskovic, A. Ullastres, L. Guio, M. Merenciano, S. Guirao-Rico, V. Horváth, D. J. Obbard, E. Pasyukova, V. E. Alatortsev, C. P. Vieira, J. Vieira, J. Roberto Torres, I. Kozeretska, O. M. Maistrenko, C. Montchamp-Moreau, D. V. Mukha, H. E. Machado, A. Barbadilla, D. Petrov, P. Schmidt, J. Gonzalez, T. Flatt, A. O. Bergland, Drosophila Evolution over Space and Time (DEST) - A New Population Genomics Resource. bioRxiv (2021), p. 2021.02.01.428994.

60. L. N. Di Santo, S. Hoban, T. L. Parchman, J. W. Wright, J. A. Hamilton, Reduced representation sequencing to understand the evolutionary history of Torrey pine (Pinus torreyana Parry) with implications for rare species conservation. bioRxiv (2021), p. 2021.07.02.450939.

61. J. von Seth, N. Dussex, D. Díez-Del-Molino, T. van der Valk, V. E. Kutschera, M. Kierczak, C. C. Steiner, S. Liu, M. T. P. Gilbert, M.-H. S. Sinding, S. Prost, K. Guschanski, S. K. S. S. Nathan, S. Brace, Y. L. Chan, C. W. Wheat, P. Skoglund, O. A. Ryder, B. Goossens, A. Götherström, L. Dalén, Genomic insights into the conservation status of the world’s last remaining Sumatran rhinoceros populations. Nat. Commun. 12, 2393 (2021).

62. Millennium Ecosystem Assessment, Millennium ecosystem assessment (Millennium Ecosystem Assessment, 2005; http://chapter.ser.org/europe/files/2012/08/Harris.pdf).

63. G. C. Hurtt, L. Chini, R. Sahajpal, S. Frolking, B. L. Bodirsky, K. Calvin, J. C. Doelman, J. Fisk, S. Fujimori, K. Klein Goldewijk, T. Hasegawa, P. Havlik, A. Heinimann, F. Humpenöder, J. Jungclaus, J. O. Kaplan, J. Kennedy, T. Krisztin, D. Lawrence, P. Lawrence, L. Ma, O. Mertz, J. Pongratz, A. Popp, B. Poulter, K. Riahi, E. Shevliakova, E. Stehfest, P. Thornton, F. N. Tubiello, D. P. van Vuuren, X. Zhang, Harmonization of global land use change and management for the period 850–2100 (LUH2) for CMIP6. Geosci. Model Dev. 13, 5425–5464 (2020).

64. S. Theodoridis, C. Rahbek, D. Nogues-Bravo, Exposure of mammal genetic diversity to mid-21st century global change. Ecography (Cop.) (2021), doi:10.1111/ecog.05588.

65. I. Overcast, M. Ruffley, J. Rosindell, L. Harmon, P. A. V. Borges, B. C. Emerson, R. S. Etienne, R. Gillespie, H. Krehenwinkel, D. Luke Mahler, F. Massol, C. E. Parent, J. Patiño, B. Peter, B. Week, C. Wagner, M. J. Hickerson, A. Rominger, A unified model of species abundance, genetic diversity, and functional diversity reveals the mechanisms structuring ecological communities. bioRxiv (2020), p. 2020.01.30.927236.

66. S. Díaz, N. Zafra-Calvo, A. Purvis, P. H. Verburg, D. Obura, P. Leadley, R. Chaplin-Kramer, L. De Meester, E. Dulloo, B. Martín-López, M. R. Shaw, P. Visconti, W. Broadgate, M. W. Bruford, N. D. Burgess, J. Cavender-Bares, F. DeClerck, J. M. Fernández-Palacios, L. A. Garibaldi, S. L. L. Hill, F. Isbell, C. K. Khoury, C. B. Krug, J. Liu, M. Maron, P. J. K. McGowan, H. M. Pereira, V. Reyes-García, J. Rocha, C. Rondinini, L. Shannon, Y.-J. Shin, P. V. R. Snelgrove, E. M. Spehn, B. Strassburg, S. M. Subramanian, J. J. Tewksbury, J. E. M. Watson, A. E. Zanne, Set ambitious goals for biodiversity and sustainability. Science. 370, 411–413 (2020).

67. C. Rahbek, R. K. Colwell, Biodiversity: Species loss revisited. Nature. 473 (2011), pp. 288–289.

68. H. M. Pereira, G. C. Daily, Modeling biodiversity dynamics in countryside landscapes. Ecology. 87, 1877–1885 (2006).

69. J. Harte, A. B. Smith, D. Storch, Biodiversity scales from plots to biomes with a universal species-area curve. Ecol. Lett. 12, 789–797 (2009).

70. D. R. Lockwood, C. M. Richards, G. M. Volk, Probabilistic models for collecting genetic diversity: Comparisons, caveats, and limitations. Crop Sci. 47, 861–866 (2007).

71. L. Czech, M. Exposito-Alonso, grenepipe: A flexible, scalable, and reproducible pipeline to automate variant and frequency calling from sequence reads. arXiv (2021), (available at http://arxiv.org/abs/2103.15167).

72. A. Miraldo, S. Li, M. K. Borregaard, A. Flórez-Rodríguez, S. Gopalakrishnan, M. Rizvanovic, Z. Wang, C. Rahbek, K. A. Marske, D. Nogués-Bravo, An Anthropocene map of genetic diversity. Science. 353, 1532–1535 (2016).

73. D. H. Parks, T. Mankowski, S. Zangooei, M. S. Porter, D. G. Armanini, D. J. Baird, M. G. I. Langille, R. G. Beiko, GenGIS 2: Geospatial Analysis of Traditional and Genetic Biodiversity, with New Gradient Algorithms and an Extensible Plugin Framework. PLoS One. 8, e69885 (2013).

74. C. Li, D. Tian, B. Tang, X. Liu, X. Teng, W. Zhao, Z. Zhang, S. Song, Genome Variation Map: a worldwide collection of genome variations across multiple species. Nucleic Acids Res. 49, D1186–D1191 (2021).

75. M. Kardos, E. Armstrong, S. W. Fitzpatrick, S. Hauser, P. Hedrick, J. Miller, D. Tallmon, W. Chris Funk, The crucial role of genome-wide genetic variation in conservation. bioRxiv (2021), p. 2021.07.05.451163.

76. D. Spielman, B. W. Brook, R. Frankham, Most species are not driven to extinction before genetic factors impact them. Proceedings of the National Academy of Sciences. 101 (2004), pp. 15261–15264.

77. Ipbes, Global assessment report of the Intergovernmental Science-Policy Platform on Biodiversity and Ecosystem Services (IPBES Secretariat, Bonn, Germany, 2019; https://www.ipbes.net/news/ipbes-global-assessment-summary-policymakers-pdf).

